# Decoding Plasticity Regulators and Transition Trajectories in Glioblastoma with Single-cell Multiomics

**DOI:** 10.1101/2025.05.13.653733

**Authors:** Manu Saraswat, Laura Rueda-Gensini, Elisa Heinzelmann, Tannia Gracia, Fani Memi, Grant de Jong, Jannes Straub, Cedar Schloo, Dirk C. Hoffmann, Erik Jung, Tim Kindinger, Bettina Weigel, Bryce Lim, Sophie Weil, Oliver Gould, Richard Mair, Katharina Mikulik, Martin Rohbeck, Wolfgang Wick, Frank Winkler, Omer Ali Bayraktar, Oliver Stegle, Moritz Mall

## Abstract

Glioblastoma (GB) is one of the most lethal human cancers, marked by profound intratumoral heterogeneity and near-universal treatment resistance. Cellular plasticity, the capacity of cancer cells to transition between phenotypic states, drives GB progression and resistance. However, the regulatory logic that permits or restricts specific state transitions remains poorly understood. Here, we integrated single-nucleus RNA and chromatin accessibility multi-ome profiles from over one million cells across primary IDH-wildtype GBs and developed scDORI, a scalable deep-learning framework to infer enhancer-driven gene regulatory networks (eGRNs) at single-cell resolution. Our analysis revealed a structured hierarchy of GB cell states governed by distinct regulatory programs, with marked variability in epigenetic plasticity that enables or constrains transitions. Neuronal-like tumor cells emerge as a low plasticity state that deploys active repression, in contrast to more permissive progenitor-like and astrocytic states. We identified the neuronal-like state-specific repressor MYT1L as a key regulator that silences master transcription factors of alternative states. MYT1L gain-of-function in patient-derived GB cells reduced chromatin accessibility, induced neuronal-like identity, and restricted proliferation and invasion *in vivo*, whereas loss-of-function reactivated plasticity and accelerated malignant features. Our findings delineate the epigenetic architecture and associated transcriptional master regulators that shape GB state trajectories, and establish safeguard repressors such as MYT1L as potential therapeutic targets to constrain malignant plasticity.

## Main

The capacity of cancer cells to switch phenotypic states, also called cellular plasticity, is an emerging hallmark of cancer involved in tumor initiation and progression, including therapeutic resistance, tumor relapse, and metastasis ^1^. Glioblastoma (GB), the most common and lethal primary brain tumor in adults, is characterized by profound cellular heterogeneity and dismal prognosis ^2,3^. Although extensive sequencing efforts have catalogued its genomic landscape, actionable genetic drivers remain scarce, and clinical responses to targeted therapies have been minimal ^4–6^. Longitudinal studies have underscored the limited contribution of therapy-induced genetic evolution in GB, revealing remarkable clonal stability despite treatment ^7,8^. These findings suggest that non-genetic mechanisms, such as epigenetic plasticity and transcriptional reprogramming, may be central to the adaptive potential and treatment resistance in GB.

Seminal single-cell transcriptomic studies have shown that malignant GB cells occupy a continuum of cellular states resembling developmental progenitors (e.g., OPC-like and NPC-like), as well as astrocytic and mesenchymal-like phenotypes ^9,10^. Patient-derived xenograft models have demonstrated that individual GB cells can give rise to multiple states, implying substantial plasticity ^10,11^. However, whether these state transitions follow stereotyped routes or variable trajectories and how they are promoted or constrained by gene regulatory mechanisms remains poorly understood.

Here, we address these questions by integrating multi-ome single-nucleus RNA and chromatin accessibility (snRNA-ATAC) profiles from over one million cells across spatially mapped regions of primary IDH-wildtype GBs generated in a companion study (de Jong *et al*., 2025)^66^. Through a novel computational approach (scDORI), we decoded enhancer-driven gene regulatory networks (eGRNs) at single-cell resolution and mapped the plasticity landscape across GB cell states. Without relying on predefined annotations, scDORI identified joint transcriptomic and epigenetic regulatory modules that capture the gene regulatory logic underlying cell identity and state transitions in GB.

Our analysis revealed that cancer cell states are arranged in a clear hierarchy, demarcated by epigenetic “barriers” and “highways” that guide transitions by hijacking neurodevelopmental transcription factors. Neuronal-like cells, in particular, showed strikingly low plasticity and elevated transcriptional repression, acting as a constrained state in the transition space of GB cells. This repression is orchestrated by state-specific factors such as MYT1L, which actively silence transcription factors governing alternative fates. Experimental perturbation of MYT1L in patient-derived GB cells confirmed its capacity to restrict plasticity, induce neuronal-like morphology, and reduce tumor growth and mortality upon transplantation in mice. These results suggest that manipulating repressors of plasticity may lock GB cells into more benign, less adaptive states and might open new avenues for treatment.

Together with our companion study mapping GB state transitions and spatial architectures, these findings reveal a universal, epigenetically controlled hierarchy of cell states and transition routes in GB. Our work establishes transcriptional repressors as key regulators of plasticity and underscores the therapeutic potential of targeting the epigenetic foundations of state switching in brain cancer.

## Results

### Glioblastoma Cells Exhibit State-Specific Levels of Epigenetic Plasticity

To characterize the regulatory landscape of glioblastoma, we mined a single-nuclei multi-ome atlas (snRNA-ATAC-seq) generated in a companion study^66^, spanning parallel gene expression and chromatin accessibility profiles of over 1 million cells collected across multiple sites (4-15) of 12 isocitrate dehydrogenase wildtype (IDH-WT) primary GB tumors, including peritumoral regions (**Fig. 1A**). Malignant GB cells exhibited vastly heterogeneous transcriptional profiles, ranging from developmental-like (OPC-NPC-like, OPC-like, OPC-neuronal-like, NPC-neuronal-like, AC-progenitor-like) to glial injury response and hypoxia-associated (AC-gliosis-like, Gliosis-like, Hypoxia) phenotypes, as well as cycling signatures (Proliferative)^66^. Expanding cell numbers from previous datasets by more than tenfold ^10,12–15^, this atlas provides the foundation for dissecting the regulatory mechanisms driving GB heterogeneity at an unprecedented scale.

**Figure 1.**
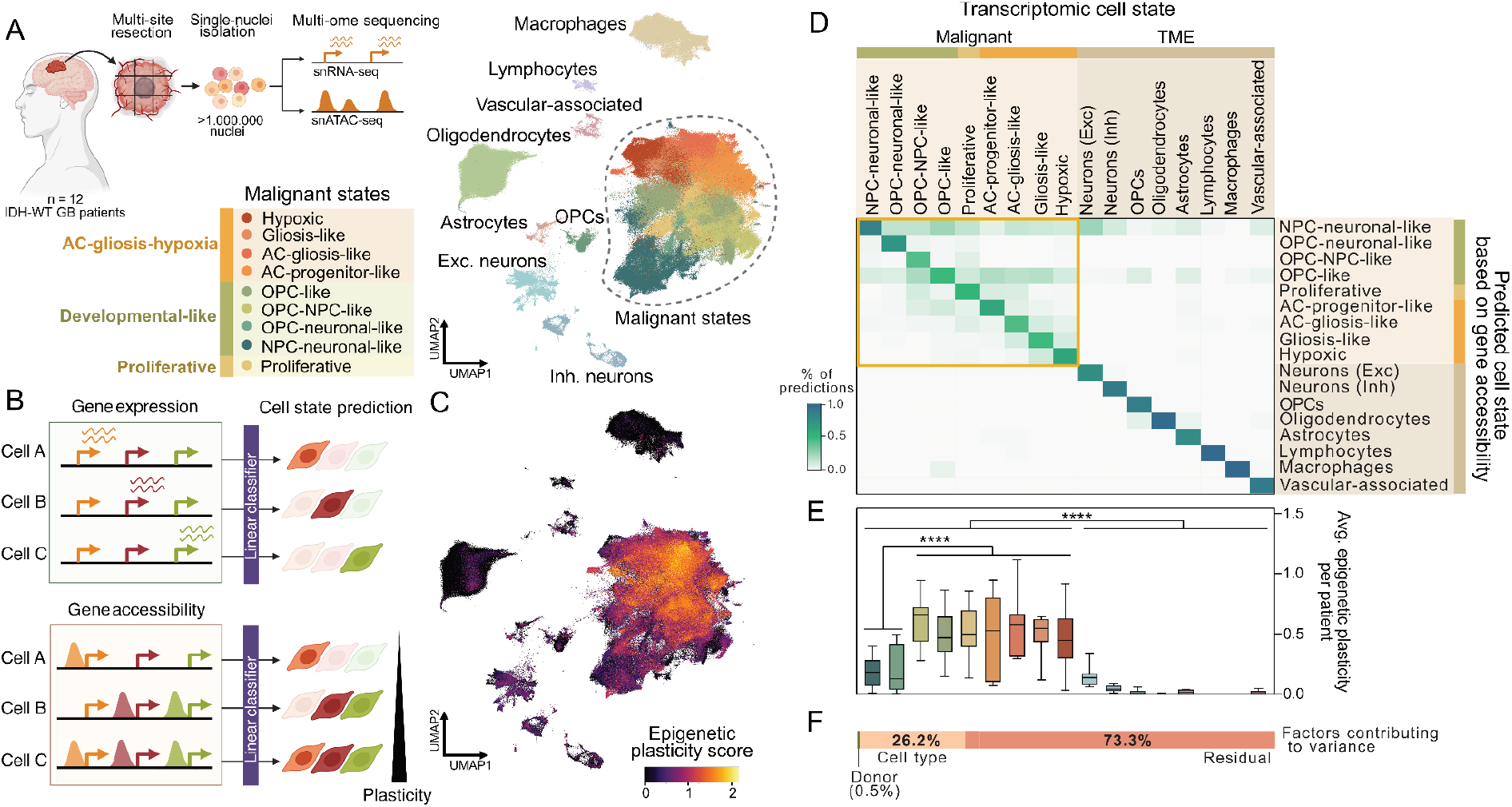
Heterogeneity of Epigenetic Plasticity in Glioblastoma Revealed by Multi-ome Analysis. (A) Left: Experimental setup. Right: UMAP projection and annotation of the RNA component of 368,054 TME and 657,275 GB cells from 12 patients processed for snRNA-ATAC multi-ome analysis. Colors and labels denote states of indicated malignant GB and TME cells. (B) Workflow schematic to assess epigenetic plasticity of cells. We compare the ability of a linear classifier to predict cell states based on gene expression and gene accessibility. Cells with uncertain or ambiguous predictions based on gene accessibility exhibit higher epigenetic plasticity (Methods). (C) Epigenetic plasticity score determined as in B, reflecting uncertainty of cell state predictions based on gene accessibility, visualised for all cells on UMAP in A. (D) Confusion matrix showing percentage of cells from a given expression-defined state (columns) assigned to different cell states (rows) based on gene accessibility profiles. Off-diagonal values correspond to incorrect predictions. Malignant cell types (yellow box) exhibited lower prediction accuracy, indicating elevated epigenetic plasticity. (E) Average epigenetic plasticity score for indicated cell states across patients. Malignant states exhibit varying degrees of epigenetic plasticity, with NPC-neuronal-like and OPC-neuronal-like cells displaying the lowest levels (p=6.7×10^−6^, two-sided t-test). (F) Variance component analysis of epigenetic plasticity scores across cells, assessing the contribution of patient, cell type, and unmodelled residual factors.

Drawing on the multi-omic dimension of our dataset, we explored the epigenetic plasticity of GB cells, following the assumption that accessible gene regions without active transcription indicate epigenetic plasticity. Inspired by Burdziak et al.^16^, we quantified how accessibility profiles of cell state markers can predict transcriptional cell states (**Fig. 1B, Extended Data Fig. 1A-B, Supplementary Table 1**; Methods). In this model, cells with uncertain predictions based on chromatin profiles are identified as more epigenetically plastic. Our analysis revealed that malignant cells display greater epigenetic plasticity compared to non-malignant tumor microenvironment (TME) cells (**Fig. 1C**; p=9.8×10^−14^, two-sided t-test), with chromatin accessibility in malignant cells often being compatible with multiple transcriptional states (**Fig. 1D**). Yet, the extent of this plasticity significantly varied across GB cell states (**Fig. 1E**), with NPC-neuronal-like and OPC-neuronal-like cells standing out as least plastic states (p=6.7×10^−6^, two-sided t-test). These variations in epigenetic plasticity could not be explained by differences in global coupling between gene expression and accessibility, were robust to differences in cell numbers per state, and were consistently observed across patients (**Fig. 1F and Extended Data Fig. 1C-F**). This suggests specific regulatory mechanisms underlie GB plasticity and govern cell state heterogeneity.

### Decoding the Gene Regulatory Architecture of Glioblastoma Heterogeneity

To determine specific gene regulatory processes regulating the observed epigenetic plasticity, we aimed to identify relevant TFs and *cis*-regulatory elements (CRE), as well as their interplay across GB states, using enhancer-driven gene regulatory networks (eGRNs). Established eGRN inference methods ^17–27^ face limitations that hamper their applicability to a highly heterogeneous single-cell tumor atlas like ours. Specifically, existing methods offer poor scalability to large datasets^28^, and their reliance on pre-defined cell type annotations means they cannot model continuous and cell state-specific differences of eGRNs. Moreover, previous methods primarily focus on signatures of transcriptional activation, despite the growing recognition of gene repression as a key regulator of cell identity and plasticity in cancer ^29,30^.

To address these limitations, we developed single-cell Deep multi-Omic Regulatory Inference (scDORI), a computational framework for inferring continuous eGRNs and their relationships across cell populations using scRNA-ATAC multi-ome data (**Fig. 2A**; Methods). Building on autoencoders ^31^, scDORI combines principles of multi-omics dimensionality reduction and Topic modeling to decompose gene expression and chromatin accessibility profiles into gene regulatory modules ^17,32–37^, hereafter referred to as “Topics”. Unlike conventional dimensionality reduction approaches that rely solely on correlative signatures, scDORI Topics adhere to regulatory constraints that govern the relationship between TFs, CREs, and target genes. This is achieved by employing multiple coupled decoders that capture established signatures of eGRNs, including peak co-accessibility, TF chromatin binding, CRE-gene linkages, and expression correlations between TFs and their target genes that support activation or repression interactions ^17–25,38^ (**Extended Data Fig. 2A**; Methods). As a result, individual scDORI Topics represent TF-to-target gene relationships mediated by specific CREs (eGRN) (**Fig. 2A**, bottom). Because the multi-omic profile of each cell is modelled as a mixture of Topics, scDORI can capture fine-grained variation in eGRNs across cell states without relying on predefined annotations (**Fig. 2A**, right). Conversely, a single Topic can be active across multiple cell states, enabling the identification of shared aspects of gene regulation. scDORI scales efficiently to datasets comprising millions of cells (**Extended Data Fig. 2B)**, and performs favorably compared to existing eGRN inference methods on benchmarks such as prediction of enhancer-promoter links (**Extended Data Fig. 2C**; Methods).

**Figure 2.**
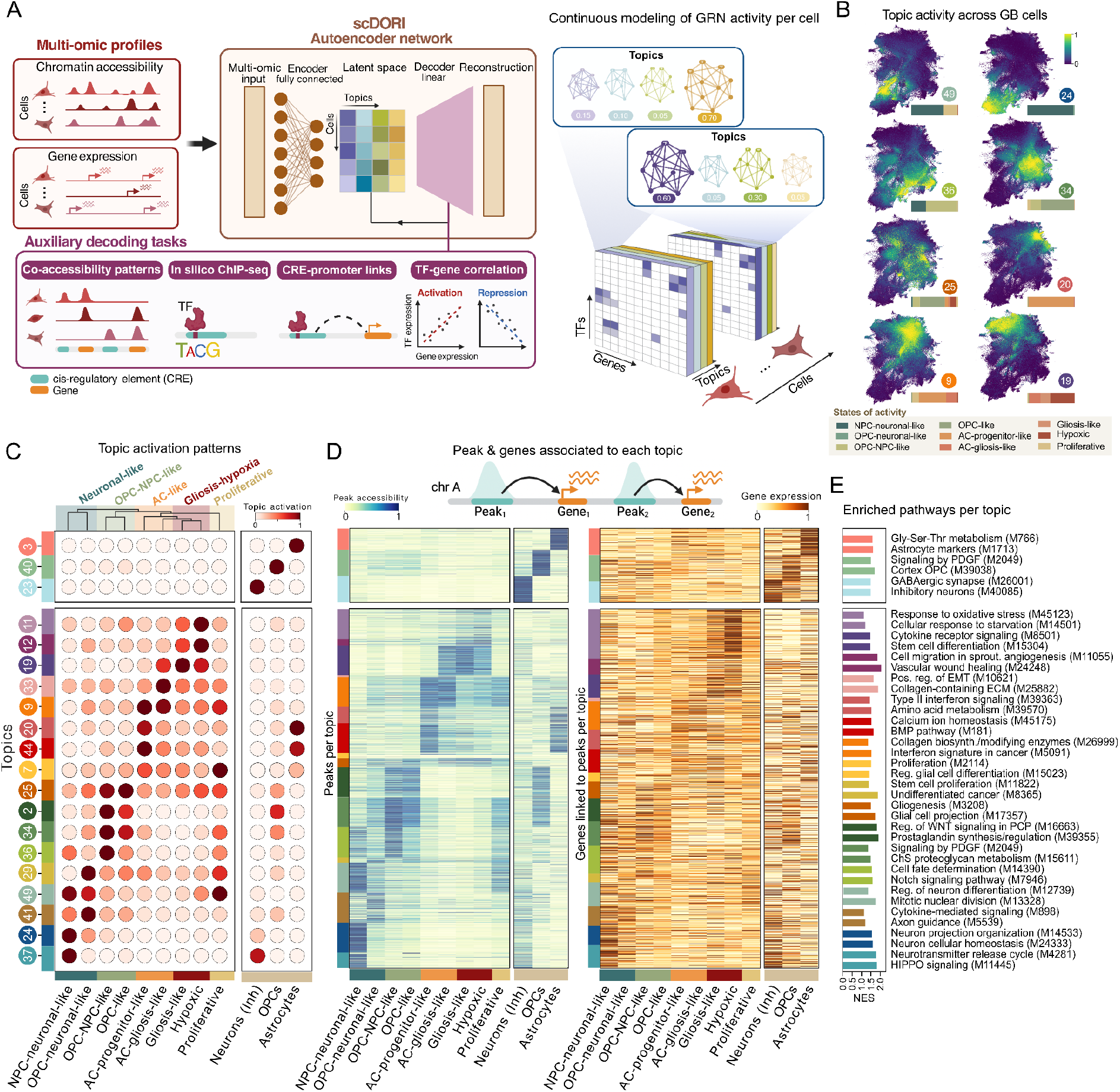
Epigenetic and Transcriptional Gene Regulatory Programs of GB. (A) Outline of the scDORI model architecture. Top: scATAC-RNA multi-ome data is used as autoencoder input to infer continuous eGRNs referred to as Topics. scDORI computes Topics via mechanistically constrained decoding tasks, modelling individual cells as a linear combination of co-active Topics. Bottom: Mechanistic constraints in scDORI, implemented as auxiliary decoding tasks incorporating information on co-accessible peak, TF binding, CRE-gene linkages, and TF-gene expression correlations. (B) Activity of selected Topics in malignant cells visualised on the UMAP representation of **Fig. 1A**. (C) Activation patterns of selected Topics across GB states, showing that GB cell states are associated with multiple Topics. Top: Hierarchical clustering of GB states based on Topic activation patterns, revealing five distinct regulatory states. Color denotes average Topic activation scaled across states. All Topic activation plots can be found in **Extended Data Fig. S3A and Supplementary Table 2**. (D) Accessibility of Topic-associated ATAC peaks (blue) and expression of predicted *cis*-linked genes (orange) across cell states. Individual rows correspond to peaks (left) and *cis*-linked genes (right), grouped by the Topic with the highest association score for each peak. (E) Enriched pathways in scDORI Topics, based on gene set enrichment analysis (GSEA) of Topic genes ranked by Topic-association score. Shown are the normalized enrichment scores (NES) of two representative pathways significantly enriched in each Topic (p-adj<0.05). For complete results, see **Supplementary Table 2**.

Applied to our GB atlas, scDORI identified 31 Topics with distinct activity patterns across cells and patients (**Fig. 2B-C, Extended Data Fig. 3, and Supplementary Table 2**), each capturing co-accessible chromatin regions *cis*-linked to co-expressed genes (**Fig. 2D and Extended Data Fig. 4A**). Notably, despite their mechanistic constraints, scDORI Topics explained the transcriptional variation within tumor cells to a similar or greater degree than conventional dimensionality reduction methods ^25,34^ (**Extended Data Fig. 4B-C**). Clustering of transcriptional GB states based on Topic activity patterns revealed four major regulatory states alongside a distinct Proliferative (cycling) population, stratifying developmental states (Neuronal-like and OPC-NPC-like), and AC-gliosis-hypoxia states (AC-like and Gliosis-hypoxia) (**Fig. 2C**). Topics specific to individual states reflected gene programs characteristic of their developmental counterparts, such as, neurotransmitter release in Neuronal-like cells (Topic 37) ^39^, PDGF signaling in OPC-NPC-like cells (Topic 34) ^40^, and calcium ion homeostasis in AC-like cells (Topic 44) ^41^ (**Fig. 2E, Extended Data Fig. 3A, and Supplementary Table 2**). These programs were also active in corresponding TME cell types, consistent with tumor cells co-opting developmental pathways.

Beyond GB state-level distinctions, scDORI resolved functional heterogeneity within individual malignant states. For instance, in Gliosis-Hypoxia cells, distinct Topics captured hallmark features such as hypoxic stress (Topics 11) ^42^, cytokine signaling (Topic 19) ^43^, and vascular wound healing (Topic 12) ^44^(**Fig. 2E**). scDORI also identified “inter-state*”* Topics (Topic 7, 25, 29, 33, 36, 49) that span multiple GB states, which were enriched for programs linked to fate commitment and differentiation, indicating their potential role in mediating state transitions (**Fig. 2C,E and Extended Data Fig. 3A-B**). Notably, these Topics were also active in cycling cells, implicating proliferation as a hub for state transitions (**Fig. 2C**). Finally, scDORI identified Topics that were broadly active across the majority of tumor cells, reflecting core features of GB biology such as postsynaptic signaling (Topic 39) ^45^, inflammatory response (Topic 31) ^46^, and cell adhesion (Topic 18) (**Extended Data Fig. 3A-B**). We tested the robustness of scDORI Topics across multiple model restarts and by confirming consistent mapping across GB patients in our dataset and within an independent single-cell GB atlas ^10^ (**Extended Data Fig. 3C and Extended Data Fig. 5A-C**). Moreover, scDORI-derived Topic-specific eGRNs were predictive of differentially expressed genes between GB cells, performing favorably compared to existing approaches ^25,26^ (**Extended Data Fig. 5D-E**; Methods). Collectively, scDORI Topics redefine GB heterogeneity from a gene regulation standpoint, building the foundation to gain universal insights into how GB cell identity is regulated, shifted, and maintained.

### Mapping the Regulatory Drivers of GB Cell State Transitions

One of the key open questions in GB biology is whether cancer cells transition according to defined rules, and if so, which TFs govern these transitions ^9,10^. Building on the hypothesis that the activation of master regulators can initiate cell state transitions, we set out to identify key regulator TFs for each Topic. Using Topic-specific eGRNs from scDORI, we defined “Topic Regulators” (TRs) as TFs with the strongest downstream regulatory influence within each Topic (**Fig. 3A and Supplementary Table 2**; Methods). On average, scDORI identified ∼9 TRs per Topic. These included known GB cell state regulators, such as ASCL1 in OPC-NPC-like cells (Topic 36), SOX11 in neuronal-like cells (Topic 24), and FOSL2 in Gliosis-Hypoxia cells (Topic 19), as well as regulators associated with respective Topic programs such as hypoxia (BACH1, Topic 19), and stress response (ATF3, JUND, Topic 11)^10,47,48^. TRs of state-specific Topics included lineage-specific TFs, such as SOX6 (Topic 34, OPCs) ^49^, MYT1L (Topic 24, neurons) ^50^, and TRPS1 (Topic 9, astrocytes) ^51^, further supporting that GB cells reutilise developmental TFs and gene programs ^52^. Our analysis also revealed TRs not previously associated with GB (POU2F2, TCF7L2, HNF4G, TFCP2L1), providing candidates for further exploration (**Fig. 3A and Supplementary Table 2**).

**Figure 3.**
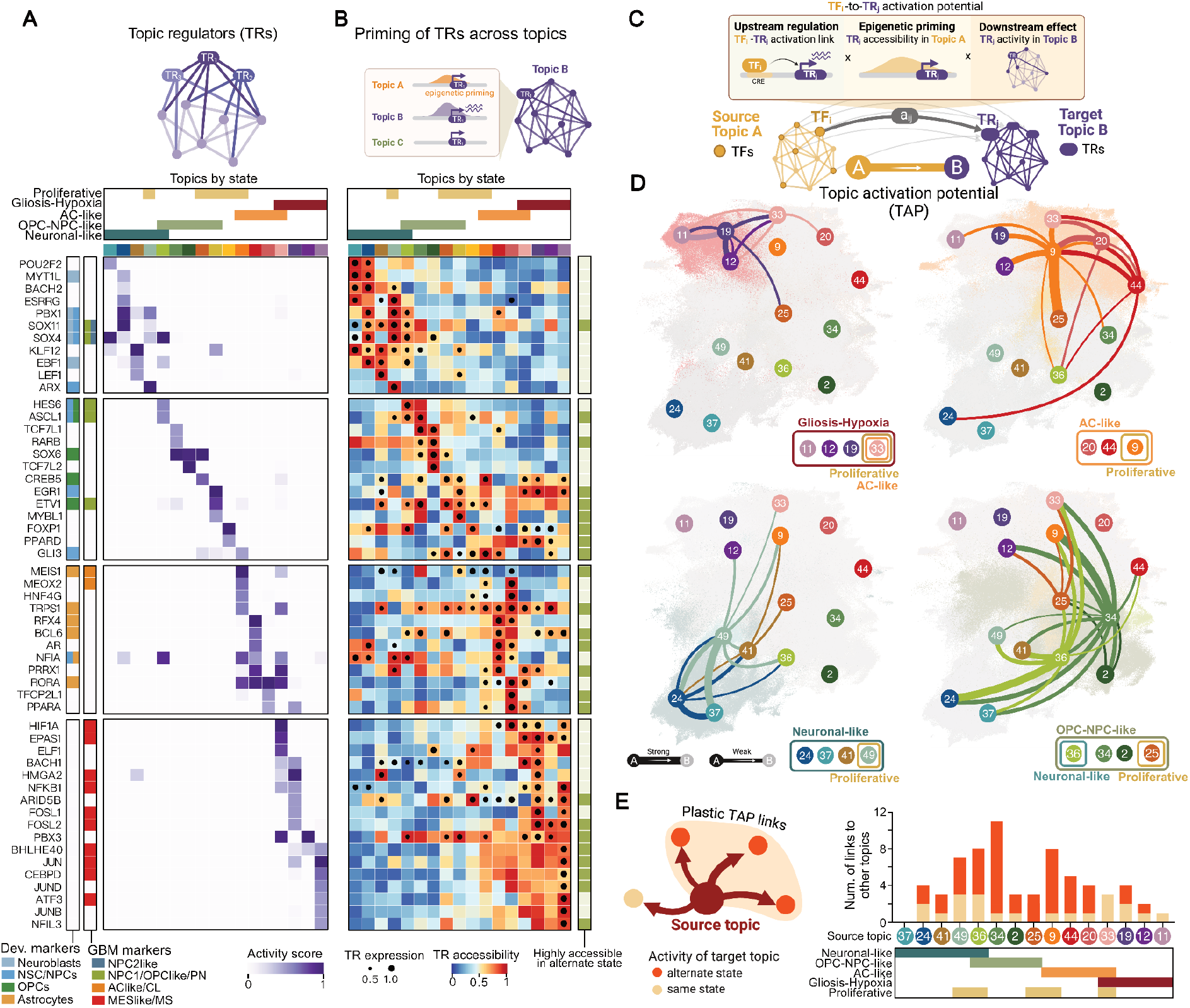
Cell-state Specific eGRNs Inform Transition Potentials of GB cell states. (A) Topic regulators (TRs) across GB cells, defined as the TFs with the highest cumulative activation of genes in each Topic-specific GRN (Methods). Shown are cumulative activation scores for selected TRs and Topics. The bar on the left indicates TFs previously reported as GB cell state markers or as highly expressed in developmental cell types (Neuroblasts, NSC/NPCs, OPCs, astrocytes). For a complete list of TRs per Topic, see **Supplementary Table 2**. (B) Mean chromatin accessibility (color intensity) and mean expression (dot size) of TRs from (A) across selected Topics. The bar on the right highlights highly accessible TRs (>0.75 scaled accessibility) in Topics of an alternate state, suggesting potential priming for transition. (C) The Topic Activation Potential (TAP) from source Topic A to target Topic B (TAP_a_,_β_) integrates: (i) predicted activation links from TFs in Topic A towards TRs of Topic B, (ii) epigenetic priming of Topic B TRs in Topic A cells, and (iii) the downstream activity scores of these TRs in Topic B (Methods). (D) TAP links between selected malignant Topics determined using the approach in C, visualised on UMAP representation of malignant cells in **Fig. 1A**. Nodes correspond to Topics; edge color indicates source Topic, and edge width TAP values. TAP values for all pairs of malignant Topics can be found in **Extended Data Fig. 6C** and **Supplementary Table 3**. (E) Number of TAP links from selected source Topics towards Topics active in the same state (yellow) or alternate states (orange). Neuronal-like Topics display limited potential to activate alternate-state Topics, contrasting with OPC-NPC-like and AC-like Topics. TAP links for all pairs of malignant Topics can be found in **Supplementary Table 3**.

Having identified TRs across GB states, we investigated whether TRs associated with one Topic are either expressed or epigenetically accessible in Topics of a different state, potentially explaining transitions between cell states. While several TRs exhibited expression in at least one other Topic (37/87, 43%), only 16% (14/87) were expressed in a Topic that marks an alternate GB cell state (**Fig. 3B, Extended Data Fig. S6A, and Supplementary Table 2**). In contrast, TR accessibility was more widespread, with 54% (47/87) being highly accessible in Topics of an alternate GB state, suggesting they are epigenetically primed for activation (**Fig. 3B and Extended Data Fig. S6B**).

Building on these observations, we developed a framework to systematically quantify the activation potential between cell states, by estimating how readily the TRs of a target Topic can be activated within a different source Topic. This metric, which we term Topic Activation Potential (TAP), integrates three components: (i) the presence of regulatory interactions from TFs in the source Topic to TRs of the target Topic; (ii) the accessibility of target TRs in the source Topic, reflecting epigenetic priming; and (iii) the relative importance of each TR within its Topic (**Fig. 3C and Supplementary Table 3**; Methods). All components contributed uniquely to the metric and jointly revealed distinct transition trajectories among GB states (**Fig. 3D and Extended Data Fig. 6C-E**), each characterized by different driver TFs (**Extended Data Fig. 6F**). While inter-state Topics (33, 36, 49) emerged as connectivity hubs across states, consistent with their enrichment for fate change-related signatures (**Fig. 2E**), striking differences were observed when examining the core GB states. In particular, OPC–NPC-like (34, 25) and AC-like (9, 44, 20) Topics exhibited strong TAP connections to multiple alternate states, whereas transitions from Neuronal-like (24, 37, 41) and Gliosis–Hypoxia (19, 12, 11) Topics were largely confined within their respective states (**Fig. 3D-E**). Together, these results highlight marked, state-specific differences in transition potential, likely shaped by distinct epigenetic landscapes and regulatory architectures that govern plasticity across GB states.

### Activation and Repression Jointly Govern GB Cell State Transitions

Having identified marked differences in transition potentials between GB states based on activation, we next asked whether the absence of activation is reinforced by active repression. To that end, we defined a Topic Repression Score (TRS), conceptually analogous to TAP but considering signatures of repression instead of activation (**Fig. 4A and Supplementary Table 3**; Methods). Remarkably, Neuronal-like Topics (24, 37) showed the most widespread repression over Topics of alternate GB states (**Fig. 4B**), driven by the aggregated effects of multiple Neuronal-like TFs (**Extended Data Fig. 7A and Supplementary Table 3)**. In contrast, other Topics showed repression restricted to specific states (e.g. Topic 2, 20, 36), or minimal overall repression (Topic 9, 44) (**Fig. 4B**). Similar trends, though less pronounced, were observed when assessing repression of Topic-specific genes and state marker genes (**Extended Data Fig. 7B-C**; Methods). Notably, total repression exerted by each Topic negatively correlated with the epigenetic plasticity of its associated cells (**Fig. 1E and 4B**), suggesting that repression is a barrier to epigenetic plasticity and state transitions.

**Figure 4.**
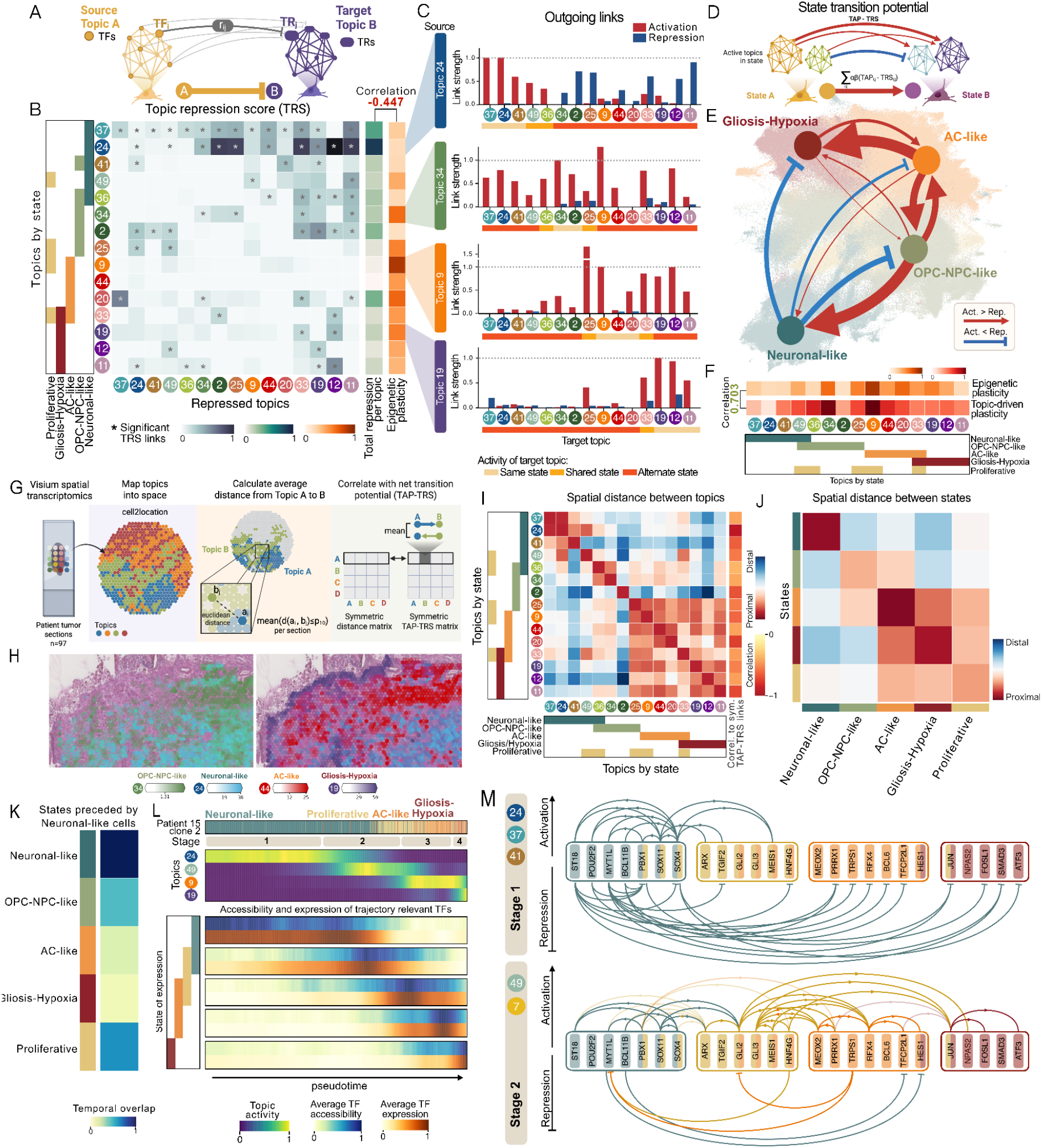
Activation and Repression Dictate Hierarchical State Transitions That Follow GB Spatio-temporal Logic. (A) The Topic repression score (TRS) from source Topic A to target Topic B integrates: (i) predicted repression links from TFs in Topic A towards TRs of Topic B, (ii) epigenetic barriers of Topic B TRs in Topic A cells, and (iii) the downstream activity scores of these TRs in Topic B (Methods). (B) TRS values between indicated pairs of Topics highlight that Neuronal-like Topics exert the highest repressive effects towards alternate state Topics. Asterisks label significant TRS links. The bar on the right shows the aggregated repression of each Topic towards alternate Topics, which inversely correlates with the epigenetic plasticity of the same Topic (derived as in **Fig. 1C**; Methods). (C) Activating (TAP) and repressive (TRS) interactions from selected state-specific Topics towards indicated target Topics. Neuronal-like Topic 24 showed the highest degree of repression, while Topics 9, 34, and 19 mainly displayed activation and little repression towards other Topics (Methods). (D) State transition potential calculation, which considers the net effect of activating (TAP) and repressive (TRS) interactions between Topics active in the source state towards those in the target state. ɑ and β denote the proportional activation of Topics in source and target states, respectively. (Methods) (E) GB state transition networks derived as described in (C), visualised over UMAP of malignant cells from **Fig. 1A**. Unlike other states, Neuronal-like cells exhibited dominant repressive interactions against alternate states, considerably limiting state transitions. (F) Topic-driven plasticity, defined as the cumulative effect of activating (TAP) and repressive (TRS) interactions towards all Topics active in an alternate state. Topic plasticity patterns positively correlated with the global epigenetic plasticity score of associated cell populations, derived as in **Fig. 1C**. (G) Schematic of spatial analysis to estimate transition probabilities based on Topic spatial organization on matching tumor sections using Visium spatial transcriptomics. (H) Representative Visium sections of GB tumors depicting the spatial distribution of dominant Topics from each GB state. (I) Average spatial distances between cell populations with different Topic activations in GB sections (N=97). OPC-NPC-like and AC-like Topics are spatially closer on average compared to Neuronal-like Topics. The bar on the right shows the correlation between the spatial distances and the net regulatory interactions (TAP-TRS) between Topics, indicating that Topic pairs with high net activation potential tend to be spatially proximal. (J) Average spatial distance between GB state-associated Topics across all sections (N=97), computed by aggregating the distance between state-specific Topics in (H), weighted by the proportional activity of each Topic within a respective state (Methods). Neuronal-like cells are the most spatially segregated in GB tumors. (K) GB states temporally preceded by Neuronal-like cells within pseudotime-aligned clonal cell populations (N=60) (Methods). Values reflect temporal co-occurrence between the Topics of each state. (L) Pseudotemporal ordering of a clonal cell population transitioning from Neuronal-like towards AC-like/Gliosis-Hypoxia states via Proliferative state. Sequential Topic activation patterns define distinct transition stages 1-4 (see **Extended Data Fig. 9A**). Matched chromatin accessibility and expression of trajectory-relevant TFs are shown across pseudotime. Transitions out of Neuronal-like state are accompanied by marked chromatin accessibility rearrangements. (M) Predicted gene regulatory interactions of transcription activators and repressors along the trajectory in (L), inferred based on active Topics at each stage. Links going opposite to the current trajectory are shaded for clarity. Stage 1 shows dominant repressive links targeting TFs of alternate states, followed by their shutdown through active repression as the trajectory progresses. See **Extended Data Fig. 9B** for regulatory links in later stages of the trajectory.

To assess how activation and repression jointly shape and balance transitions, we defined a net transition score of TAP minus TRS (**Fig. 4C and Extended Data Fig. 7D**). Aggregation of this score at the level of GB states revealed a hierarchy of cell state transition potentials and constraints (**Fig. 4D-E, Extended Data Fig. 7E, and Supplementary Table 3**), which is consistent with the epigenetic plasticity of each state (**Fig. 1E and Fig. 4F**). At the top of this hierarchy, OPC–NPC-like and AC-like states exhibited strong overall activating links to other states, reflecting permissive epigenetic configurations conducive to transition. In contrast, Neuronal-like and Gliosis–Hypoxia cells showed limited activation towards alternate states, with Neuronal-like cells further restricting their plasticity with repressive barriers (**Extended Data Fig. 7E**). In general, activation and repression potentials were often asymmetric, i.e. OPC-NPC-like to Neuronal-like and AC-like to Gliosis–Hypoxia transitions were much more likely than their opposite counterparts, highlighting preferential transition directions between states (**Fig. 4E and Extended Data Fig. 7E**). Together, these observations reconstructed a cellular trajectory in GB, offering critical insights into the regulatory logic underlying GB progression.

To assess to what extent the modelled transition hierarchy can explain tumor architecture, we analyzed the spatial distribution of Topics within matched tumor sections using Visium spatial transcriptomics. This approach is based on the premise that cell states with high transition potential are more likely to occur in close spatial proximity, whereas those separated by regulatory barriers are expected to be spatially segregated (**Fig. 4G-H, Supplementary Table 4**)^66^. Indeed, Topic pairs with high net transition values (TAP-TRS), such as those between OPC-NPC-like and AC-like cells (Topic 9–25, 44–34, 44–36, 9–19), demonstrated closer spatial proximity (**Fig. 4I-J**). In contrast, Topic pairs with weak or absent transition links, particularly involving Neuronal-like and Gliosis-Hypoxia states (Topic 24–19, 24–2, 37–44), exhibited marked spatial separation (**Fig. 4I-J**).

We also analyzed transition trajectories across pseudotime-aligned clonal cell populations in our dataset^66^, finding that net transition scores (TAP-TRS) aligned with temporal co-occurrence of Topics (**Extended Data S8A-C, Supplementary Table 5**, Methods). Notably, the regulatory logic enabled by scDORI topic-specific eGRNs showed stronger concordance with cell state plasticity, spatial organization, and pseudo-temporal transition patterns when compared to either averaged Topic-specific eGRNs from scDORI or eGRNs inferred through alternative methods (**Extended Data S8D-F;** Methods), demonstrating the advantage of continuous Topic-resolved modeling. Together, these findings suggest that GB cells transition along structured and spatially coherent trajectories, largely shaped by the regulatory architecture of each state.

The pseudo-temporal analysis of topic co-occurrence also revealed how Neuronal-like cells, despite their repressive barriers, can transition into other GB states. Across all trajectories (N=60 clones; Methods), Neuronal-like cells preceded either Proliferative or OPC-NPC-like cells (**Fig. 4K and Extended Data Fig. 8B)**. State changes along these transition routes were accompanied by marked changes in the chromatin landscape, whereby TFs from subsequent trajectory stages only become accessible as cells leave the Neuronal-like state (**Fig. 4L and Extended Data Fig 9A-D**). Notably, the transition required the shutdown of key plasticity repressors (MYT1L, ST18, POU2F2, NR3C2) that otherwise silenced transition-initiating TFs (PBX1, SOX4) (**Fig. 4M and Extended Data Fig. 9A-D**). Silencing these Neuronal-like state-specific repressors, combined with induction of transition-promoting Topics (49, 36, 34), enabled a feed-forward cascade of TF activations driving further state changes (**Fig. 4M and Extended Data Fig. 9A-D**). This silencing of Neuronal-like state-specific repressors persisted throughout subsequent transition stages (**Extended Data Fig. 9C-F**), suggesting their continuous suppression is important for GB progression. This analysis indicates that a fine balance between activation and repression dictates whether Neuronal-like state transitions occur, and gives us valuable insights into how this balance could be tilted in either direction.

### The Repressor MYT1L Restricts GB Plasticity and Promotes the Neuronal-like State

Increased cellular plasticity likely contributes to tumor progression and therapy resistance in GB ^53,54^. Given the comparatively low plasticity of Neuronal-like cells and the predicted role of transcriptional repression in restricting GB plasticity in patients, we sought to investigate to what extent we could harness repressors to manipulate GB plasticity (**Fig. 5A**). Of all scDORI-predicted Neuronal-like repressor TFs, MYT1L is the only TF exclusively and highly expressed in over 90% of Neuronal-like cells (**Fig. 5B and Extended Data Fig. 10A**). In addition, MYT1L exhibited the most broad predicted repression of alternate GB states among all candidates and was previously described as a safeguard repressor in neurons ^50,55^ (**Fig. 5B, Extended Data Fig. 7A, and Extended Data Fig. 10B**), making it a promising candidate to manipulate.

**Figure 5.**
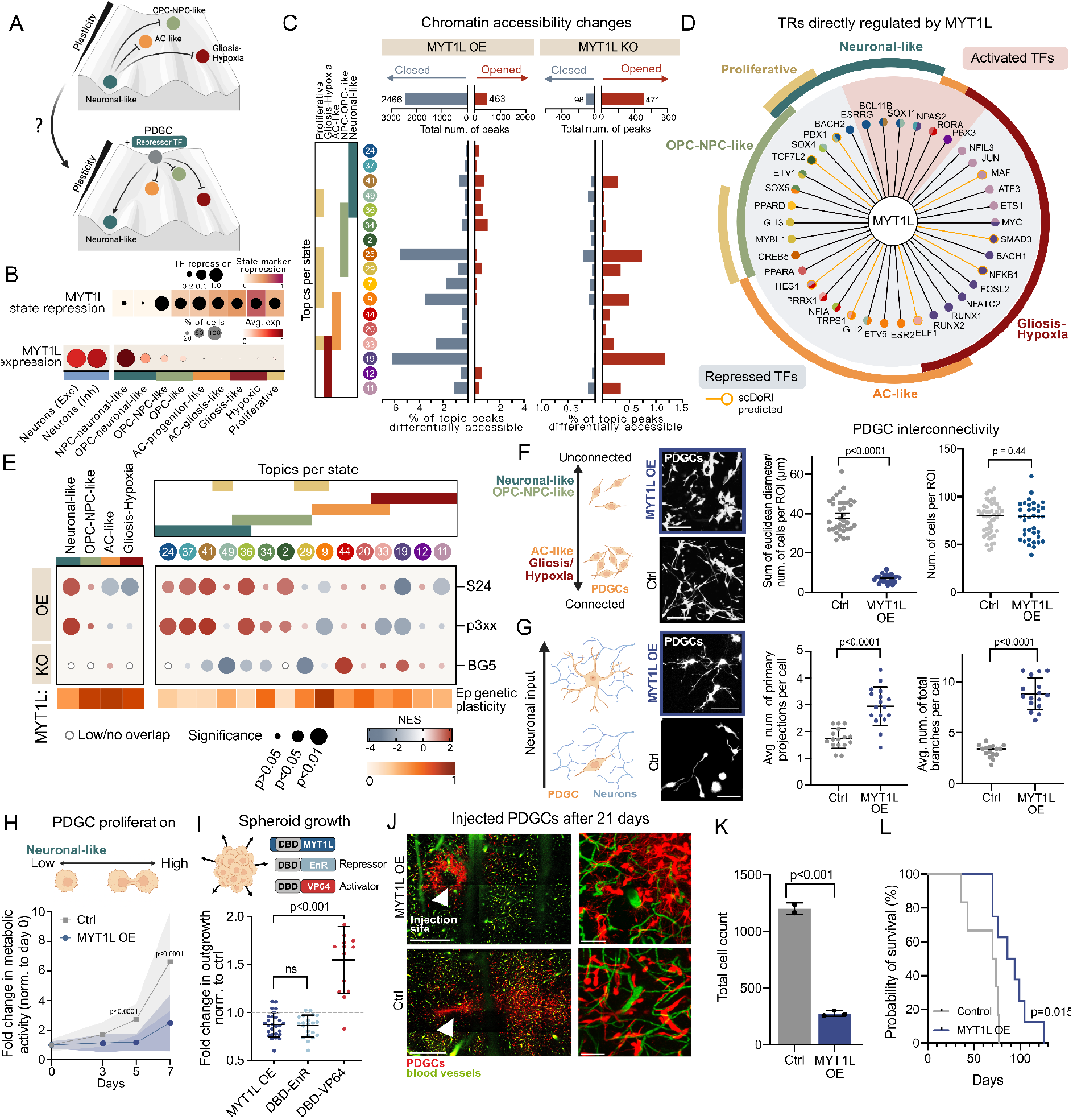
MYT1L Promotes Neuronal-like GB Identity by Repressing Alternate States. (A) Schematic depiction of repressor-mediated decreased plasticity regulation in Neuronal-like GB states and experimental hypothesis that targeted manipulation of repressor TFs of Neuronal-like states enables reprogramming of GB cell plasticity and identity. (B) MYT1L expression and repression patterns. (top) Repression of alternate state genes by MYT1L. Color denotes repression of state marker genes and dot size denotes repression of expressed TFs in alternate states. (Bottom) MYT1L expression in TME neurons or GB states from **Fig. 1A**. Colors indicate average expression, and dot sizes show the percentage of cells with expression per state. MYT1L is exclusively expressed in Neuronal-like GB cells at comparable levels to TME neurons and exhibits strong repression towards all alternate states. (C) Differential chromatin accessibility determined by ATAC-seq following 3 days of MYT1L overexpression (OE) or knockout (KO) in S24 or BG5 patient-derived GB cells (PDGCs) compared to controls, respectively (N=3 per condition, FDR<0.05). The total number of differentially closed or opened peaks (top) and their overlap with top-scoring peaks per Topic (>0.95 Topic-association score) are shown (bottom). Chromatin is predominantly closed in the presence of MYT1L, and opened upon MYT1L deletion. MYT1L remodeled peaks associated with Topics from all other GB states. (D) Selected TRs bound by MYT1L as determined by CUT&RUN sequencing (day 8 post-transduction, N=3 per condition) whose chromatin is differentially opened (red) or closed (blue) upon MYT1L OE in S24 PDGCs. Dot colors denote Topics of activity, and TRs whose regulation was predicted by scDORI are circled in yellow. MYT1L repressed TRs from all alternate GB states. Full list of TRs regulated by MYT1L can be found in **Supplementary Table 6**. (E) Normalised enrichment score (NES) of indicated state and Topic signatures in bulk RNA sequencing data 3 weeks following MYT1L OE (S24, p3xx) or MYT1L KO (BG5) compared to controls (N=3 per condition). The epigenetic plasticity of each state and Topic is displayed as determined in **Fig. 1C**. Neuronal-like identity at the level of state markers and Topic-defining genes is enriched upon MYT1L OE and depleted upon MYT1L KO, respectively. (F) Quantification of tumour microtube (TM) interconnectivity within 2D monocultures of S24 PDGCs with or without MYT1L OE (n = 48 ROIs (Ctrl) and 37 ROIs (MYT1L OE) of 3 independent experiments). The average length of TMs is significantly reduced upon MYT1L OE, leading to significantly less connected morphologies. (Right) total number of cells per region of interest (ROI) with comparable numbers of cells in all areas. P values of two-sided t-test are shown. Scale bar = 200 µm. (G) Average number of primary branches (left) and total neurite-like projections (right) in S24 PDGCs with and without MYT1L OE upon co-culture with primary mouse hippocampal neurons for 7 days (14 days post-transduction, N=3 per condition). The presence of MYT1L significantly increased the number and complexity of neurite-like projections. P values of two-sided t-test are displayed. Scale bar = 100 µm. (H) Proliferation time-course quantification of S24 PDGCs with or without MYT1L OE determined by Alamar blue assay normalized to day 0. N=50 per condition (10 technical replicates x 5 independent experiments). MYT1L OE significantly reduced cell proliferation, p values denote two-way ANOVA Tukey test. (I) Spheroid growth of S24 PDGCs upon doxycycline-induced OE of indicated MYT1L constructs for 7 days, normalized to uninduced controls. OE of full-length MYT1L (N>27) and MYT1L DNA-binding domain (DBD) fused to a repressor (EnR; N>21) reduced spheroid growth, while an MYT1L DBD activator (VP64; N>13) fusion enhanced growth. Data from 3-4 independent experiments, p values of two-sided t-test are reported. (J) Representative micrographs of mouse cortices 3 weeks after injection with S24 PDGCs with or without MYT1L OE captured with 2-photon live imaging. PDGCs are labelled with tdTomato (red) and blood vessels with intravenously injected dextran-FITC (green). Tile scanned region (5000×5000 µm^2^) around injection site (white arrow) and zoomed-in image shows decreased migration and increased complexity upon MYT1L OE. Scale bars = 500 µm (left), 50 µm (right). (K) Quantification of the total number of S24 PDGCs with or without MYT1L OE from (J). The cell count comprises two tile-scanned regions (5000×5000 µm^2^), including the injection site at different depths (200 and 300 µm), per mouse. Mice injected with MYT1L OE PDGCs exhibit significantly less tumor cells (two-sided t-test). N =3 (MYT1L OE) vs 2 (control). (L) Kaplan-Meier survival curve of mice injected as in (J) with S24 PDGCs with or without MYT1L OE (N=7 vs 6), showing a significant survival advantage in the presence of MYT1L (log-rank test, p=0.015).

To test this, we performed MYT1L gain-of-function in patient-derived GB cells (PDGCs; S24) with no detectable endogenous MYT1L expression, which induced major chromatin accessibility rearrangements with >80% of differentially accessible genomic regions closing upon MYT1L overexpression (OE), consistent with a repressive function (**Fig. 5C and Extended Data Fig. 10C**). MYT1L closed regions contained *cis*-regulatory and/or coding regions of multiple alternate state TRs and marker genes and were strongly associated with Topics from all non-Neuronal-like states (Topics 9, 19, 25) (**Fig. 5C and Extended Data Fig. 10D**). Conversely, MYT1L loss-of-function in PDGCs (BG5) with endogenous *MYT1L* expression caused inverse, albeit less pronounced, effects on chromatin accessibility, with >80% of differentially accessible genomic regions opening upon MYT1L knockout, including multiple regions associated with non-Neuronal-like TRs and markers (**Fig. 5C and Extended Data Fig. 10E**,**F**). Together, this suggests that MYT1L is indeed restricting the epigenetic plasticity of GB cells through transcriptional repression of alternate GB states.

To determine how MYT1L could suppress plasticity and repress such a diverse set of alternate GB states, we identified MYT1L target genes using CUT&RUN sequencing (**Extended Data Fig. 10G and Supplementary Table 6**). Out of all differentially accessible genes in MYT1L-overexpressing PDGCs, 55.5% were directly bound by MYT1L (**Extended Data Fig. 10H**), suggesting it directly induces the majority of the observed effects. As predicted by scDORI, MYT1L directly repressed TRs from Topics of all alternate states, including SMAD3 (Topic 19, Gliosis-Hypoxia), TRPS1 (Topic 9, AC-like), TCF7L2 (Topic 2, OPC-NPC-like) and MYBL1 (Topic 29, Proliferative) (**Fig. 5D, Extended Data Fig. 10H, and Supplementary Table 6**). The smaller fraction of TFs activated by MYT1L was predominantly associated with Neuronal-like Topics (SOX11, ESRRG). At the level of genes, scDORI-predicted MYT1L regulatory effects better than alternative eGRN inference methods, but most notably, 85% of scDORI-predicted TFs downstream of MYT1L were CUT&RUN bound targets (**Extended Data Fig. 10I**). These experiments validate our scDORI-based predictions and show that active suppression of alternate state regulators is a key mechanism by which Neuronal-like cells restrict their plasticity.

Beyond this marked epigenetic remodeling, MYT1L manipulation also induced broad changes at the transcriptional level (**Extended Data Fig. 10J and Supplementary Table 7**). MYT1L loss-of-function in PDGCs (BG5) downregulated several Neuronal-like Topics (Topic 49, 41) and induced genes from alternate state Topics (Topic 44, 19), despite not provoking significant changes in state identity (**Fig. 5E**). Conversely, MYT1L overexpression in PDGCs (S24 and p3xx) with no endogenous MYT1L expression significantly upregulated Neuronal-like state signatures and Topics and caused, to varying degrees, downregulation of alternate states (**Fig. 5E**). Remarkably, induced fates were strongly associated with reduced epigenetic plasticity, while repressed states exhibited higher plasticity, further supporting that MYT1L suppresses plasticity and can shift GB identity towards Neuronal-like states.

This Neuronal-like reprogramming of PDGCs was accompanied by the upregulation of neuron-associated pathways upon MYT1L gain-of-function, including genes related to synaptogenesis and neurite-like outgrowth (**Extended Data Fig. 10K**). To assess if these transcriptional changes translated to functional effects, we assessed characteristic features of the Neuronal-like state upon MYT1L manipulation, such as their low intra-tumoral connectivity and their synapse-driven neurite-like morphology ^11,56^. Indeed, MYT1L significantly impacted the formation of interconnected tumour microtubes within PDGC monocultures, as evidenced by their significantly impaired and enhanced formation upon MYT1L gain and loss-of-function, respectively (**Fig. 5F and Extended Data Fig. 10L**). When co-cultured with primary hippocampal neurons, MYT1L-overexpressing PDGCs exhibited a significantly higher number and branching degree of neurite-like projections, suggesting enhanced neuron-to-glioma crosstalk (**Fig. 5G**) ^11^. Overall, these experiments show that MYT1L-driven epigenetic and transcriptional changes induce functional and morphological shifts in GB cells that promote Neuronal-like features while reducing traits associated with tumour microtube hyperconnected and therapy-resistant states ^57,58^.

To discriminate induced Neuronal-like phenotypes from those of OPC-NPC-like cells, which share functional characteristics ^11,56^, we analyzed the impact of MYT1L on PDGC proliferation, a feature characteristically higher in OPC-NPC-like cells ^10^. Remarkably, MYT1L gain-of-function halted PDGC proliferation, whereas loss-of-function significantly enhanced it (**Fig. 5H and Extended Data Fig. 10M**), suggesting that MYT1L uniquely promotes Neuronal-like identity. These effects were also consistent with MYT1L directly repressing regulators of proliferative Topics (MYBL1, GLI3), as well as TFs promoting transitions towards Proliferative states (PBX1, SOX4) (**Fig. 4M and 5D**). PDGC spheroid growth within tissue-mimicking extracellular matrices was also significantly reduced upon MYT1L overexpression. Importantly, this reduced proliferation and growth upon MYT1L overexpression was phenocopied when fusing the DNA-binding domain (DBD) of MYT1L to a well-characterized repressor (EnR) and reverted when fused to a well-known activator (VP64) (**Fig. 5I and Extended Data Fig. 10N-O**). This shows that MYT1L can induce Neuronal-like identity and reduce tumor cell plasticity and proliferation by a repressive mechanism.

Finally, to assess how restricting plasticity through MYT1L induction might affect tumor growth *in vivo*, we followed MYT1L overexpressing PDGCs orthotopically transplanted in the cortex of mice with two-photon live imaging throughout 21 days (**Fig. 5J**). Notably, MYT1L overexpressing tumors exhibit significantly slower tumor cell growth and restricted brain invasion from the original injection site when compared to control PDGCs (**Fig. 5J-K and Extended Data Fig. 10P**). Ultimately, this significantly delayed tumor development and improved survival outcomes (**Fig. 5L**). This observation is consistent with previous reports correlating higher MYT1L expression with better overall patient survival using TCGA data ^59^. These results demonstrate that we can predict and harness state-specific factors such as the plasticity repressor MYT1L to readily promote transitions towards more benign states, opening the door to future therapies targeting epigenetic plasticity.

## Discussion

Cellular plasticity is a core feature of GB progression, yet the regulatory principles shaping how tumor cells transition between distinct states have mainly remained elusive ^10,60,61^. Leveraging large-scale single-cell multi-omic profiling and a novel computational framework for enhancer-driven gene regulatory network (eGRN) inference, our study delineates a hierarchy of GB cell state plasticity and transition trajectories that are shaped not solely by activation cues but are actively constrained by state-specific repression networks.

To uncover the regulatory mechanisms underlying GB heterogeneity and transition, we developed scDORI, a deep-learning-based framework that infers continuous eGRNs from paired single-cell RNA and chromatin accessibility data. This approach addresses the limitations of previous methods by offering scalability to large datasets, flexibility beyond predefined cell type annotations, and the ability to model cell state-specific differences in eGRNs, including gene activation and repression ^19,22,25,26^. Applied to our GB multi-ome atlas, we identified several distinct regulatory Topics with unique activation patterns across cells. These Topics correspond to specific gene programs characteristic of their respective cell states, such as neurotransmitter release in neuronal-like cells ^39^ and PDGF signaling in OPC-NPC-like cells^40^. Importantly, scDORI revealed functional heterogeneity within individual GB states, capturing hallmark features such as hypoxic stress and cytokine signaling in gliosis-hypoxia cells ^42,43^. We identified many transcription factors likely driving specific GB cell states and mapped how they regulate one another. This revealed a complex but structured landscape of possible state transitions which correlated with the level of epigenetic plasticity of each state. These findings are consistent with prior reports of state-specific features and developmental-like differentiation programs in GB, yet extend them by mapping the underlying gene regulatory logic and transcription factors that control them.

Among the most epigenetically confined populations, Neuronal-like cells deploy a distinct mechanism of plasticity suppression. This is orchestrated, at least in part, by MYT1L, a transcriptional repressor with a well-established role in neuronal differentiation and development ^50,55,62^. We found that MYT1L binds and represses regulators of all alternate malignant states, including AC-like (TRPS1, STAT3), Gliosis-Hypoxia-like (FOSL2, BACH1), and OPC-NPC-like (ASCL1, SOX10) programs, thereby locking cells in a low-plasticity neuronal-like state. Unlike a passive absence of permissive cues, this repressive architecture actively silences potential transitions, forming a barrier to phenotypic conversion. Functional experiments in patient-derived GB cells validate MYT1L as a key factor in this repressive program. Overexpression of MYT1L induces widespread chromatin compaction, suppresses alternate state regulators, and enforces a transcriptional and morphological shift toward a neuronal-like phenotype, marked by reduced proliferation, loss of tumor microtube connectivity, and an increase in neurite-like outgrowth. Conversely, MYT1L loss leads to chromatin opening and derepression of alternate fate regulators, increasing proliferative capacity and tumor microtube connectivity associated with highly aggressive mesenchymal-like states ^57,63^.

These findings position MYT1L as a factor suppressing plasticity in GB that can reshape chromatin landscapes, curb tumor growth, and stabilize less aggressive intrinsic states. Upon transplantation in mice, MYT1L overexpression delays tumor progression and prolongs survival *in vivo*, suggesting a therapeutic relevance for targeting MYT1L. This is supported by TCGA patient data and previous studies showing that MYT1L suppresses glioma formation in neural stem and GB cells ^59,64^. Critically, we find that the functional effects of MYT1L depend on its repressor activity, as demonstrated by engineered fusions of the MYT1L DNA-binding domain with transcriptional repressors and activators, underscoring that repression is the key mode of plasticity control in this context and validating our computational model prediction. The logic of MYT1L-mediated repression mirrors emerging safeguard mechanisms in development and cancer, where lineage-specific repressors maintain cell identity by silencing alternative fates ^1,65^. These findings highlight how tumor cells can co-opt developmental repressors to fortify stable, low-plasticity phenotypes. From a cancer perspective, similar factors that suppress plasticity by repression have been reported in the liver and colon ^29,30^, suggesting that such factors can be found and manipulated in different tumor contexts.

Our findings complement and extend our companion study^66^, which resolved a conserved spatiotemporal trajectory of glioblastoma cells transitioning from developmental-like to gliosis and hypoxia-associated states across patients. This study mapped cell state transitions by integrating single-cell, spatial transcriptomic, and spatial whole-genome sequencing data. Here, we provide a missing mechanistic layer, showing that GB cell states differ markedly in their epigenetic plasticity and that state-specific eGRNs shape transitions along the spatial trajectory. Importantly, our Topic-based inference framework captures spatial proximity between transcriptionally related cell states and supports the observation that state transitions follow preferred paths rather than arbitrary rewiring. Notably, we validate MYT1L as a Neuronal-like state-specific repressor that establishes an epigenetic barrier to plasticity, thereby anchoring cells at one end of the spatiotemporal axis. Together, our studies unify spatial and regulatory dimensions of GB heterogeneity and suggest that epigenetic restriction, as much as permissiveness, is a determinant of tumor architecture and evolution.

While our study provides significant insights into the regulatory mechanisms of GB heterogeneity, several limitations have to be considered. First, although scDORI offers a robust framework for eGRN inference, its predictions require experimental validation to confirm the functional roles of identified regulators, such as MYT1L. Second, our analysis focused on primary GB samples, but investigating recurrent tumors and their therapy response would provide an exciting future perspective. Third, our findings position the Neuronal-like state as a low-plasticity, more benign GB state, but this may not fully capture its role in the GB ecosystem. Future work is needed to test whether this epigenetically constrained state serves specialized, tumor-promoting functions via non-cell-autonomous mechanisms, such as neuron-to-cancer synaptic or paracrine signaling, or even self-stimulatory cancer-intrinsic neural-like “auto-synaptic or auto-paracrine” interactions ^11,45^.

Our findings suggest a revised model of GB plasticity in which tumor cells reside in a structured landscape of epigenetic potential, with transitions governed by enabling and constraining forces. The Neuronal-like state could represent a low-plasticity attractor state actively maintained by transcriptional repression and potentially exploitable for therapeutic reprogramming. Future work should explore whether other repressors operate similarly in alternate states and whether their modulation can tip the balance toward benign or therapy-sensitive configurations. Combining single-nucleus multiome data, plasticity quantification, continuous eGRN inference, and functional validation offers a blueprint for uncovering regulatory bottlenecks in other heterogeneous cancers. Beyond GB, targeting repressors like MYT1L may provide a general strategy to stabilize tumor states, suppress malignant evolution, and enhance differentiation-based therapies.

## Methods

Complete methods and supplementary methods are provided as a separate file (methods.pdf).

## Data availability

The processed GB snRNA datasets can be explored and downloaded from our interactive web portal (www.gbmspace.org/). ATAC datasets will be made accessible on the data portal upon peer review. The raw PDGC next-generation sequencing data is deposited on Gene Expression Omnibus and will be made available upon peer review.

## Code availability

The code to run scDORI is available on (https://github.com/bioFAM/scDORI). Scripts and Python notebooks used to compute plasticity scores and TAP/TRS analysis are available on (https://github.com/PMBio/GBM_analysis).

## Extended data

Extended data figures 1-10 are provided as separate files.

## Supplementary information

Supplementary tables 1-8 are provided as separate files.

**Supplementary Table 1. Epigenetic Plasticity of GB Cells**.

(Tab A) Epigenetic plasticity of GB meta-cells, computed as described in **Fig. 1B**.

(Tab B) Epigenetic plasticity of the top 5000 cells per Topic. Values at the cell level were determined by assigning the same epigenetic plasticity value to all cells that make up a meta-cell from (Tab A).

**Supplementary Table 2. scDORI Topics**.

(Tab A) Average activity of scDORI Topics in each GB and TME state.

(Tab B) Topic-association score for all genes used for scDORI training.

(Tab C) Enrichment of gene signatures and pathways per Topic, as determined by gene set enrichment analysis (GSEA) using genes ranked by respective topic-association score in (Tab B). A comprehensive list of neurodevelopmental and cancer pathways/functions was selected for analysis.

(Tab D) TF activity per Topic. TFs with an activity score above 0.05 are designated Topic Regulators (TRs).

(Tab E) TF repressor activity per Topic. TFs with repression score above 0.05, those with the highest repressive effects against genes in alternate Topics, are designated Topic-specific repressors.

**Supplementary Table 3. Inter-Topic Regulatory Interactions**.

(Tab A) Topic Activation Potential (TAP) and (Tab B) Topic Repression Score (TRS) between pairs of selected GB Topics, reflecting the cumulative activating or repressive influence of a Topic over every other Topic. Pan-state Topics are excluded since their non-specific activity across cells limits the interpretation of resulting links.

(Tab C) TFs driving activating and (Tab D) repressive interactions between indicated Topics.

(Tab E) State transition potential derived from the cumulative regulatory effects of all Topics in a state towards Topics active in every other state.

**Supplementary Table 4. Spatial Organization of Topics**.

(Tab A) Average spatial distance between every pair of Topics per Visium section. The metric represents euclidean distance between Visium spot coordinates. Distance of 1 is equivalent to 100µm.

(Tab B) Average spatial distance between Topics across all Visium sections.

(Tab C) Spatial distance between states, as determined by aggregating spatial distances between state-associated Topics (Methods).

**Supplementary Table 5. Temporal Co-activation of Topics**.

(Tab A) Average temporal co-activation of indicated Topics across pseudo-temporally ordered clonal populations in the dataset (N=60). Values denote average cosine similarity between the temporal activation of each Topic (Methods).

(Tab B) Temporal overlap between states, as determined by aggregating Topic co-activation values across state-associated Topics.

**Supplementary Table 6. MYT1L Chromatin Binding and Differential Chromatin Accessibility in GB Cells**.

(Tab A) MYT1L bound chromatin peaks determined by CUT&RUN sequencing upon MYT1L overexpression for 8 days in S24 PDGCs.

(Tab B) Differentially accessible chromatin regions determined by ATAC-seq upon MYT1L overexpression for 3 days or 8 days (Tab C) in PDGCs (S24). Fold change, significance and information on the nearest gene are provided for each peak. Peaks that target TRs and whose nearest gene is also bound by MYT1L are labeled.

(Tab D) Differentially accessible chromatin regions determined by ATAC-seq upon MYT1L knockout for 3 days PDGCs (BG5). Fold change, significance and information on the nearest gene are provided for each peak. Peaks that target TRs and whose nearest gene is also bound by MYT1L are labeled.

**Supplementary Table 7. Differentially Expressed Genes upon MYT1L Manipulation in GB Cells**.

(Tab A) Differentially expressed genes determined by RNA-seq upon MYT1L overexpression for 21 days in S24 and p3xx (Tab B) PDGCs.

(Tab C) Differentially expressed genes determined by RNA-seq upon MYT1L knockout for 21 days in BG5 PDGCs.

(Tab D) Significantly enriched pathways upon MYT1L overexpression in S24 and p3xx (Tab E) PDGCs or knockout in BG5 (Tab F) PDGCs determined by gene set enrichment analysis (GSEA).

**Supplementary Table 8. Viral Vectors, DNA Oligos, and Antibodies Used in Study**.

(Tab A) List of Lentiviral vectors.

(Tab B) List of DNA Primers.

(Tab C) List of Antibodies.

## Acknowledgments

We acknowledge the DKFZ Center for Preclinical Research (A. Riedasch), Small Animal Imaging (M. Jugold), Light Microscopy Core (F. Bestvater), Sequencing OpenLab (N. Glaser), Genomics Core Facility (A. Schulz), and the Heidelberg University Nikon Imaging Centre (C. Ackermann) for excellent service. We thank Judith Zaugg, Aryan Kamal, Pau Badia-i-Mompel, Kai Ueltzhoeffer, Brian Clarke, and Danai Vagiaki for input on eGRN modelling and Donald Hansen for input on figure design. We thank M.M., O.S., O.A.B., W.W., and F.W. lab members for critical discussions. Fellowships were provided by the Helmholtz International Graduate School (to M.S.) and the DAAD (to L.R.G.). M.M. was supported by the State Parliament of Baden-Württemberg for the Innovation Campus Health + Life Science Alliance Heidelberg Mannheim, the German Research Foundation, the Hector Stiftung II gGmbH, and ERC StG (804710). O.S was supported by the German Federal Ministry of Education and Research (BMBF) through the project “DeepSC2” (031L069A). This work was supported by Wellcome Leap as part of the Delta Tissue programme (O.S. and O.A.B.) and funded by the Wellcome Trust Grant institutional grants to the Wellcome Sanger Institute (206194 and 220540/Z/20/A).

## Author contributions

M.S., L.R.G., E.H., O.A.B., O.S., and M.M. conceptualized the study. M.S. conceptualised, developed and applied scDORI to GB atlas. M.S. and L.R.G. developed the TAP metric and performed all data analysis for multi-ome data. L.R.G, E.H., J.S., C.S., D.C.H., E.J., T.K., B.W., B.L. and S.W. performed the experiments. T.G. and F.M. generated multiome data with contributions from O.G.. R.M. provided GB tissue samples. G.D. performed multi-ome data pre-processing, analysed spatial data and contributed to interpretation of GB cell-states. M.S., L.R.G., and E.H. interpreted the results with inputs from O.A.B, O.S., and M.M. K.M. and M.R. provided support in benchmarking scDORI. L.R.G. made the figures with inputs from M.S.. W.W. and F.W. provided resources. O.A.B., O.S., and M.M. secured the funding. M.S., L.R.G. O.A.B., O.S., and M.M. wrote the manuscript with contributions from all authors.

## Ethics declarations

O.S. is a paid advisor of Insitro. The other authors declare no competing interests.

