## Supplementary methods, extended data figures and supplementary tables for "Decoding Plasticity Regulators and Transition Trajectories in Glioblastoma with Single-cell Multiomics": extended_data_figs.pdf

Extended Data Figures 1-10 on the next pages.

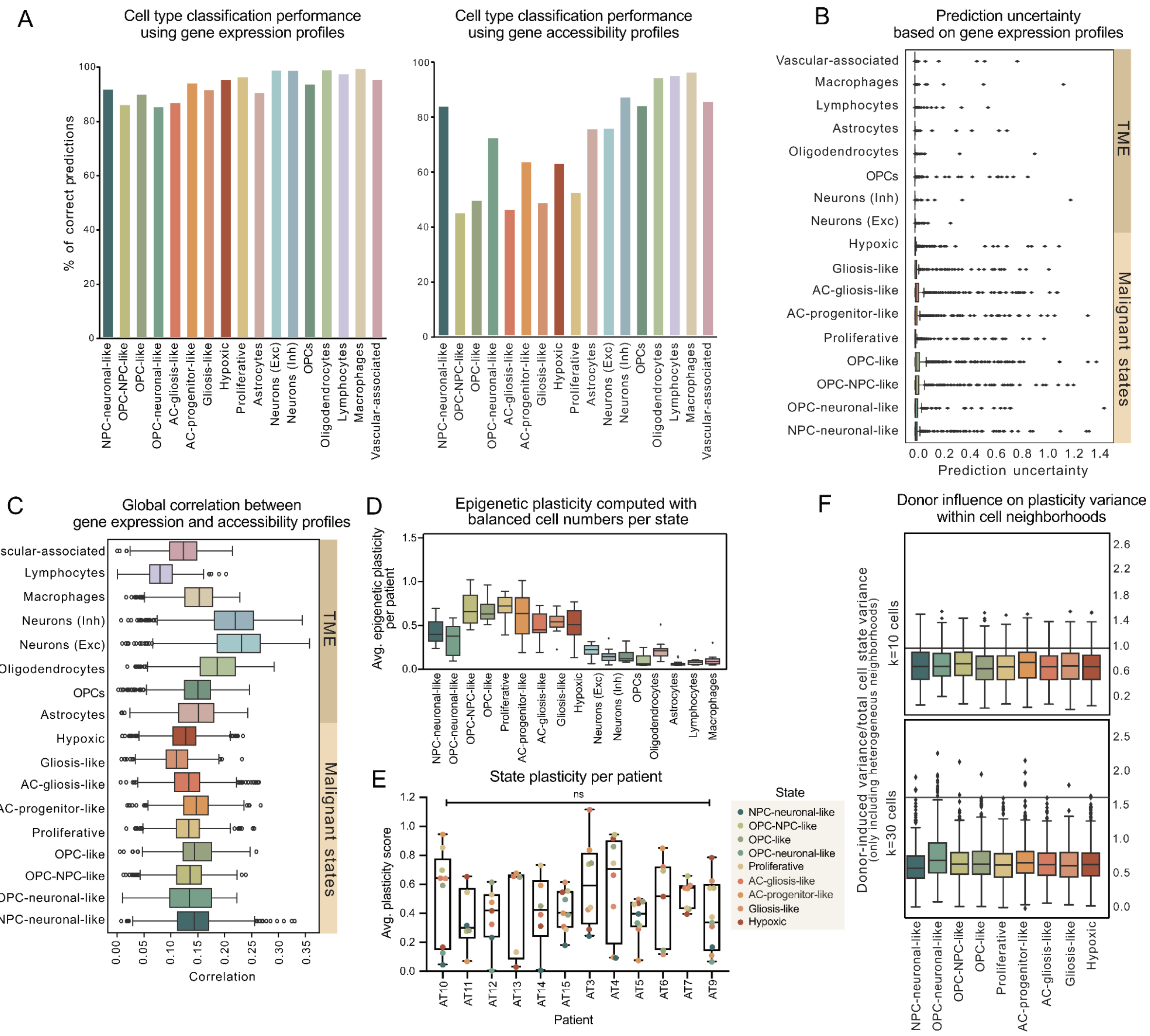


**Extended Data Figure 1. Quantification of Epigenetic Plasticity and Robustness of Estimates**

(A) Cell-state prediction accuracy of a linear classifier trained on gene expression levels. Predictions were tested with either gene expression (left) (5-fold cross validation) or gene accessibility (right) as input, evaluated on 9,464 GB and 6,959 TME meta-cells from 12 patients. GB Cell-state predictions from gene accessibility show reduced accuracy.
(B) Uncertainty (entropy) of cell-state predictions based on gene expression data. Low uncertainty in predictions from all cell states confirmed the linear classifier achieves high-confidence cell state predictions from gene expression profiles.
(C) Correlation between gene expression and accessibility across n=21,369 genes in GB meta-cells, stratified by cell state (Methods). Similar levels across GB cell states suggest that variation in epigenetic plasticity is not driven by global coupling between gene expression and chromatin accessibility.
(D) Average epigenetic plasticity per patient for the indicated cell states, computed using a linear model trained on balanced cell numbers per state (as in **Fig. 1E**). Consistent trends across states suggest that the observed differences are not driven by variations in cell counts.
(E) Patient-specific average plasticity scores for malignant GB states. Although individual patients exhibit variation in plasticity across states, the average plasticity is comparable across patients (one-way ANOVA, p-val=0.53).
(F) Variance in plasticity scores within defined neighborhoods of cells across donors versus across cell-states, shown as a ratio. Values were computed using k=10 neighbors or k=30 neighbors for each metacell, stratified by state. Donor-driven variance in plasticity is lower than state-driven variance in all malignant states (Methods).

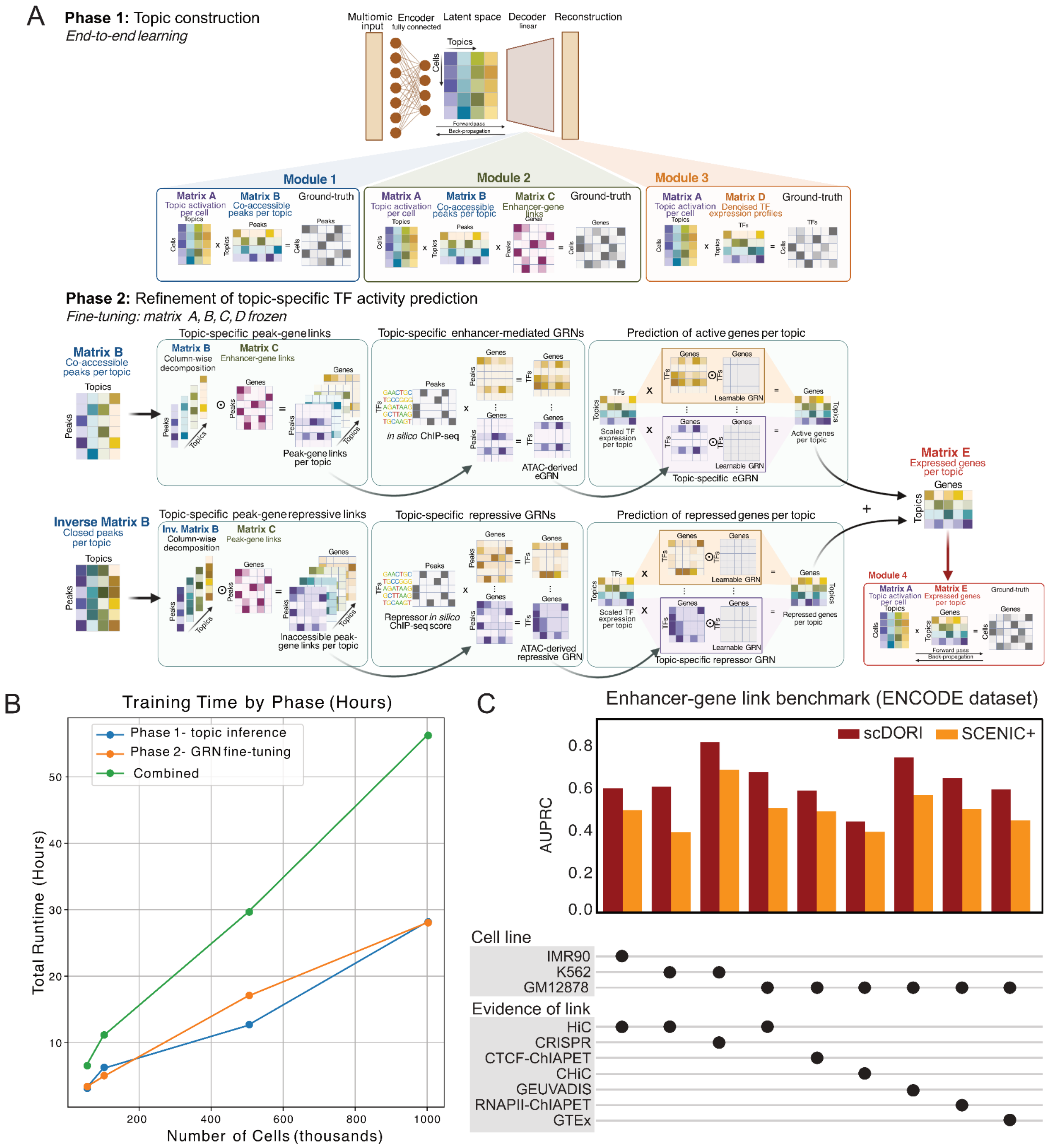


1

2 **Extended Data Figure 2. scDORI Model Architecture and Benchmarks**

3 Figure legend on next page.

(A) Phase 1: Topic Construction via joint training of core autoencoder modules. The autoencoder integrates scRNA-seq and scATAC-seq multi-ome data through a shared latent Topic space. Module 1 reconstructs ATAC-seq peaks using a linear decoder that maps Topic distributions to peak accessibility, with batch-specific adjustments. Module 2 predicts gene expression by combining denoised Topic-specific chromatin accessibility (from Module 1) with a learnable gene-peak linkage matrix, which refines enhancer-gene connections while preserving genomic distance constraints (enhancer links within a fixed gene window). Module 3 reconstructs transcription factor (TF) expression via Topic-TF weights, ensuring latent Topics capture TF co-expression patterns. All modules are trained jointly with reconstruction losses and regularization to improve interpretability of latent space (Methods).
Phase 2: Fine-tuning for eGRN inference. A fourth module learns Topic-specific TF-gene regulatory links by integrating three key inputs: (1) *in silico* TF-peak binding scores (activator/repressor), (2) gene-peak accessibility linkages (from Module 2), and (3) TF expression levels (from Module 3 or true TF expression from data). Topic-specific TF-gene links are computed by weighting TF-peak interactions by chromatin accessibility (from Module 1) and linking peaks to target genes. Learnable TF-gene-Topic matrices further adjust these links using TF-gene expression co-variation. Activator and repressor contributions are combined into a net regulatory score, which predicts gene expression using Topic-aggregated TF expression scores (Module 3). During fine-tuning, the encoder, peak-gene links, and Topic-peak decoder may be frozen to stabilize latent Topics, with additional sparsity constraints on learnt TF-gene weights (Methods).
(B) Computational requirements for training scDORI, considering Phase 1 and Phase 2 training separately or combined. scDORI can be applied to datasets with millions of cells. The empirical runtime on subsampled multi-ome data from the GB atlas is shown.
(C) Assessment of peak-gene link predictions using a semi-synthetic ENCODE single-cell dataset [1]. The Area Under the Precision-Recall Curve (AUPRC) is shown for predicting true peak-gene pairs using scDORI or SCENIC+ (Methods).

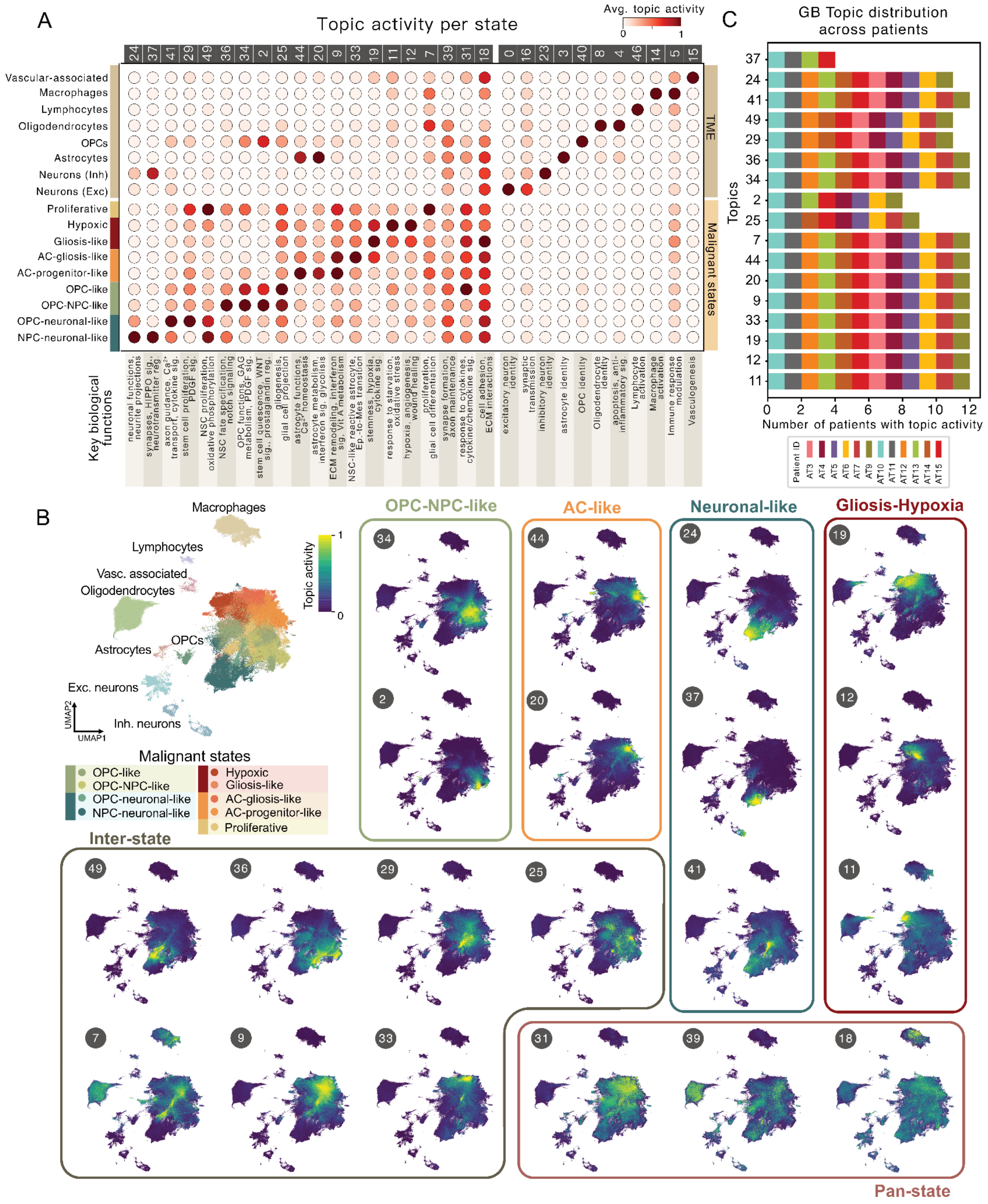


**Extended Data Figure 3. scDORI Topic Activity Patterns Across Cell States and Patients**

(A) Average Topic activity in malignant and TME states. Extended version of **Fig. 2C** including all Topics. Key biological functions of each Topic based on Gene Set Enrichment Analysis (GSEA) are shown. Raw values and complete pathway analysis are reported in **Supplementary Table 2**.
(B) Activity of GB-associated Topics in all cells visualised on the UMAP representation of **Fig. 1A**. State-specific, inter-state, and pan-state Topics are labeled accordingly.
(C) Activity of GB-associated topics across patients. Shown are the number of patients with at least 1% of their cells exhibiting activity (>0.05) for each Topic. All GB Topics are active in at least 4 out of 12 patients, confirming that Topics capture universal regulatory programs of GB.

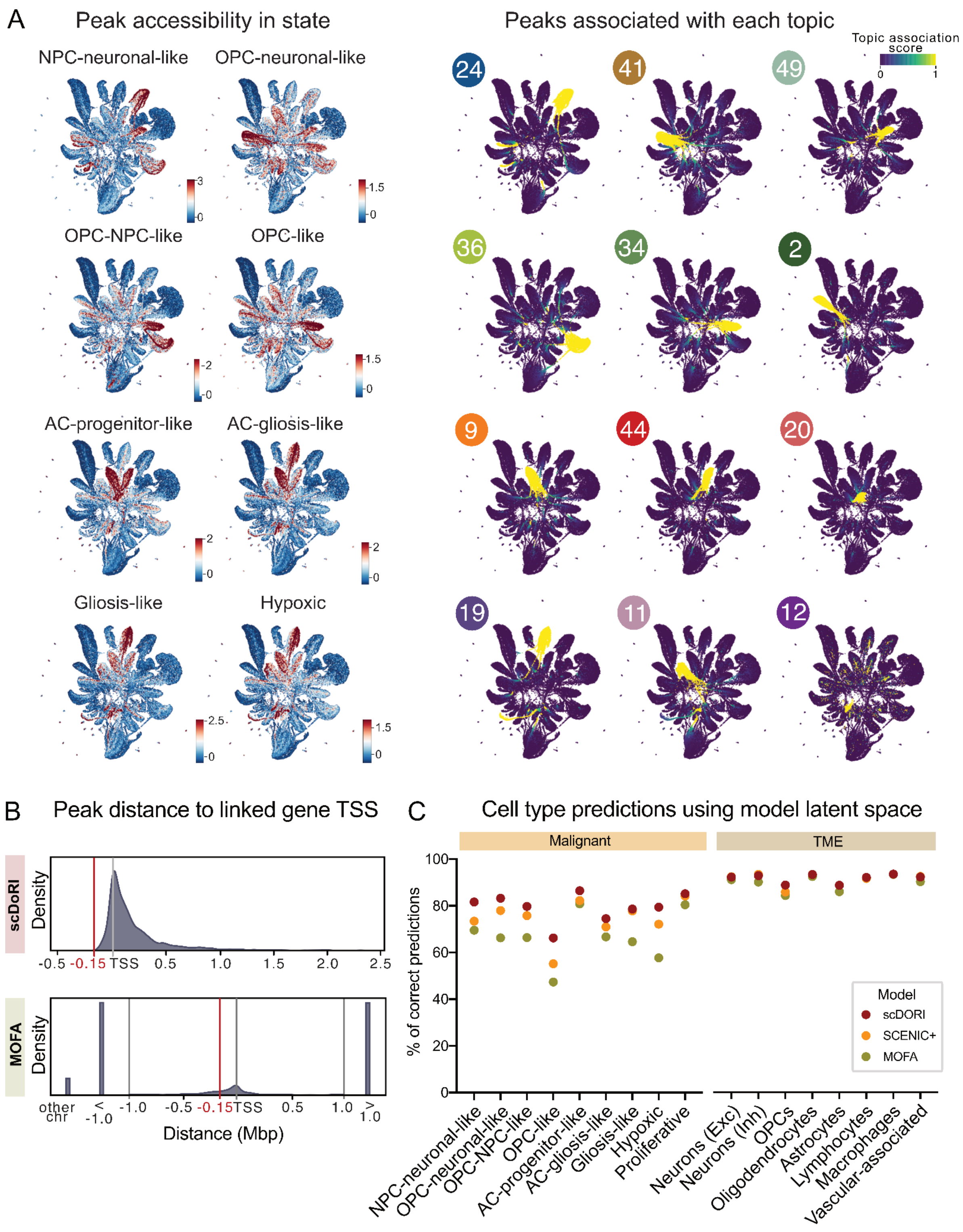


**Extended Data Figure 4. Assessment of scDORI Regulatory Relationships and Latent Space**

Figure legend on next page.

(A) UMAP visualization of ATAC-seq peaks in the dataset using Topic-peak scores (from scDORI module 1) as embedding. (Left) Z-scored accessibility of peaks in each GB state. (Right) Topic-peak score for selected Topics. Topics identify distinct groups of co-accessible peaks within each GB state, reflecting different gene regulatory programs active in those states.

(B) Genomic distance distributions between peak-gene pairs inferred by scDORI or MOFA [2]. For each peak in a MOFA factor, we picked the closest gene that was also part of the same factor. Red lines mark the *cis*-distance (150kb) constraint applied to scDORI. Black lines indicate the ±1 Mb assessment window, and interchromosomal peak-gene pairs are shown separately on the left. TSS: transcription start site.

(C) Comparison of predicting cell-type annotations in our GB dataset using three latent space embeddings: scDORI (all 50 Topics), MOFA (50 factors), and SCENIC+ (AUCell-based TF activity scores from 195 TFs). Logistic regression models were trained using a five-fold cross-validation scheme to predict the cell-type label using the latent embedding, considering 9,464 glioblastoma (GB) metacells and 6,959 tumor microenvironment (TME) metacells as input. Despite their mechanistic constraints, scDORI Topics outperform other latent spaces in capturing transcriptional variation across cell types.

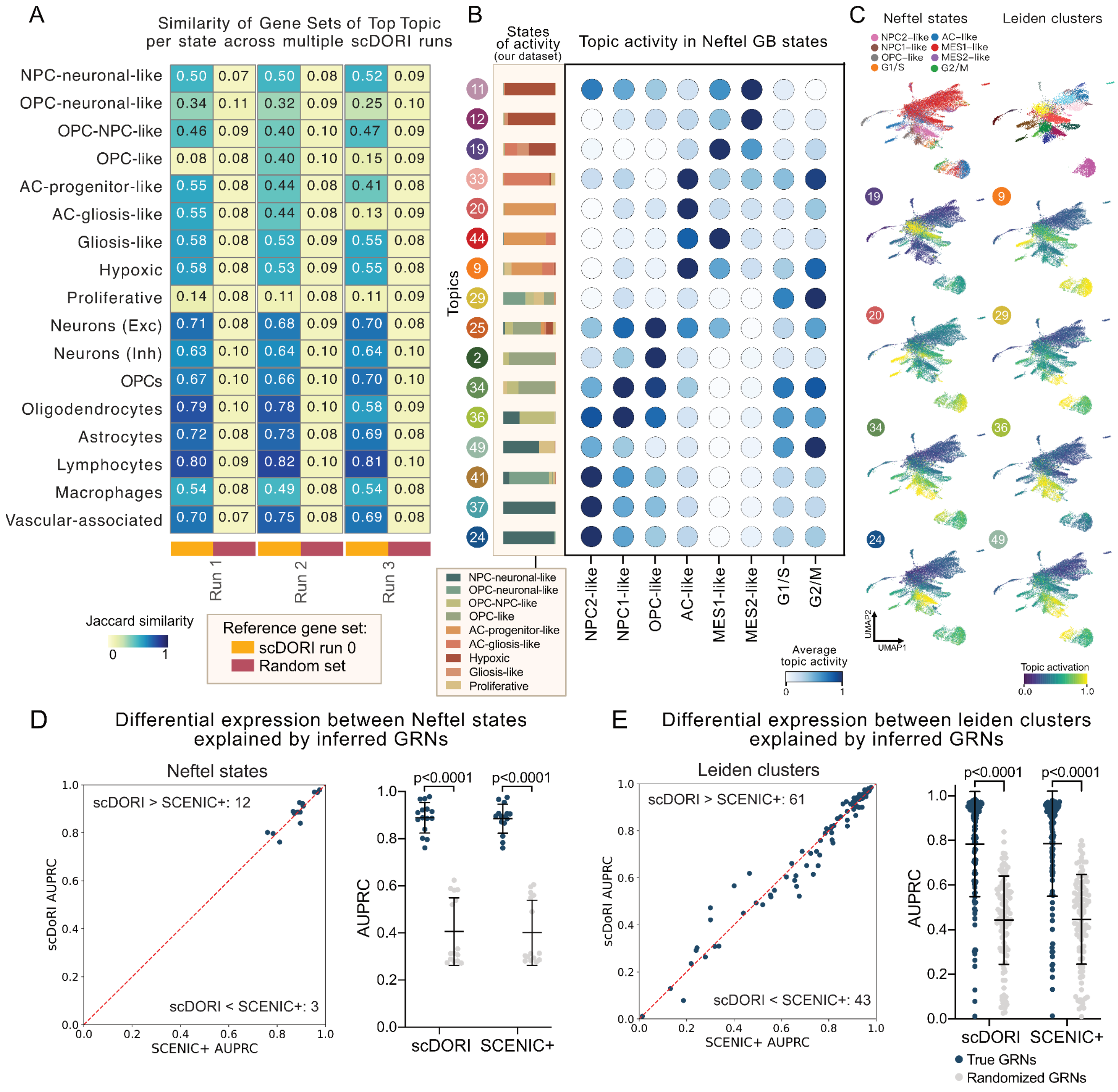


**Extended Data Figure 5. Robustness of scDORI Topics Across Multiple Runs and Datasets**

Figure legend on next page.

(A) Jaccard similarity between genes associated with the top Topic per state across multiple scDORI runs (with different random seeds for initialisation). Topic-associated gene sets of three additional scDORI runs were compared to those of the reference run (original Run 0) and a randomized version of the reference Topic-gene set as control. Top 500 associated genes were considered for each comparison. Overall, top Topics per state capture consistent gene programs across multiple runs.

(B) Activity patterns of selected scDORI Topics derived from our GB atlas (de Jong et al., co-submitted manuscript) projected onto a reference single-cell GB atlas by Neftel et al.[3]. Color denotes average Topic activity across cell state annotations. The bars on the left show the proportion of cells from each state among the top 5000 cells for each scDORI Topic. scDORI Topics map to highly concordant GB cell populations.

(C) Activity of selected scDORI Topics within malignant cells of the Neftel GB atlas, visualized on a UMAP embedding. Neftel state annotations and Leiden clustered subpopulations are indicated.

(D) Performance of scDORI and SCENIC+ to predict differentially expressed genes between fine-grained GB cell-states (Methods, adapted from [4]). For each pair of states or Leiden clusters, differentially expressed genes (DEGs) were identified. A logistic regression model was trained using binary GRN links from upstream TFs as features to predict DEG status. The performance was evaluated using AUPRC with ten-fold cross-validation. AUPRC values are shown for pairs of Neftel cell states or Leiden clusters (as in B), using either SCENIC+ eGRN or scDORI eGRNs (using average TF-gene links from Topics with activation $> 0.2$ in both states/clusters). Randomized eGRNs result in significantly lower AUPRC, serving as a control.

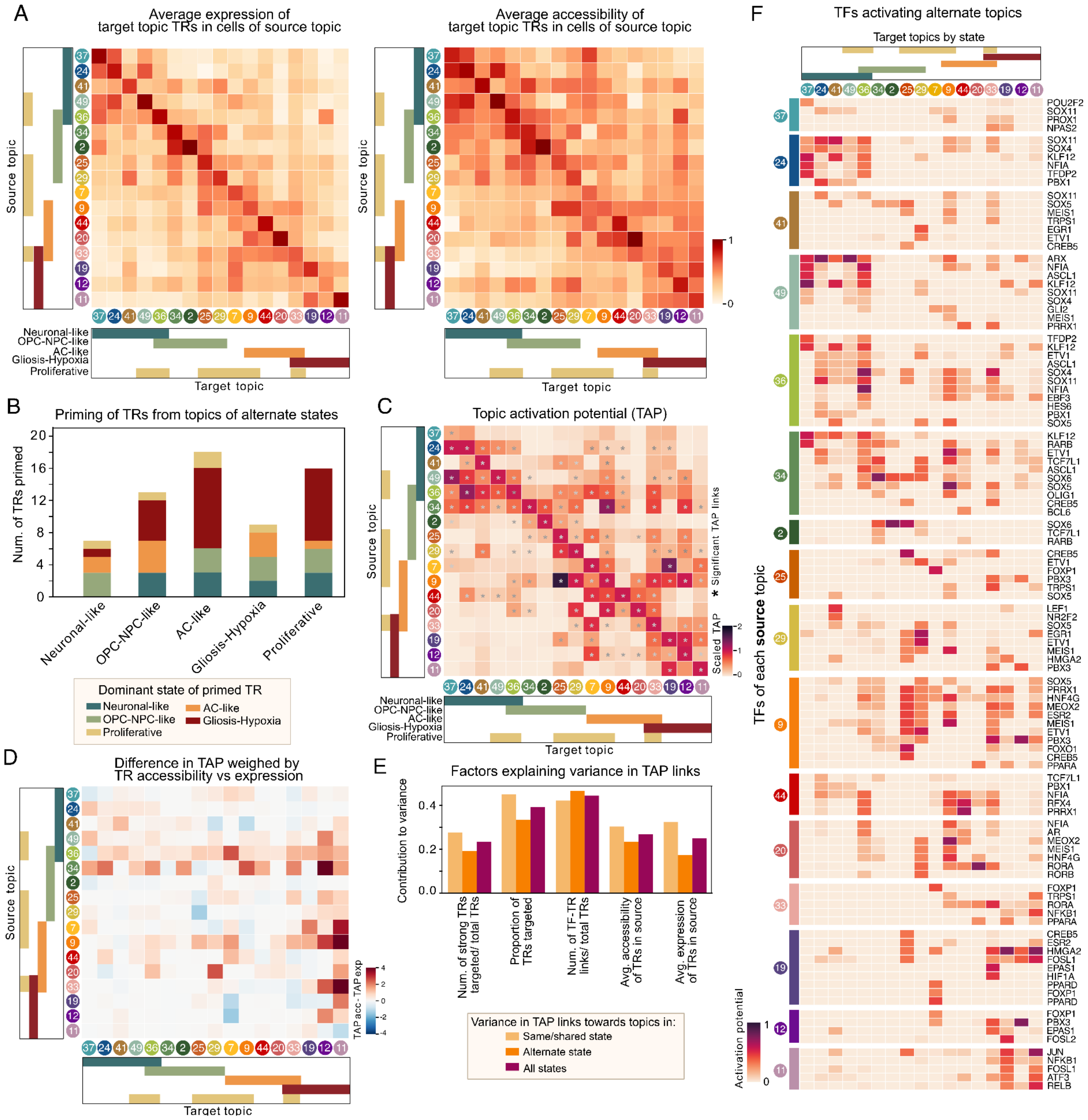


**Extended Data Figure 6. Epigenetic Priming and Topic Activation Potential (TAP) Dynamics in GB**

Figure legend on next page.

(A) Average expression (left) and accessibility (right) of top Topic Regulatos (TRs) (activity score>0.05) from target Topic (columns) in source Topic (rows). While TR expression is mainly confined to Topics within the same GB state, TRs are broadly accessible in Topics of alternate states, suggesting that epigenetic priming enables future activation.

(B) Number of TRs that are epigenetically primed (scaled accessibility>0.5, scaled expression<0.3) in each GB state. Colors indicate the dominant state where the activity of respective TRs is the highest. OPC-NPC-like and AC-like cells exhibit the highest number of primed TRs from alternate state Topics, whereas Neuronal-like cells display the lowest.

(C) Topic Activation Potential (TAP) between pairs of GB Topics, scaled by the number of expressed TFs in the source Topic and weight of self-regulatory links in the target Topic. While OPC-NPC-like Topics (34, 25) and AC-like Topics (9,44) show multiple strong TAP links towards Topics of alternate states, those from Neuronal-like Topics are mostly confined to Topics of their own state. Asterisks denote significant TAP links (>90th percentile of TAP values predicted with random eGRNs).

(D) Difference in TAP scores computed using TR accessibility vs TR expression. Positive values highlight TAP links most influenced by epigenetic priming of TRs, many of which predominantly target alternate state Topics.

(E) Explained variance in TAP scores across Topic pairs by different biological features. Each bar shows the proportion of variance explained by a given feature in an Ordinary Least Squares (OLS) model, stratified by whether the target Topic belongs to the same state (yellow), an alternate state (orange), or any state (purple). Regulatory links inferred from the eGRNs explain the largest share of TAP variance. During transitions between alternative cell states, epigenetic priming of TRs has a greater influence on TAP scores than their expression levels (Methods).

(F) TFs in each source Topic (row grouping) driving the activation (TAP) of other Topics by targeting their TRs. TFs with strong activation of alternate state Topics putatively underlie GB state transitions.

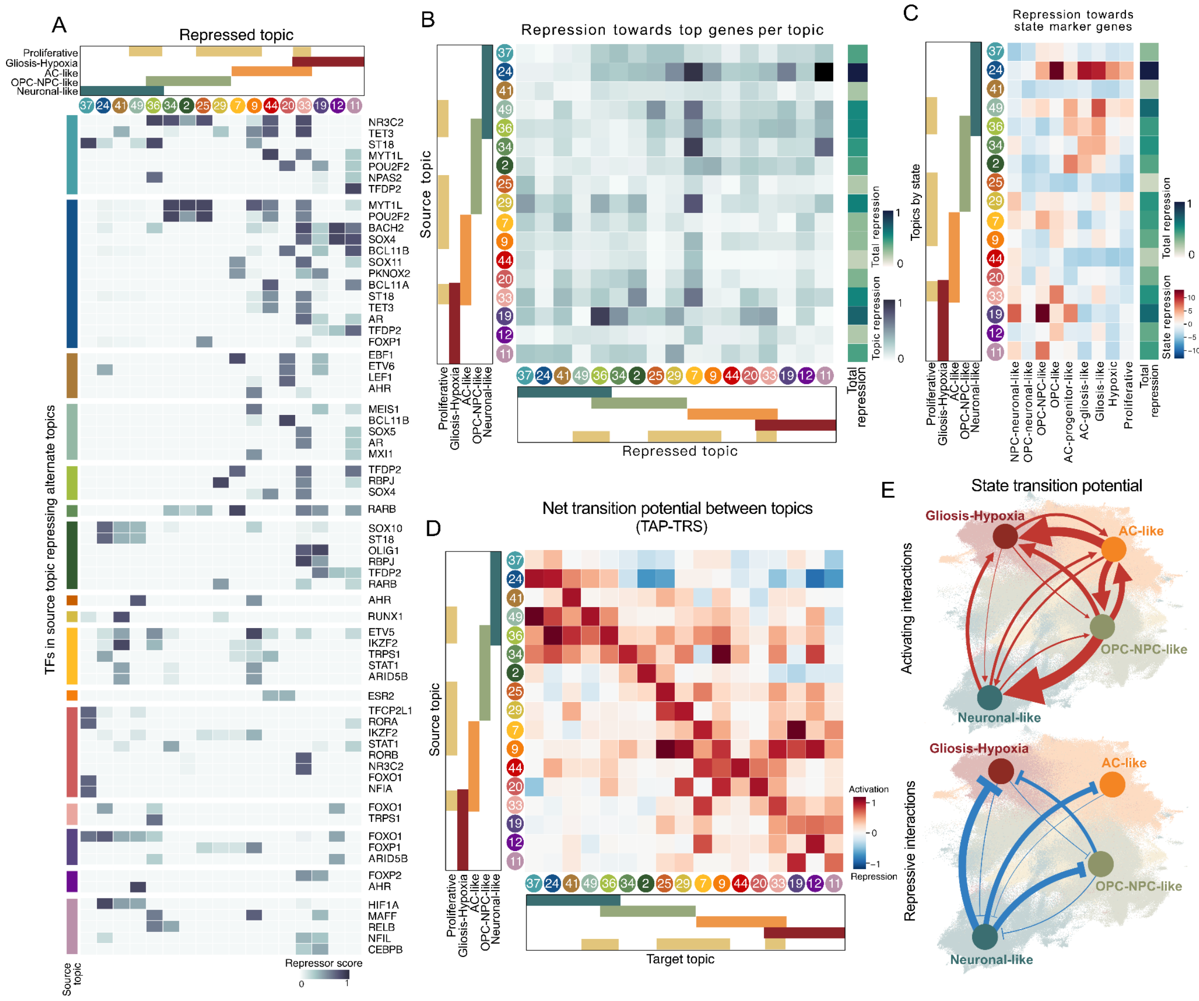


**Extended Data Figure 7. Repressive and Net Regulatory Interactions between Topics**

Figure legend on next page.

(A) Repression exerted by individual TFs in a source Topic (row groupings) toward TRs in other Topics, calculated as shown in **Fig. 4A** (Methods). Green bars indicate TFs with the highest number and strength of repressive links in each source Topic (top 10).

(B) Repression exerted by each source Topic towards the top 50 genes in other Topics, instead of TRs. Gene ranking within Topics was performed based on the Topic-association score predicted by scDORI. Total repression exerted by each Topic is depicted on the right.

(C) Repression exerted by each source Topic towards the top 50 marker genes from each state identified by the linear model in **Fig. 1B**, instead of TRs. Total repression towards alternate states exerted by each Topic is depicted on the right. Similar to TR-level repression, Neuronal-like Topics (24, 49) exert the strongest repression towards genes of alternate Topics and states, although closely followed by Gliosis-Hypoxia Topics (19).

(D) Net Transition Potential between Topics, as determined by the difference between Topic Activation Potential (TAP) and Topic Repression Score (TRS) values between Topic pairs (Methods). Neuronal-like Topics exhibit predominant repressive interactions against alternate state Topics, unlike Topics from other states, where activation is often higher.

(E) Aggregated effect of activating (TAP, top) and repressive (TRS, bottom) interactions between Topics active in the source state towards those in the target state, visualised over UMAP of malignant cells from **Fig. 1A**. Nodes denote states and edge width denotes strength of interaction. Distinct levels of activating and repressive interactions determine transitions across GB states.

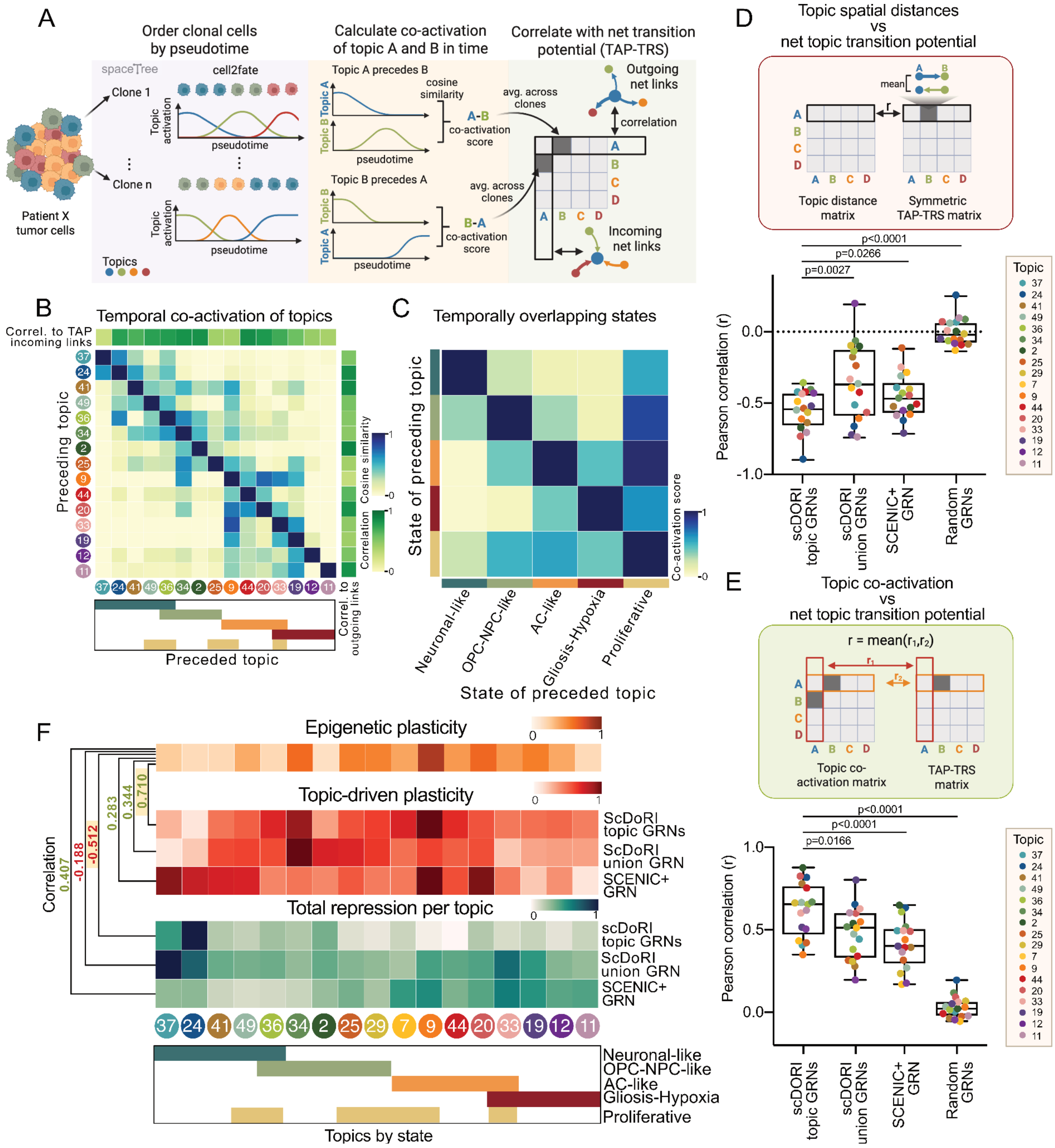


**Extended Data Figure 8. Temporal Co-activation of Topics and Performance of eGRN Inference Methods at Predicting Spatiotemporal and Plasticity Patterns of GB**

Figure legend on next page.

A) Schematic workflow for computing temporal co-activation of Topics by comparing pseudotime-aligned clonal populations to Net Topic Transition Potential (TAP-TRS) values between corresponding Topics (Methods).

(B) Temporal co-activation of Topics averaged across all clones (N=60), determined as described in A (Methods). Clonal Topic co-activation patterns positively correlated with estimated outgoing and incoming TAP-TRS links of each Topic.

(C) Average pseudo-temporal overlap of states, computed by aggregating co-activation scores of state-associated Topics in (B), weighted by the proportional activity of each Topic within a respective state (Methods). Higher values highlight state pairs whose Topics often overlap and are sequentially active, thus suggesting state transitions.

(D) Pearson correlation between Topic spatial distances and Net Topic Transition Potentials (TAP-TRS) associated with each source Topic. TAP and TRS values were computed using Topic-specific scDORI eGRNs, average GRN across all scDORI Topics (union), SCENIC+ eGRNs, or random eGRNs (Methods). TAP and TRS values for each source-target Topic pair were averaged to match the symmetric nature of spatial distances. Net Topic Transition Potential values estimated with Topic-specific scDORI eGRNs inversely correlated with spatial distance and significantly outperformed those derived from alternate methods (paired t-test).

(E) Same as in (D) but correlating temporal co-activation patterns based on clonal populations and Net Topic Transition Potentials (TAP-TRS). Values denote the average between row-wise (outgoing net links) and column-wise (incoming net links) correlations for each Topic. Net TAP-TRS values estimated with Topic-specific scDORI eGRNs positively correlated with Topic co-activation, significantly outperforming those derived from alternate methods (paired t-test).

(F) Comparison of epigenetic plasticity of each Topic (derived as in **Fig. 1C**) with Topic-driven plasticity and total Topic repression modeled using a Topic-specific scDORI eGRNs, an averaged scDORI GRN (union), or SCENIC+ eGRN (Methods). Plasticity and repression models with Topic-specific scDORI eGRNs best recapitulate the observed epigenetic plasticity, exhibiting the expected positive and negative correlation, respectively.

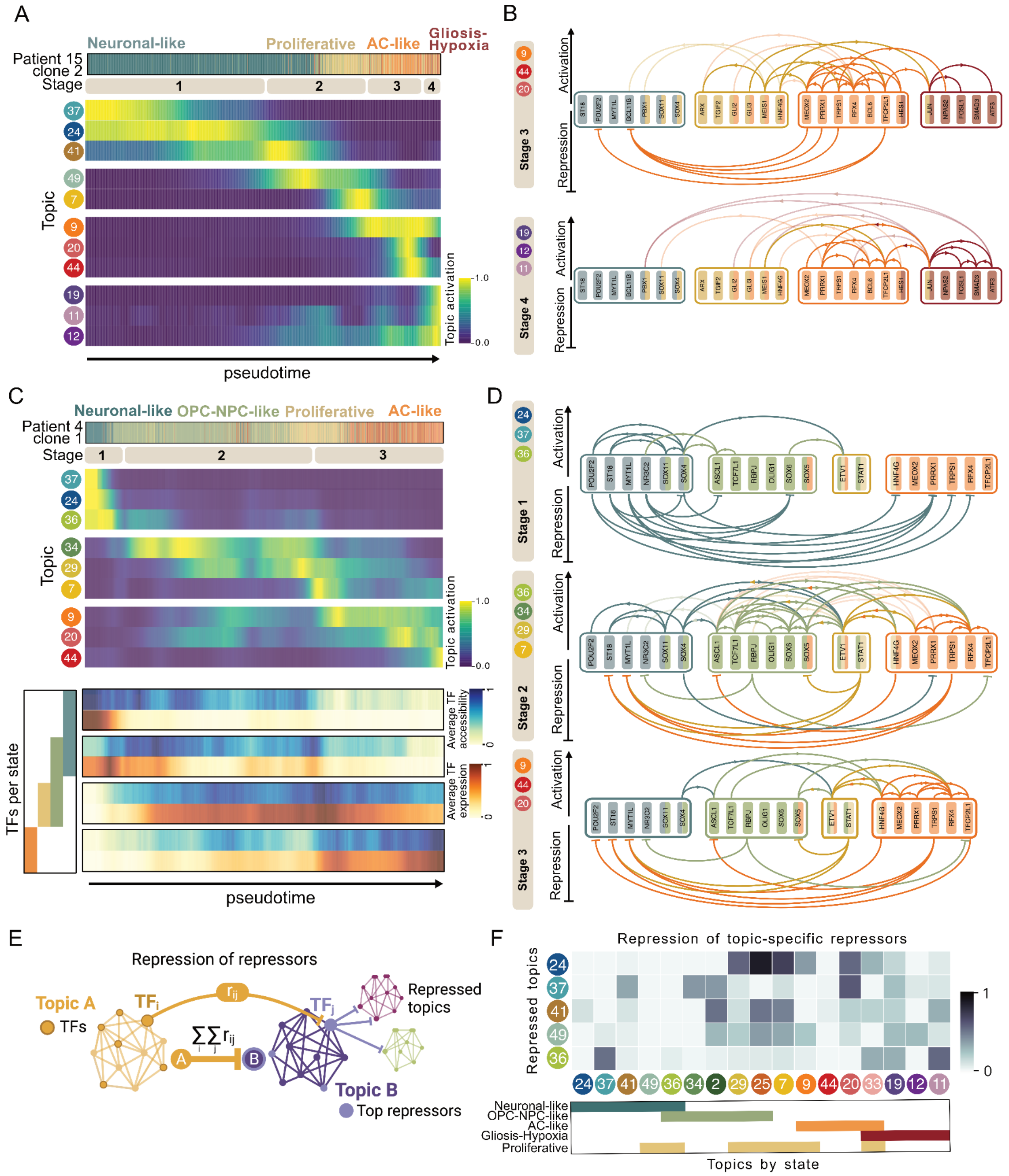


1

2 **Extended Data Figure 9. Gene Regulatory Interactions During Transitions of Neuronal-like GB**
3 **cells**

4 Figure legend on next page.

(A) Pseudotime-ordered clonal population of cells (AT15 clone 2) depicting transition trajectories from Neuronal-like towards AC-like/Gliosis-Hypoxia states via Proliferative state. The trajectory is divided into four stages, defined by distinct Topic activation patterns: (Stage 1) Neuronal-like Topics (e.g., 24, 37, 41), (Stage 2) Neuronal-like-Proliferative transition Topics (e.g., 49, 7), (Stage 3) Proliferative-AC-like transition Topics (e.g., 9, 44, 20), and (Stage 4) Gliosis-Hypoxia Topics (e.g., 19,11,12).
(B) scDORI-predicted activation and repression links between relevant TFs driven by Topics active in Stages 3 and 4 of the trajectory in A. Corresponding links in Stages 1 and 2 are shown in **Fig. 5E**. Links going in the opposite direction to the current trajectory are shaded.
(C) Pseudotemporal ordering of another clonal cell population (AT4 clone 1) transitioning from Neuronal-like towards OPC-NPC-like and then AC-like states. Sequential Topic activation patterns define distinct transition stages: (Stage 1) Neuronal-like Topics (e.g., 24, 37, 36), (Stage 2) OPC-NPC-like-Proliferative Topics (e.g., 34, 29, 7), and (Stage 3) Proliferative-AC-like Topics (e.g., 9, 20, 44). Matched chromatin accessibility and expression of trajectory-relevant TFs are shown across pseudotime. Transitions outward of Neuronal-like state are accompanied by marked chromatin accessibility rearrangements, unlike those from OPC-NPC-like to AC-like.
(D) Predicted gene regulatory interactions of transcription activators and repressors along the trajectory in (C), inferred based on active Topics at each stage. Links going in the opposite direction to the trajectory are shaded. Stage 1 shows dominant repressive links targeting TFs of alternate states, followed by their shutdown through active repression as the trajectory progresses.
(E) Repressive effects from TFs in each Topic towards top repressors in Neuronal-like Topics, computed with a modified version of Topic Activation Potential (TAP)/Topic Repression Score (TRS) computations that consider repressive links towards top repressors, instead of TRs (Methods).
(F) Targeted repression of repressor TFs in Neuronal-like Topics, computed as in (E). Most non-Neuronal-like Topics exert some level of repression against Neuronal-like repressors, reinforcing the hypothesis that their shutdown is required to exit the Neuronal-like state.

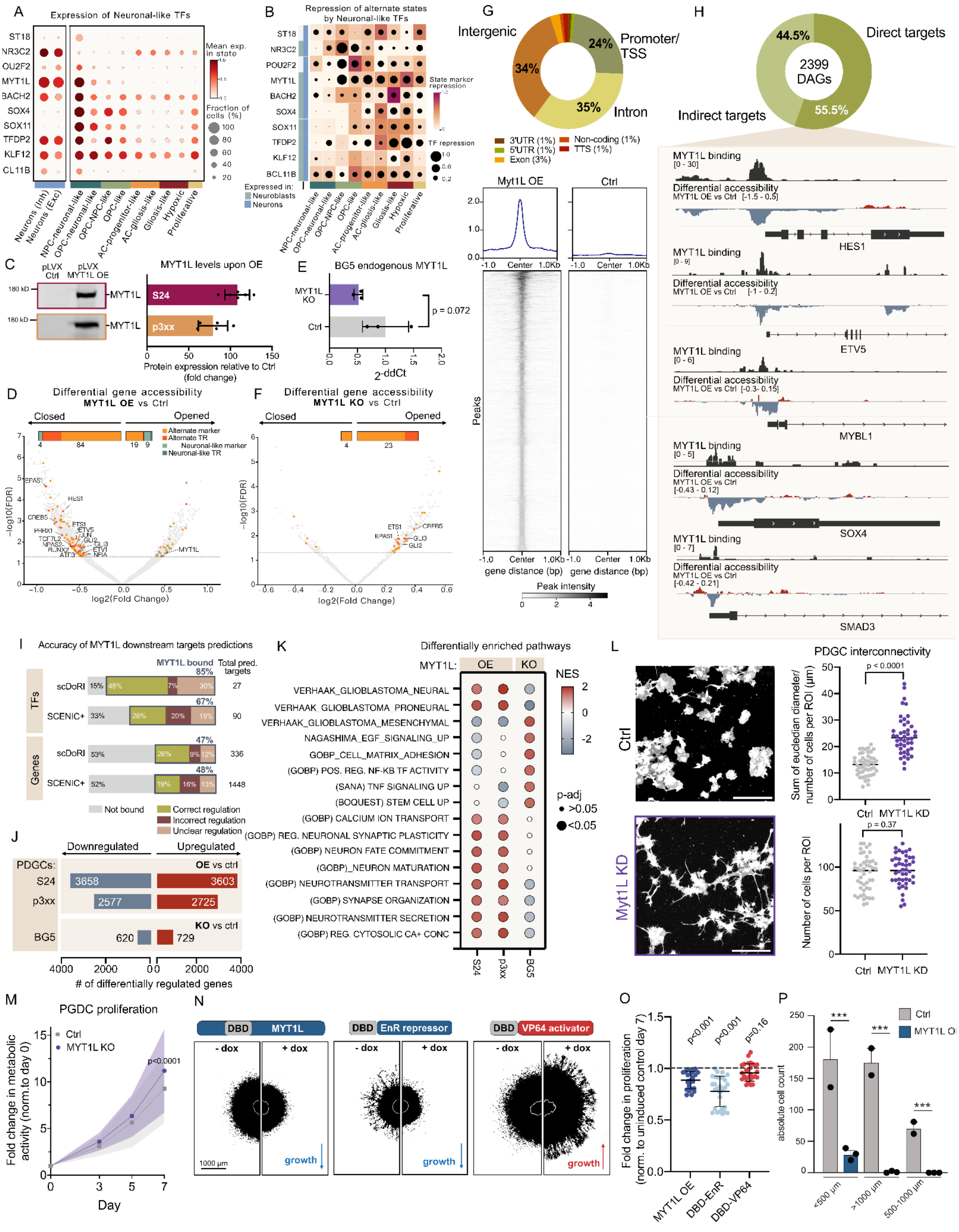


**Extended Data Figure 10. Epigenetic, Transcriptional, and Functional Effects upon MYT1L Manipulation**

Figure legend on next page.

(A) Expression of top repressor TFs in Neuronal-like state in TME neurons (left) or GB states (right) from **Fig. 1A**. Colors indicate average expression, and dot sizes show the percentage of cells with expression per state. MYT1L is exclusively expressed in Neuronal-like GB cells and at comparable levels to TME neurons.
(B) Repression of alternate state genes by top repressor TFs in Neuronal-like cells. Color denotes repression of state marker genes and dot size denotes repression of expressed TFs in alternate states. The bar on the left labels expression in healthy neurons and neuroblasts (CellxGene Portal[5]). MYT1L shows the broadest and strongest repression of alternate state genes.
(C) Representative Western blots (WB) of MYT1L protein levels in S24 and p3xx PDGCs, 7 days upon lentiviral MYT1L overexpression (pLVX MYT1L OE) compared to controls (pLVX-puro), including quantifications shown as Fold change compared to controls (N>5).
(D) Differential gene accessibility upon 3 days of MYT1L OE in S24 PDGCs compared to controls determined by ATAC-seq (N=3 per condition; FDR<0.05). TRs and marker genes from non-Neuronal-like states are labeled in orange, and those from the Neuronal-like state in blue. Selected TRs from alternate state Topics are highlighted.
(E) Quantification of *MYT1L* expression three weeks following MYT1L knockout (KO) compared to controls in BG5 PDGCs via qPCR (N=3 per condition, two-sided t-test).
(F) Differential accessibility of genes three days following MYT1L KO in BG5 PDGCs compared to controls determined by ATAC-seq displayed as in (D) (N=3 per condition; FDR<0.05).
(G) Genomic distribution of MYT1L-bound regions in S24 PDGCs as determined with CUT&RUN sequencing following 8 days of MYT1L OE (N=3) (top). Intensity histogram of MYT1L-bound CUT&RUN peaks in MYT1L OE and control conditions (bottom).
(H) Proportion of differentially accessible genes upon MYT1L OE in (D) that are directly or indirectly regulated by MYT1L determined via CUT&RUN in (G) (top). Representative genome browser track examples of MYT1L-bound TRs closed upon MYT1L OE (bottom).
(I) Precision of MYT1L downstream target prediction with scDORI or SCENIC+ eGRNs at the level of TFs and all genes. An average scDORI eGRN containing predicted MYT1L downstream targets across all Topics was used. Blue boxes highlight the proportion of predicted targets directly bound by MYT1L determined by CUT&RUN in (G). Colors denote the proportion of predicted targets that are MYT1L-bound and are differentially accessible in a consistent (green), inconsistent (brown), or that do not show differential accessibility (light brown) between the prediction and upon experimental manipulation in (D). scDORI enables superior MYT1L regulon predictions than SCENIC+, especially at TF level.
(J) Total number of differentially expressed genes 3 weeks after MYT1L OE in S24 or p3xx PDGCs, or MYT1L KO in BG5 PDGCs determined by RNA-seq compared to controls (N=3 per condition; adjusted p-value<0.05). Full list of differentially expressed genes can be found in **Supplementary Table 7**.
(K) Selected pathways enriched in cells from (J), as determined by gene set enrichment analysis (GSEA). Color denotes normalized enrichment score and dot size denotes significance. Full list of significantly enriched pathways in each cell line is shown in **Supplementary Table 7**.
(L) Representative microscopy images and quantification of tumour microtube (TM) interconnectivity within 2D monocultures of BG5 PDGCs with or without MYT1L KD 3 days after seeding (N=3 per condition; with >44 ROIs analyzed). The average length of TMs is significantly increased upon MYT1L KD, leading to distinctly more connected morphologies. Comparable total number of cells in each region of interest (ROI) confirmed by DAPI quantification. Two-sided t-tests. Scale bar: 200 µm.
(M) Proliferation time-course quantification of BG5 PDGCs with or without MYT1L KO determined by Alamar blue assay normalized to day 0. N=30 per condition (10 technical replicates x 3 independent experiments). MYT1L KO significantly increases PDGC proliferation. Two-sided t-tests.

(N) Representative fluorescent microscopy images of TdTomato-labeled S24 PDGC spheroids transduced with doxycycline-inducible constructs of full-length MYT1L (left), MYT1L DNA-binding domain (DBD) fused with a VP64 activator domain (center), or MYT1L DBD fused with an EnR repressor domain (right). Images of non-induced and doxycycline-induced spheroids were taken at day 7 after embedding and analyzed. Corresponding quantifications are shown in **Fig. 5I**. Scale bar=1000 µm.

(O) Fold change in proliferation of S24 PDGCs with OE of indicated MYT1L constructs in (N) for 7 days normalized to uninduced controls. OE of full-length MYT1L (N>24) and MYT1L DNA-binding domain (DBD) fused to a repressor (EnR; N>27) reduced spheroid growth, while an MYT1L DBD activator (VP64; N>27) fusion enhanced growth. Data from 3 independent experiments, P values of two-sided t-test are reported.

(P) Quantification of the total number of S24 PDGCs with or without MYT1L OE within mouse cortices three weeks following transplantation, grouped by euclidean distance to injection site. MYT1L OE PDGCs reside significantly closer to the injection site (N = 3 MYT1L OE vs N = 2 Ctrl mice, two-way ANOVA with Tukey test).
