## Supplementary methods, extended data figures and supplementary tables for "Decoding Plasticity Regulators and Transition Trajectories in Glioblastoma with Single-cell Multiomics": methods.pdf

April 14, 2025

### Contents

**Contents** **1**

**1 scDoRI** **4**
1.1 Overview 4
1.2 Input Data 4
1.3 Data Preprocessing 5
1.3.1 Pseudobulk Aggregation for TF–Peak Matrix Construction 5
1.3.2 Correlation Estimation 5
1.3.3 Motif Match Calculation 5
1.3.4 Filtering 6
1.3.5 Activators vs. Repressors 6
1.3.6 Integration of Motif and Correlation Signals 6
1.4 scDoRI Model Architecture 6
1.4.1 Encoder 6
1.4.2 Decoder 7
1.4.3 Module 1: ATAC Reconstruction 7
1.4.4 Module 2: RNA Reconstruction from ATAC 7
1.4.5 Module 3: TF Expression Reconstruction 8
1.4.6 Module 4: GRN Inference with Activators and Repressors 9
1.5 Loss Functions 10
1.5.1 ATAC Reconstruction Loss 10
1.5.2 RNA Reconstruction Losses 11
1.5.3 TF Expression Reconstruction Loss 11
1.5.4 Regularization Terms 11
1.6 Training Procedure: Two-Phase Optimization Approach 11
1.6.1 Phase 1: Core Reconstruction (Modules 1–3) 12
1.6.2 Phase 2: eGRN inference (Module 4) 12
1.6.3 Optimization Details and Constraints 13
1.7 Summary of Model Components 13
1.8 Software Usage Guidelines 13
1.9 Downstream Analysis enabled by scDoRI 15
1.9.1 scDoRI Latent Space 15
1.9.2 Topic-Peak Matrix: Co-Accessibility Patterns 16
1.9.3 Topic-Gene Matrix (Module 2) & Gene Set Enrichment 16
1.9.4 Peak-Gene Matrix (Module 2): Enhancer–Gene Links 16
1.9.5 Empirical thresholding of eGRNs 16
1.9.6 Final GRN Construction 17
1.9.7 TF Activity Scores and Topic Regulators (TRs) 17
1.9.8 Identifying Downstream Targets of TFs 18
1.10 Related Works: eGRN Inference and Multi-Omic Dimensionality Reduction 18
1.10.1 Methods for eGRN Inference from Single-Cell Multi-Omics 18
1.10.2 TF–Peak Mapping 19
1.10.3 Peak–Gene Linking 19
1.10.4 TF–Gene Inference 19

**4 Statistical Analysis** **37**

**References** **38**

### 1 scDoRI

#### 1.1 Overview

**scDoRI** (single-cell Deep Multi-Omic Regulatory Inference) *jointly* models single-cell RNA-seq and ATAC-seq data to infer enhancer-mediated gene regulatory networks (eGRNs). Unlike methods that handle dimensionality reduction and eGRN inference as separate steps, scDoRI unifies these tasks so that its latent factors naturally capture coordinated changes in gene expression and chromatin accessibility, without relying on predefined cell-type clusters. Operationally, scDoRI adopts an **Encoder–Decoder** design, reminiscent of variational autoencoders like scVI[2], wherein an **encoder** learns a shared low-dimensional representation, and **mechanistically constrained decoders** map that representation back to observed RNA and ATAC profiles. These decoders are designed to enforce key regulatory logic, such as how open chromatin regions activate nearby genes through enhancer–gene links, and how transcription factors (TFs) act as activators or repressors by binding to regulatory elements.

Through this design, scDoRI learns **topics**: factors that simultaneously group co-accessible peaks, co-expressed genes, and TF regulators into putative TF–gene networks mediated by enhancers. Since scDoRI represents each cell as a *mixture* of topics (instead of forcing cells into discrete clusters), the model can capture continuous, cell-specific changes in GRNs. Moreover, by processing data in mini-batches, scDoRI is scalable to millions of cells.

In the following sections, we detail:

- the input data to the model (Section 1.2),
- data preprocessing to compute in silico ChIP-seq TF-peak matrices (Section 1.3),
- the model architecture, including the encoder and decoder modules (Section 1.4),
- the loss functions (Section 1.5),
- the two-phase training procedure (Section 1.6),
- downstream analysis enabled by the trained model (Section 1.9),
- principles of eGRN inference and multi-omic dimensionality reduction, rationale for design choices in scDORI and comparison to prior work (Section 1.10),

#### 1.2 Input Data

The **scDoRI** framework requires the following input matrices derived from paired scRNA-seq and scATAC-seq (multi-omic) data. Let $C$ denote the number of cells (or metacells), $G$ the number of genes, $P$ the number of peaks in the chromatin accessibility data, $F$ the number of TFs, and $K$ the number of batches or experimental conditions.

1. **Gene Expression Matrix**
$$\mathbf{X}_{\text{RNA}} \in \mathbb{R}^{C\times G}.$$
This contains raw RNA-seq counts for each cell or metacell.

2. **ATAC-seq Counts Matrix**
$$\mathbf{X}_{\text{ATAC}} \in \mathbb{R}^{C\times P}.$$
Each entry corresponds to the number of Tn5 insertions mapped to a specific genomic peak in a particular cell or metacell.

3. **Batch Information**
$$\mathbf{B} \in \mathbb{R}^{C\times K}.$$
This one-hot encoded matrix identifies the batch or experimental condition associated with each observation.

4. **Library Sizes**
$$\mathbf{L} = (\mathbf{L}_{\text{ATAC}}, \mathbf{L}_{\text{RNA}}) \in \mathbb{R}^{C\times 2}.$$
These vectors contain log-transformed library sizes (total counts) for the RNA and ATAC modalities, respectively.

5. **Cell Counts per Metacell**
$$\mathbf{n} \in \mathbb{R}^{C}.$$
If metacells are used, $\mathbf{n}[c]$ reflects the number of aggregated cells in the $c$-th metacell; for single-cell input, $\mathbf{n}$ is a vector of ones.

6. **Transcription Factor Indices:** These are column indices indicating which genes in the expression matrix $\mathbf{X}_{\text{RNA}} \in \mathbb{R}^{N \times G}$ correspond to transcription factors. They are used to extract the TF-specific expression subset $\mathbf{X}_{\text{TF}} = \mathbf{X}_{\text{RNA}}[:, \text{indices}]$, which serves as input to scDoRI's TF-specific decoder branch 1.4.5.

**Note**: While **scDoRI** is designed for single-cell input, users have the option to preprocess the data by aggregating transcriptionally similar cells into metacells/pseudobulks prior to training. This strategy can substantially reduce computational load and training time, particularly for large datasets. Metacell construction is not natively supported in the current implementation. **Importantly, scDoRI performs effectively on standard single-cell multi-ome datasets with >1M cells without the need to aggregate (Extended Data Fig. S2B)**

#### 1.3 Data Preprocessing

A key preprocessing step in **scDoRI** is the computation of signed TF–peak interaction matrices, which serve as a central input for downstream GRN inference. These matrices build upon the "in silico ChIP-seq" framework introduced in [3], which in turn extends **DiffTF** [4], to estimate the functional binding behavior of TFs at regulatory regions.

The core idea is to capture not just where a TF might bind (via its motif), but also how its expression correlates with the accessibility of those sites across the dataset. Specifically, a *positive correlation* between TF expression and peak accessibility suggests the TF is associated with **opening** chromatin when bound—an activator-like signature. Conversely, a *negative correlation* implies that TF expression is associated with **closing** chromatin—suggestive of repressive activity. This modeling choice for repressors adapts prior work in **DiffTF** [4], which treats chromatin closing as a specific signature of repression. We note that while effective, this approach is limited in that it may miss repressive regulatory events that do not involve measurable chromatin closure.

To capture these patterns, we integrate three signals:

1. **Motif match scores** that estimate the likelihood of TF binding to a given peak;

2. **TF–accessibility correlations** to determine whether binding tends to open or close chromatin;

3. **Filtering** using a TF-specific background of non-motif peaks to filter false positives.

The output consists of two matrices:

$$\mathbf{W}_{\text{TF-peak}}^{\text{act}} \in \mathbb{R}^{P \times F} \quad \text{and} \quad \mathbf{W}_{\text{TF-peak}}^{\text{rep}} \in \mathbb{R}^{P \times F},$$

representing predicted TF–peak interactions that are activating or repressive, respectively. These matrices are subsequently used in scDoRI's GRN inference pipeline (Section 1.4.6).

The specific steps to construct these matrices are described below:

##### 1.3.1 Pseudobulk Aggregation for TF–Peak Matrix Construction

To improve the signal-to-noise ratio in estimating TF-peak correlations, we aggregate transcriptionally similar cells into pseudobulks (metacells) prior to computing the signed TF–peak interaction matrices. Pseudobulk construction is performed via high-resolution Leiden clustering on batch-corrected PCA embeddings (using Harmony [5]) from the RNA modality.

**Note:** This aggregation is required for preprocessing but is *not* used for training the scDoRI model itself.

##### 1.3.2 Correlation Estimation

For each pseudobulk (metacell):

- ATAC-seq peak counts are library-size normalized and min-max scaled.

- TF expression from RNA-seq is normalized by library size and log-transformed.

- Pearson correlation is computed between TF expression and peak accessibility, resulting in a correlation matrix:

$$\mathbf{R}_{\text{peak-TF}} \in \mathbb{R}^{P \times F}.$$

##### 1.3.3 Motif Match Calculation

Motif match scores are computed using FIMO [6] against a reference genome, using a database of TF binding motifs. These scores are aggregated into a matrix:

$$\mathbf{M} \in \mathbb{R}^{P \times F},$$

where each entry reflects the motif match strength between a peak and a transcription factor.

###### 1.3.4 Filtering

To control for spurious correlations, we apply TF-specific significance thresholds using the motif scores:

- For each TF, we define a background set of peaks with the lowest motif scores (e.g., bottom 20%).
- A background distribution of correlations is constructed using this subset.
- We apply a TF-specific cutoff (e.g., 95th percentile for activators, 5th percentile for repressors) on the correlation matrix $\mathbf{R}_{\text{peak-TF}}$.
- Correlation values in $\mathbf{R}_{\text{peak-TF}}$ that do not pass this threshold are set to zero.

###### 1.3.5 Activators vs. Repressors

The filtered correlation matrix is then split into two matrices according to sign:

$$\mathbf{R}_{\text{act}}[i,j] = \max(\mathbf{R}_{\text{peak-TF}}[i,j], 0), \quad \mathbf{R}_{\text{rep}}[i,j] = \min(\mathbf{R}_{\text{peak-TF}}[i,j], 0).$$

###### 1.3.6 Integration of Motif and Correlation Signals

Finally, we compute the signed TF–peak interaction matrices by integrating the filtered correlation matrices with motif evidence:

$$\mathbf{W}^{\text{act}}_{\text{TF-peak}} = \mathbf{M} \odot \mathbf{R}_{\text{act}}, \quad \mathbf{W}^{\text{rep}}_{\text{TF-peak}} = \mathbf{M} \odot \mathbf{R}_{\text{rep}},$$

where $\odot$ denotes element-wise multiplication.

These matrices serve as inputs to the downstream GRN inference module, capturing both sequence-driven binding potential and regulatory directionality.

##### 1.4 scDoRI Model Architecture

The scDoRI model consists of an **Encoder** and a **Decoder**. The Encoder maps the input data into a shared latent topic space, while the Decoder reconstructs the observed data modalities and infers GRNs incorporating both activator and repressor components.

###### 1.4.1 Encoder

**Objective.** Map high-dimensional single-cell inputs (RNA + ATAC) into a lower-dimensional topic representation $\boldsymbol{\theta} \in \mathbb{R}^{C \times T}$, where $C$ corresponds to the number of cells (or metacells) and $T$ denotes a set number of topics to be inferred.

**Implementation.** We use two parallel neural networks (one for RNA, one for ATAC), each with two fully connected layers. Their outputs are concatenated into a combined vector $\mathbf{h}_{\text{combined}}$. A final linear layer $\mathbf{W}_\theta$ produces *topic logits*, which pass through a softmax to give rise to $\boldsymbol{\theta}$:

$$\begin{aligned}
\mathbf{h}_{\text{RNA}} &= \text{ReLU}\Big(\mathbf{W}_2^{\text{RNA}}\,\text{ReLU}\big(\mathbf{W}_1^{\text{RNA}}\,\mathbf{x}_{\text{RNA}} + \mathbf{b}_1^{\text{RNA}}\big) + \mathbf{b}_2^{\text{RNA}}\Big),\\
\mathbf{h}_{\text{ATAC}} &= \text{ReLU}\Big(\mathbf{W}_2^{\text{ATAC}}\,\text{ReLU}\big(\mathbf{W}_1^{\text{ATAC}}\,\mathbf{x}_{\text{ATAC}} + \mathbf{b}_1^{\text{ATAC}}\big) + \mathbf{b}_2^{\text{ATAC}}\Big),\\
\mathbf{h}_{\text{combined}} &= [\mathbf{h}_{\text{RNA}};\ \mathbf{h}_{\text{ATAC}}],\\
\boldsymbol{\mu}_\theta &= \mathbf{W}_\theta\,\mathbf{h}_{\text{combined}} + \mathbf{b}_\theta, \quad \boldsymbol{\theta} = \text{Softmax}(\boldsymbol{\mu}_\theta).
\end{aligned}$$

Here, $\mathbf{x}_{\text{RNA}}$ includes raw scRNA counts concatenated with RNA library size (sum of log-transformed RNA counts in a cell), while $\mathbf{x}_{\text{ATAC}}$ includes peak fragment counts concatenated with ATAC library size (sum of log-transformed fragment counts in a cell). This explicit inclusion of library size is intended to reduce the risk of the model encoding technical variation as biological signal in the latent space. Additionally, if the model is trained using metacells, the number of single cells constituting each metacell is also provided as input (set to 1 for single-cell data). The final softmax ensures that each cell's topic loadings sum to 1, paralleling the idea of a topic mixture in classical LDA [7].

###### 1.4.2 Decoder

The objective of the decoder is to reconstruct the observed data (both ATAC-seq and RNA-seq) *from the latent topic activity* $\boldsymbol{\theta}$. Unlike conventional autoencoders and other dimensionality reduction methods, scDoRI imposes biologically motivated constraints in the design of the decoders, thereby enabling the inference of eGRNs. We describe the design and biological priors encoded in the four reconstruction modules below (see also Supp. Fig. 2.1).

###### 1.4.3 Module 1: ATAC Reconstruction

**Objective: Objective:** Reconstruct single-cell ATAC-seq counts ($\mathbf{X}_{\text{ATAC}}$) from the latent topic activity ($\boldsymbol{\theta}$), with the goal of identifying sets of regulatory regions that tend to be co-accessible across cells. Such co-accessibility is expected to reflect coordinated regulatory activity and may correspond to underlying biological programs or cell states. This formulation is similar to **cisTopic** [8], where topics capture groups of chromatin regions that are frequently co-accessible and exhibit varying activity across cells, facilitating interpretable representations of chromatin accessibility landscapes.

**Parameters:**

- *Topic-Peak Linkage Matrix*, $\mathbf{W}_{\text{topic-peak}} \in \mathbb{R}^{T\times P}$. This matrix encodes the relationship between topics and chromatin peaks, where each row corresponds to a topic and each column corresponds to a peak. The matrix is learned during training to capture the patterns of co-accessible chromatin regions.
- *Batch correction weight*, $\mathbf{B}_{\text{ATAC}} \in \mathbb{R}^{K\times P}$, specifies batch-specific effects across ATAC peaks.

**Reconstruction Details:** We first compute a linear prediction for the ATAC-seq data as follows:

$$\mathbf{Z}_{\text{ATAC}} = \boldsymbol{\theta}\,\mathbf{W}_{\text{topic-peak}} \; + \; \mathbf{B}\,\mathbf{B}_{\text{ATAC}},$$

where $\boldsymbol{\theta}$ contains the topic activity for each cell, and $\mathbf{B}$ indicates the (one-hot) batch memberships. A batch normalization step is applied to $\mathbf{Z}_{\text{ATAC}}$, followed by the Softmax function:

$$\mathbf{R}_{\text{ATAC}} = \text{Softmax}\Big(\text{BatchNorm}\big(\mathbf{Z}_{\text{ATAC}}\big)\Big).$$

**Predicted Counts:** To obtain the final prediction of observed counts, we multiply by an exponential of the log-transformed library size $\mathbf{L}_{\text{ATAC}}$:

$$\hat{\mathbf{X}}_{\text{ATAC}} = \exp\big(\mathbf{L}_{\text{ATAC}}\big) \odot \mathbf{R}_{\text{ATAC}},$$

where $\odot$ denotes element-wise multiplication. This rescales the normalized distribution to account for variability in total sequencing depth across cells, producing $\hat{\mathbf{X}}_{\text{ATAC}}$ as the estimated per-cell ATAC-seq counts.

**Sparsity and Interpretability:** An $\ell_1$ penalty is imposed on $\mathbf{W}_{\text{topic-peak}}$ to encourage sparse loadings, which further facilitates the biological interpretation of the topics(refer 1.5).

###### 1.4.4 Module 2: RNA Reconstruction from ATAC

**Objective: Objective:** Reconstruct RNA expression levels ($\mathbf{X}_{\text{RNA}}$) based on the predicted chromatin accessibility landscape of a cell, captured through the decoder $\mathbf{W}_{\text{topic-peak}}$ in Module 1. The biological rationale is that open chromatin regions associated with a gene are predictive of that gene's expression.

**Parameters:**

- *Gene-Peak Linkage Matrix*, $\mathbf{W}_{\text{gene-peak}} \in \mathbb{R}^{G\times P}$. This matrix encodes which peaks regulate which genes, split into a static binary mask ($\mathbf{W}^{\text{mask}}_{\text{gene-peak}} \in \{0,1\}^{G\times P}$) and a learnable part ($\mathbf{W} \in [0,1]^{G\times P}$). We combine them element-wise:

  $$\mathbf{W}_{\text{gene-peak}} = \mathbf{W}^{\text{mask}}_{\text{gene-peak}} \odot \mathbf{W},$$

  The mask $\mathbf{W}^{\text{mask}}_{\text{gene-peak}}$ encodes a constraint that only peaks within a genomic window around a gene can contribute to that gene's expression. This window is a user-defined hyperparameter, set by default to 150 kbp around the gene body for the human genome.

  The learned gene-peak weights $\mathbf{W}$ are initialized with a distance-based exponential decay term, where the distance is the genomic distance between the peak and the gene. The decay rate parameter controls the rate at which the weight decays with distance, similar to how ArchR ([9]) calculates gene scores based on scATAC-seq data
- $\mathbf{B}_{\text{RNA}} \in \mathbb{R}^{K\times G}$ parameterises batch-specific effects for individual genes.

**Reconstruction Details:** We first scale $\mathbf{W}_{\text{topic-peak}}$ (the topic-to-peak weights learned from Module 1) by applying a Softmax, which ensures each row sums to 1, followed by a MinMax normalization across topics:

$$\tilde{\mathbf{W}}_{\text{topic-peak}} = \text{MinMaxNorm}\Big(\text{Softmax}\big(\mathbf{W}_{\text{topic-peak}}\big)\Big).$$

The resulting matrix $\tilde{\mathbf{W}}_{\text{topic-peak}}$ encodes each topic's relative preference for different peaks, with normalization applied to reduce sensitivity to outliers.
Next, raw gene activity scores ($\mathbf{R}_{\text{RNA}}$) are computed by multiplying the latent topic activity $\boldsymbol{\theta}$ against the scaled topic-peak matrix and then applying the gene-peak linkage:

$$\mathbf{R}_{\text{RNA}} = \boldsymbol{\theta}\,\tilde{\mathbf{W}}_{\text{topic-peak}}\,\mathbf{W}_{\text{gene-peak}}^{\top} \;+\; \mathbf{B}\,\mathbf{B}_{\text{RNA}}.$$

Here, $\boldsymbol{\theta}\,\tilde{\mathbf{W}}_{\text{topic-peak}}$ results in predicted accessibility profile for each cell, analogous to module 1, which is then mapped to genes via $\mathbf{W}_{\text{gene-peak}}^{\top}$. We add the offsets $\mathbf{B}_{\text{RNA}}$ to account for systematic variation across technical batches.
To ensure a proper probability-like distribution of relative gene activities, we apply batch normalization followed by a Softmax:

$$\mathbf{R}_{\text{RNA}} = \text{Softmax}\Big(\text{BatchNorm}\big(\mathbf{R}_{\text{RNA}}\big)\Big).$$

**Predicted Counts:** Finally, we multiply by a learned library size factor $\mathbf{L}_{\text{RNA}}^{\text{lib}} \in \mathbb{R}^{C\times 1}$, which is broadcasted across genes to perform element-wise scaling of the normalized output:

$$\hat{\mathbf{X}}_{\text{RNA}} = \mathbf{L}_{\text{RNA}}^{\text{lib}} \odot \mathbf{R}_{\text{RNA}},$$

where $\mathbf{R}_{\text{RNA}} \in \mathbb{R}^{C\times G}$. This step adjusts for varying sequencing depths across cells. The library size $\mathbf{L}_{\text{RNA}}^{\text{lib}}$ is learned from the data using a two-layer fully connected neural network applied to raw gene expression counts, similar to the approach used in scVI [2].

**Interpretation:** The matrix $\mathbf{W}_{\text{gene-peak}}$ captures topic-agnostic, enhancer–gene associations within a gene window. Since both the peak accessibility values and $\mathbf{W}_{\text{gene-peak}}$ are constrained to be non-negative, the resulting predictions $\hat{\mathbf{X}}_{\text{RNA}}$ primarily reflect the contribution of accessible chromatin to gene activation. That is, increased accessibility at linked peaks contributes additively to higher gene expression. This is similar to the enhancer-driven gene expression prediction logic used in prior frameworks like *SCENIC+*[10] or *Scarlink*[11], however integrated directly into our autoencoder-style framework. Note that scDoRI does not assume that all regulation is activating—rather, the mechanisms for capturing repressive TF effects are handled explicitly in Module 4, where signed TF–gene interactions are modeled using chromatin, motif, and co-expression cues.

##### 1.4.5 Module 3: TF Expression Reconstruction

**Objective:** To reconstruct TF expression such that each latent topic captures coherent groups of co-expressed TFs, reflecting upstream regulatory programs. This enables topics to encode not only patterns of chromatin accessibility and gene expression, but also their potential regulators. The learned *Topic–TF Decoder* yields a denoised, topic-level TF expression profile by smoothing over noisy single-cell measurements. This representation can serve as an input to Module 4 (Section 1.4.6), where it is used to reconstruct gene expression from topic-specific TF-gene links.

**Parameters:**

- *Topic–TF Decoder*, $\mathbf{W}_{\text{topic-TF}} \in \mathbb{R}^{T\times F}$. Each row corresponds to a topic and each column to a TF, encoding how strongly TF expression is associated with topic.
- *Batch correction weights*, $\mathbf{B}_{\text{TF}} \in \mathbb{R}^{K\times F}$. Parameterises batch-specific effects for individual TFs.

**Reconstruction Details:** We first compute a raw prediction $\mathbf{Z}_{\text{TF}}$ of TF expression as:

$$\mathbf{Z}_{\text{TF}} = \boldsymbol{\theta}\,\mathbf{W}_{\text{topic-TF}} \;+\; \mathbf{B}\,\mathbf{B}_{\text{TF}},$$

where $\boldsymbol{\theta} \in \mathbb{R}^{C\times T}$ represents per-cell topic activity and $\mathbf{B} \in \mathbb{R}^{C\times K}$ encodes batch identity.
We then normalize and convert to a compositional output:

$$\mathbf{R}_{\text{TF}} = \text{Softmax}\Big(\text{BatchNorm}\big(\mathbf{Z}_{\text{TF}}\big)\Big),$$

yielding a per-cell distribution over TFs.

**Predicted Counts:** Finally, we multiply by a learned library size factor $\mathbf{L}_{\text{TF}}^{\text{lib}} \in \mathbb{R}^{C\times 1}$ (similar to 1.4.4), which is broadcasted across TFs to perform element-wise scaling of the normalized output:

$$\hat{\mathbf{X}}_{\text{TF}} = \mathbf{L}_{\text{TF}}^{\text{lib}} \odot \mathbf{R}_{\text{TF}},$$

##### 1.4.6 Module 4: GRN Inference with Activators and Repressors

**Overview and Motivation.**
This final module estimates topic-specific eGRNs by integrating multiple signatures, either derived from previous modules or precomputed: in silico ChIP-seq TF binding (activator or repressor, 1.3), chromatin accesibility profiles(Module 1), enhancer–gene associations(Module 2), and TF–gene co-expression patterns(Module 3). Unlike previous approaches for eGRN inference that primarily on focus on activation signatures and rely on negative TF–gene correlation for repressors, **scDoRI** explicitly models the directionality (activation vs. repression) and context (topic specificity) of TF-driven regulation.

**Key Ideas and Rationale:**

- **Signed TF–Peak Binding** (*in silico* ChIP-seq): As described in Section 1.3, we encode how TFs open or close chromatin at a peak, indicating potential activator or repressor roles.
- **Enhancer–Gene Links (Module 2):** Not all TF-bound peaks matter for a particular gene. We use the $\mathbf{W}_{\text{gene-peak}}$ matrix to focus on peaks actually linked to each gene.
- **TF–Gene Co-Expression (Module 3):** Even with motif presence, the TF must be sufficiently expressed, and for activators we typically observe positive TF–gene correlation, whereas for repressors this correlation can be negative.
- **Topic Specificity (Modules 1 & 3):** Peak accessibility (open vs. closed) and TF expression vary by topic, allowing a TF to function as an activator in one topic but a repressor in another, depending on chromatin context and co-expression patterns.

**Biological Assumption for Repression.** Building on approaches like DiffTF and GRaNIE [4, 12], we treat negative TF–peak correlations (i.e., TF expression increases while accessibility decreases) as indicative of repressive activity. Although not all repressors close chromatin, this provides a tractable signature for capturing key repressive events.

**Workflow:**

1. **Topic–Peak Distributions** ($\mathbf{W}_{\text{topic-peak}}$, Module 1): Identifies which peaks are open or closed in each topic.
2. **TF–Peak Scores** ($\mathbf{W}^{\text{act/rep}}_{\text{TF-peak}}$): Captures whether a TF tends to open (activator) or close (repressor) chromatin at a given peak.
3. **Peak–Gene Matrix** ($\mathbf{W}_{\text{gene-peak}}$, Module 2): Determines which peaks regulate which genes.
4. **TF Expression** ($\hat{\mathbf{X}}_{\text{TF}}$, Module 3): Provides per-topic TF abundance, enforcing that TFs must be expressed to regulate genes.

**Combining Signals for Topic-Specific GRNs.** We integrate the various signatures from Modules 1–3 in a multi-step process to produce a final, signed, topic-aware GRN:

1. **Open vs. Closed Chromatin.** Convert $\mathbf{W}_{\text{topic-peak}}$ into a normalized distribution of peak accessibility per topic. For each topic $t$, we extract a vector highlighting open peaks ($\mathbf{v}^{\text{act}}_{\text{peak}}[t]$) and a reciprocal vector for closed peaks ($\mathbf{v}^{\text{rep}}_{\text{peak}}[t]$).
2. **Topic-Specific Peak–Gene Links.** From Module 2, $\mathbf{W}_{\text{gene-peak}}$ encodes which peaks regulate which genes in a generic manner. We now scale these links by the topic-specific accessibility state:
   - *Activator scenario*: Multiply $\mathbf{W}_{\text{gene-peak}}$ by $\mathbf{v}^{\text{act}}_{\text{peak}}[t]$ so that only peaks open in topic $t$ strongly contribute to activation.
   - *Repressor scenario*: Multiply $\mathbf{W}_{\text{gene-peak}}$ by $\mathbf{v}^{\text{rep}}_{\text{peak}}[t]$, which emphasizes peaks that are relatively closed in topic $t$. These can mediate repressive regulation if bound by a repressor TF.

   This step allows the same peak–gene link to act as an enhancer in one topic (open) and as a potential silencer or as being inactive in another topic (closed).
3. **TF–Peak $\times$ Peak–Gene $\to$ Candidate TF–Gene.** We multiply the signed TF–peak matrices $\mathbf{W}^{\text{act}}_{\text{TF-peak}}$ and $\mathbf{W}^{\text{rep}}_{\text{TF-peak}}$ by the corresponding topic-scaled peak–gene link matrices from step (2). This yields a raw set of candidate TF–gene relationships for each topic, denoted as $\mathbf{G}^{\text{act}}_{\text{ATAC}}[t]$ and $\mathbf{G}^{\text{rep}}_{\text{ATAC}}[t]$, capturing whether a TF bound to certain peaks can influence specific genes via those peaks.

4. **Refine by TF Expression.** The raw TF–gene potential is then modulated by $\mathbf{G}^{\text{act/rep}}_{\text{TF-gene-topic}}$, ensuring that the TF must be sufficiently expressed and that its expression correlates appropriately with the target gene. Positive expression correlation reinforces activator logic, while negative correlation boosts repressor logic.

5. **Assemble Signed GRN.** We combine the activator and repressor potentials into a TF-gene matrix $\mathbf{G}_{\text{net}}[t]$ for each topic $t$:
$$\mathbf{G}_{\text{net}}[t] = \mathbf{G}^{\text{act}}_{\text{combined}}[t] \; - \; \mathbf{G}^{\text{rep}}_{\text{combined}}[t].$$
Positive entries indicate net activation; negative entries indicate net repression. This is the final eGRN we are interested in.

6. **Predict RNA Expression.** Finally, we multiply $\mathbf{G}_{\text{net}}[t]$ by the topic-level TF expression $\hat{\mathbf{X}}_{\text{TF}}[t,:]$ and sum across topics using the cell's topic mixture $\boldsymbol{\theta}$ to produce a gene-expression reconstruction. A negative binomial loss aligns the model's predictions with observed single-cell data, completing the end-to-end training of the GRN inference module.

**Equations and Implementation Details.**

$$\begin{aligned}
\tilde{\mathbf{W}}_{\text{topic-peak}} &= \text{MinMaxNorm}\big(\text{Softmax}(\mathbf{W}_{\text{topic-peak}})\big),\\
\mathbf{v}^{\text{act}}_{\text{peak}}[t] &= \tilde{\mathbf{W}}_{\text{topic-peak}}[t,:], \quad \mathbf{v}^{\text{rep}}_{\text{peak}}[t] = \frac{1}{\tilde{\mathbf{W}}_{\text{topic-peak}}[t,:]+\epsilon},\\
\mathbf{G}^{\text{act}}_{\text{ATAC}}[t] &= \big(\mathbf{v}^{\text{act}}_{\text{peak}}[t] \odot \mathbf{W}^{\text{act}}_{\text{TF-peak}}\big)^{\top}\mathbf{W}^{\top}_{\text{gene-peak}}, \quad \mathbf{G}^{\text{rep}}_{\text{ATAC}}[t] = \big(\mathbf{v}^{\text{rep}}_{\text{peak}}[t] \odot \mathbf{W}^{\text{rep}}_{\text{TF-peak}}\big)^{\top}\mathbf{W}^{\top}_{\text{gene-peak}},\\
\mathbf{G}^{\text{act}}_{\text{combined}}[t] &= \mathbf{G}^{\text{act}}_{\text{ATAC}}[t] \; \odot \; \text{ReLU}\big(\mathbf{G}^{\text{act}}_{\text{TF-gene-topic}}[t]\big),\\
\mathbf{G}^{\text{rep}}_{\text{combined}}[t] &= \mathbf{G}^{\text{rep}}_{\text{ATAC}}[t] \; \odot \; \text{ReLU}\big(\mathbf{G}^{\text{rep}}_{\text{TF-gene-topic}}[t]\big),\\
\mathbf{G}_{\text{net}}[t] &= \mathbf{G}^{\text{act}}_{\text{combined}}[t] \; - \; \mathbf{G}^{\text{rep}}_{\text{combined}}[t],\\
\mathbf{C}[t,:] &= \hat{\mathbf{X}}_{\text{TF}}[t,:] \; \mathbf{G}_{\text{net}}[t],\\
\mathbf{R}_{\text{RNA-GRN}} &= \boldsymbol{\theta}\,\mathbf{C} + \mathbf{B}\,\mathbf{B}_{\text{RNA-GRN}},\\
\hat{\mathbf{X}}_{\text{RNA-GRN}} &= \mathbf{L}^{\text{lib}}_{\text{RNA}} \; \odot \; \text{Softmax}\Big(\text{BatchNorm}(\mathbf{R}_{\text{RNA-GRN}})\Big).
\end{aligned}$$

**Interpretation and Practical Notes:**

- **Context-Dependent Signed Regulation** Because chromatin openness and TF expression vary, one TF may be an activator in topic $A$ but a repressor in topic $B$ and can have different set of target genes between topics.
- **Biological Plausibility:** By requiring both chromatin-level evidence (TF opens or closes a peak) and expression-level evidence (TF and gene co-variation), the model yields more mechanistically informed GRNs.

#### 1.5 Loss Functions

scDoRI employs several data reconstruction and regularization losses to balance accurate reconstruction with interpretable parameters. Specifically, we model ATAC-seq fragment counts using a Poisson likelihood and RNA-seq using a negative binomial (NB) likelihood, which better captures the mean–variance relationship typical of scRNA-seq data, following practices established in **scVI** [2]. In all cases, the model predicts the mean of the respective distribution; for NB likelihoods, gene- or TF-specific overdispersion parameters are learned from the data.

##### 1.5.1 ATAC Reconstruction Loss

$$\mathcal{L}_{\text{ATAC}} = -\sum_{c=1}^{C}\sum_{p=1}^{P}\Big(\mathbf{X}_{\text{ATAC}}[c,p]\,\log\big(\hat{\mathbf{X}}_{\text{ATAC}}[c,p]\big) - \hat{\mathbf{X}}_{\text{ATAC}}[c,p]\Big).$$

We treat the observed ATAC fragment counts under a Poisson likelihood, following recommendations from [13]. The model predicts the Poisson mean $\hat{\mathbf{X}}_{\text{ATAC}}$ for each cell and peak.

##### 1.5.2 RNA Reconstruction Losses

- **RNA-from-ATAC Loss ($\mathcal{L}_{\textbf{RNA-ATAC}}$):**

$$\mathcal{L}_{\text{RNA-ATAC}} = -\sum_{c=1}^{C}\sum_{g=1}^{G} \log \text{NB}\Big(\mathbf{X}_{\text{RNA}}[c,g]; \hat{\mathbf{X}}_{\text{RNA}}[c,g], \alpha_{\text{RNA}}[g]\Big),$$

where $\text{NB}(x;\mu,\alpha)$ denotes the negative binomial likelihood with mean $\mu$ and overdispersion $\alpha$. This term enforces that accessibility of nearby peaks (modulated by topic activity) predicts gene expression. The model outputs the mean $\hat{\mathbf{X}}_{\text{RNA}}$, while the overdispersion $\alpha_{\text{RNA}}[g]$ is a **gene-specific parameter** learned from data.

- **RNA-GRN Loss ($\mathcal{L}_{\textbf{RNA-GRN}}$):**

$$\mathcal{L}_{\text{RNA-GRN}} = -\sum_{c=1}^{C}\sum_{g=1}^{G} \log \text{NB}\Big(\mathbf{X}_{\text{RNA}}[c,g]; \hat{\mathbf{X}}_{\text{RNA-GRN}}[c,g], \alpha_{\text{RNA}}[g]\Big).$$

Here, $\hat{\mathbf{X}}_{\text{RNA-GRN}}$ is predicted based on TF activity and signed TF–gene GRNs. Minimizing this loss encourages accurate modeling of transcriptional regulation driven by TF binding profiles. As above, $\alpha_{\text{RNA}}[g]$ is learned per gene.

##### 1.5.3 TF Expression Reconstruction Loss

$$\mathcal{L}_{\text{TF}} = -\sum_{c=1}^{C}\sum_{f=1}^{F} \log \text{NB}\Big(\mathbf{X}_{\text{TF}}[c,f]; \hat{\mathbf{X}}_{\text{TF}}[c,f], \alpha_{\text{TF}}[f]\Big).$$

This loss models TF expression using an NB likelihood. The model predicts the mean $\hat{\mathbf{X}}_{\text{TF}}$ and learns a **TF-specific overdispersion parameter** $\alpha_{\text{TF}}[f]$ from data.

##### 1.5.4 Regularization Terms

- **L1 on GRN Weights:** To promote sparsity in TF–gene edges, we use:

$$\mathcal{L}_{\text{reg}}^{\text{act}} = \lambda_{\text{act}} \big\|\mathbf{G}_{\text{TF-gene-topic}}^{\text{act}}\big\|_1, \quad \mathcal{L}_{\text{reg}}^{\text{rep}} = \lambda_{\text{rep}} \big\|\mathbf{G}_{\text{TF-gene-topic}}^{\text{rep}}\big\|_1.$$

- **L1 on Decoder Weights ($\mathbf{W}_{\textbf{topic-peak}}$):**

$$\mathcal{L}_{\text{reg}}^{L1} = \lambda_1 \big\|\mathbf{W}_{\text{topic-peak}}\big\|_1.$$

- **L2 on Gene-Peak Weights ($\mathbf{W}_{\textbf{gene-peak}}^{\textbf{learned}}$):**

$$\mathcal{L}_{\text{reg}}^{L2} = \lambda_2 \big\|\mathbf{W}_{\text{gene-peak}}^{\text{learned}}\big\|_2.$$

- **Total Regularization:**

$$\mathcal{L}_{\text{reg}} = \mathcal{L}_{\text{reg}}^{\text{act}} + \mathcal{L}_{\text{reg}}^{\text{rep}} + \mathcal{L}_{\text{reg}}^{L1} + \mathcal{L}_{\text{reg}}^{L2}.$$

#### 1.6 Training Procedure: Two-Phase Optimization Approach

Like a conventional autoencoder, **scDoRI** is trained using reconstruction-based objectives for each modality. However, because the data are multi-modal and **scDoRI** integrates multiple, mutually dependent decoders (Modules 1–4), we adopt a *two-phase optimization strategy* to ensure stable training. Since Module 4 depends on the latent topics and representations learned by Modules 1–3, we first train these core components independently. In the second phase, we introduce the signed GRN module (Module 4) and fine-tune the model to capture TF-mediated regulatory effects, including activation and repression. The key steps, training objectives, and implementation details of each phase are outlined below.

##### 1.6.1 Phase 1: Core Reconstruction (Modules 1–3)

**Goals and Overview.** In this initial phase, the model is trained using only Modules 1–3. The objective is to learn a stable and informative latent topic space $\boldsymbol{\theta}$, while capturing the core relationships across chromatin accessibility (ATAC), gene expression (RNA), and transcription factor (TF) activity. This forms the foundation for accurate multi-omic integration and prepares the model for the introduction of regulatory logic in the next phase.

**Loss Function for Phase 1.** We define a composite objective, $\mathcal{L}_{\text{phase1}}$, that combines reconstruction losses across three modalities—ATAC, RNA, and TF expression—along with regularization terms:

$$\mathcal{L}_{\text{phase1}} = \beta_{\text{ATAC}}\,\mathcal{L}_{\text{ATAC}} + \beta_{\text{RNA-ATAC}}\,\mathcal{L}_{\text{RNA-ATAC}} + \beta_{\text{TF}}\,\mathcal{L}_{\text{TF}} + \lambda_1\|\mathbf{W}_{\text{topic-peak}}\|_1 + \lambda_2\|\mathbf{W}^{\text{learned}}_{\text{gene-peak}}\|_2^2.$$

- $\mathcal{L}_{\text{ATAC}}, \mathcal{L}_{\text{RNA-ATAC}}, \mathcal{L}_{\text{TF}}$: Negative log-likelihoods based on Poisson (ATAC) or negative binomial (RNA and TF) distributions, measuring reconstruction error for each modality in Modules 1–3.
- $\lambda_1, \lambda_2$: Regularization coefficients that encourage sparsity in topic–peak mappings and learned gene–peak associations.
- **Loss Weighting:** In practice, the weights $\beta_{\text{ATAC}}, \beta_{\text{RNA-ATAC}}, \beta_{\text{TF}}$ are tuned to place the reconstruction losses on comparable numerical scales.

All parameters in the encoder and Modules 1–3 are optimized jointly, including modality-specific batch offsets ($\mathbf{B}_{\text{ATAC}}, \mathbf{B}_{\text{RNA}}, \mathbf{B}_{\text{TF}}$).

##### 1.6.2 Phase 2: eGRN inference (Module 4)

**Goals and Overview.** In Phase 2, we introduce Module 4, which models TF–gene interactions using inferred activator and repressor relationships. This module integrates in silico ChIP-seq signals, peak–gene links, topic-level chromatin accessibility and TF expression to capture topic-specific regulation.

A user-defined choice in this phase is whether to update the encoder and Modules 1–3:

- **Freezing:** Keeps the latent topics and upstream modules fixed as learned in Phase 1, focusing optimization solely on the newly introduced GRN parameters in Module 4. These include the activator and repressor TF–gene–topic tensors ($\mathbf{G}^{\text{act/rep}}_{\text{TF-gene-topic}}$), as well as any additional GRN-specific batch offsets. Freezing preserves interpretability, speeds up training, and avoids destabilizing previously learned representations.
- **Fine-Tuning:** Allows the entire model—including the encoder and Modules 1–3—to adapt in response to the newly introduced regulatory structure. This can improve GRN specificity by aligning the latent topic space with regulatory variation but may reduce interpretability.

The choice depends on dataset size, computational resources, and user goals. Freezing earlier modules can significantly speed up training and is often preferred for interpretability or when working with large datasets under limited compute. In contrast, fine-tuning the full model may improve the specificity and accuracy of the inferred GRNs by allowing the latent space to adapt to regulatory signals.

**Note:** In the glioblastoma application presented in this study, we freeze the encoder and Modules 1–3 during Phase 2 training to maintain interpretability and reduce training time.

**Optional Input: Empirical vs. Learned TF Expression.** By default, Module 4 uses *empirical* topic-level TF expression, computed by averaging raw TF counts across cells, weighted by topic loadings $\boldsymbol{\theta}$. This reflects the true measured TF expression and may provide stronger biological grounding when expression measurements are reliable and well-powered.

Alternatively, users may use the *learned* topic-level TF expression inferred by Module 3 (via $\mathbf{W}_{\text{topic-TF}}$). This representation denoises TF counts across cells and captures co-expression structure, offering robustness in noisy, sparse, or low-coverage datasets.

**Note:** In the glioblastoma application presented in this study, we use the empirical (true) TF expression as input to Module 4.

**Phase 2 Loss Function.** We extend the training objective with a GRN-specific term, $\mathcal{L}_{\text{RNA-GRN}}$, which models RNA reconstruction based on signed TF–gene interactions:

$$\mathcal{L}_{\text{phase2}} = \beta_{\text{ATAC}}\,\mathcal{L}_{\text{ATAC}} + \beta_{\text{RNA}}\big(\mathcal{L}_{\text{RNA-ATAC}} + \mathcal{L}_{\text{RNA-GRN}}\big) + \beta_{\text{TF}}\,\mathcal{L}_{\text{TF}} + \mathcal{L}_{\text{reg}}.$$

Here, $\mathcal{L}_{\text{RNA-GRN}}$ is a negative binomial loss assessing how well the GRN-based RNA prediction (Module 4) matches observed expression. $\mathcal{L}_{\text{reg}}$ includes optional L1 penalties on the activator and repressor tensors $\mathbf{G}^{\text{act/rep}}_{\text{TF-gene-topic}}$, promoting sparsity and interpretability in the inferred networks.

If Modules 1–3 are frozen during Phase 2, the losses $\mathcal{L}_{\text{ATAC}}, \mathcal{L}_{\text{RNA-ATAC}}, \mathcal{L}_{\text{TF}}$ are excluded from the objective. This ensures that optimization focuses solely on refining the GRN without altering the latent topic structure or upstream modality reconstructions.

##### 1.6.3 Optimization Details and Constraints

Across both training phases, we apply several practical strategies to ensure stability, convergence, and interpretability:

- **Optimizer:** We use the Adam optimizer[14] with an appropriate learning rate $\eta$. See Section 2.3 for hyperparameters specific to the scDoRI GB atlas experiments.
- **Early Stopping:** Training is halted if the validation loss does not improve for a fixed number of epochs, reducing the risk of overfitting and unnecessary computation.
- **Constraints:**
  - **Signed TF–gene edges:** Non-negativity constraints are applied to $\mathbf{G}^{\text{act}}_{\text{TF-gene-topic}}$ and $\mathbf{G}^{\text{rep}}_{\text{TF-gene-topic}}$, ensuring consistent interpretation as activators and repressors, respectively.
  - **Gene–peak link weights:** The matrix $\mathbf{W}^{\text{learned}}_{\text{gene-peak}}$ is clamped to the interval $[0, 1]$ to enhance interpretability. A value of 0 indicates no influence of a peak on the gene, while 1 indicates maximal linkage.

#### 1.7 Summary of Model Components

As summarized in Table 2, the scDoRI model comprises several interconnected components, each designed to capture different aspects of the regulatory landscape.

#### 1.8 Software Usage Guidelines

We provide here an overview of best practices and design choices for configuring and executing the scDoRI framework. This includes recommendations for input preparation, hyperparameter tuning, and key considerations when training on large-scale single-cell multi-omic datasets.

###### Feature Selection and Resource Considerations

scDoRI operates on a defined subset of peaks, genes, and transcription factors (TFs), typically selected based on variability or prior biological interest. To ensure feasible training times and memory efficiency, users should select input features appropriate to the available computational resources. For instance, GPUs with 12–15 GB memory typically support 4000 genes, 70,000 peaks and 300 TFs. Higher memory availability allows for more fine-grained models.

Incorporating prior knowledge is encouraged: users may include known marker genes or candidate regulators to ensure their presence in the final model. For TFs, we recommend ensuring motif annotations are available in the supplied database for proper integration into regulatory modeling.

###### Motif Scanning and TF Annotation

scDoRI leverages motif scanning to estimate TF binding potential across chromatin peaks, requiring a position weight matrix (PWM) database in MEME format. Users may use curated collections such as cisBP[15], JASPAR [16] or Hocomoco [17], or substitute custom motif files as long as they maintain this format. The stringency of motif matching influences the specificity of the inferred TF–peak connections and should be chosen based on the desired confidence level.

Table 1: Overview of scDoRI model components. Each module captures a distinct aspect of regulatory inference, ranging from raw accessibility and gene expression reconstruction (Modules 1–3) to the final context-aware inference of signed TF–gene regulatory networks (Module 4). The table details inputs, outputs, loss functions, sparsity constraints, and biological utility of each component. We ignore batch correction terms for conciseness.

Table 2: Summary of scDoRI Model Components

| **Component** | **Major Inputs** | **Outputs** | **Loss Function** | **Sparsity / Other Constraints** | **Downstream Application** |
|---|---|---|---|---|---|
| **Encoder** | • scRNA-seq counts $\mathbf{X}_{\text{RNA}}$<br>• scATAC-seq counts $\mathbf{X}_{\text{ATAC}}$<br>• Library sizes $\mathbf{L}$ (log-transformed)<br>• Number of cells per metacell $\mathbf{n}$ (set to 1 for single-cell data) | Latent topic distribution $\boldsymbol{\theta}$ for each cell | NA | • Softmax normalization on latent space | • Cell state representation<br>• Dimensionality reduction<br>• Integration of multi-omic data |
| **Decoder Module 1: ATAC Reconstruction** | • Latent topics $\boldsymbol{\theta}$<br>• Topic-Peak decoder weights $\mathbf{W}_{\text{topic-peak}}$ | Reconstructed scATAC-seq fragment counts $\hat{\mathbf{X}}_{\text{ATAC}}$ | Poisson loss $\mathcal{L}_{\text{ATAC}}$ | • L1 regularization on $\mathbf{W}_{\text{topic-peak}}$ | • Capturing co-accessibility patterns of peaks<br>• Inferring topic-specific accessible regions<br>• Enhancer identification |
| **Decoder Module 2: RNA Reconstruction from ATAC** | • Latent topics $\boldsymbol{\theta}$<br>• Topic-Peak decoder weights $\mathbf{W}_{\text{topic-peak}}$<br>• Peak-Gene linkage matrix $\mathbf{W}_{\text{peak-gene}}$ | Predicted RNA expression counts $\hat{\mathbf{X}}_{\text{RNA-ATAC}}$ | Negative binomial reconstruction loss $\mathcal{L}_{\text{RNA-ATAC}}$ | • L2 regularization on $\mathbf{W}_{\text{peak-gene}}$<br>• Clamped to $[0, 1]$ for interpretability | • Inferring enhancer-gene links<br>• Deciphering cis-regulatory mechanisms |
| **Decoder Module 3: TF Expression Reconstruction** | • Latent topics $\boldsymbol{\theta}$<br>• Topic-TF decoder weights $\mathbf{W}_{\text{topic-TF}}$ | Predicted TF expression counts $\hat{\mathbf{X}}_{\text{TF}}$ | Negative binomial reconstruction loss $\mathcal{L}_{\text{TF}}$ | • None by default but optionally L1 regularization on $\mathbf{W}_{\text{topic-TF}}$ | • Denoised expression profiles of TFs |
| **Decoder Module 4: GRN Inference with Activators and Repressors** | • Latent topics $\boldsymbol{\theta}$<br>• Composite $\text{GRN}_{\text{ATAC}}$ signal (derived from topic–peak, peak–gene, and signed TF–peak matrices)<br>• Topic–TF matrix $\mathbf{W}_{\text{topic-TF}}$<br>• Learnable GRN weights: $\mathbf{G}^{\text{act}}_{\text{TF-gene-topic}}$, $\mathbf{G}^{\text{rep}}_{\text{TF-gene-topic}}$<br>• RNA library size $\mathbf{L}_{\text{RNA}}$ | • Predicted RNA expression from GRN $\hat{\mathbf{X}}_{\text{RNA-GRN}}$ | • Negative binomial reconstruction loss $\mathcal{L}_{\text{RNA-GRN}}$ | • L1 penalty and non-negativity constraints on $\mathbf{G}^{\text{act/rep}}_{\text{TF-gene-topic}}$ | • Inferring topic-specific TF-gene regulatory links<br>• Enabling cell-specific GRN estimation |

##### Metacell Construction for Robust Correlation Estimates for Insilico-chipseq

Metacells are constructed by clustering cells in a low-dimensional RNA space after batch correction, typically using Leiden community detection. These aggregated units reduce sparsity and improve the accuracy of TF–peak correlation estimates. We recommend generating at least 50 metacells for robust downstream analyses. If fewer clusters are obtained, the resolution parameter in the clustering step can be increased accordingly.

##### Peak–Gene Linking Window

Enhancers are linked to their target genes using a fixed genomic distance threshold, typically 150 kb upstream and downstream of gene bodies. This window reflects a trade-off between capturing distal regulatory elements and minimizing spurious associations. Users may adjust this parameter depending on species-specific genome architecture or desired sensitivity.

##### Topic Number

The number of topics should ideally exceed the number of expected cell types to allow the model to capture finer regulatory programs. A good starting point for number of topics is 1.5 * number of expected celltypes. Topics with little data support will be automatically downweighted. Each topic has its own regulatory network, and thus more topics also increase model complexity.

##### Training Regime and Epoch Configuration

Training proceeds in two phases (Section 1.6): an initial phase that reconstructs RNA and ATAC independently of TF regulation, followed by a fine-tuning phase that introduces GRN modeling. The number of training epochs can be set by estimating the total number of gradient updates (60,000 steps is a good default start) and dividing by the number of batches per epoch (number of cells in dataset / minbatch size). Early stopping is employed based on validation loss with a patience of approximately 5% of the total epochs.

##### Usage

scDoRI is implemented in Pytorch [18] and supports GPU acceleration. The pipeline can be executed end-to-end using configuration files that define input preprocessing, model architecture, and training parameters.

#### 1.9 Downstream Analysis enabled by scDoRI

This section details major downstream tasks after training the scDoRI model, including:

1. Visualizing the **scDoRI latent space** and aggregating topic usage across cell-states,
2. Analyzing the **topic-peak** matrix (for inspecting co-accesibility patterns of enhancers),
3. Computing a **topic-gene** matrix (using Module 2) and performing gene set enrichment (GSEA),
4. Inspecting the **peak-gene** matrix for enhancer–gene links,
5. Building **Topic-specific GRNs** with empirical permutation-based significance,
6. Identifying **top activators or repressors** per topic (these serve as TRs as used in TAP analysis (Figure 3, 2.5)),
7. Computing **TF activity** at both the cell and topic levels.
8. Downstream targets of **TFs** across topics.

Detailed code snippets for these analysis are provided in the tutorial notebooks included with the scDoRI package.

##### 1.9.1 scDoRI Latent Space

Each cell (or metacell) $c$ is assigned a topic distribution,

$$\boldsymbol{\theta}[c,:] \;=\; \big(\theta[c,1],\ldots,\theta[c,T]\big), \quad \sum_{t=1}^{T}\theta[c,t]=1,$$

arising from scDoRI's **Encoder** (Section 1.4.1). These distributions can be used to:

- **Visualize cells** in 2D via dimensionality reduction (e.g. UMAP) on $\boldsymbol{\theta}$. Cells with similar topic usage cluster together, indicating related regulatory states.
- **Visualize latent topic activity** in single cells.
- **Aggregate $\boldsymbol{\theta}$** across known groups (e.g. cell types) by mean or median to form a (Groups $\times$ $T$) table. One may min–max scale these values to highlight the dominant topics in each group.

##### 1.9.2 Topic-Peak Matrix: Co-Accessibility Patterns

In **module 1**, scDoRI learns **Topic-Peak Decoder**, $\mathbf{W}_{\text{topic-peak}} \in \mathbb{R}^{T \times P}$, which represents the contribution of each peak $p$ to each topic $t$. This can be used to:

- Identify sets of enhancers that tend to be active together in specific cellular contexts (e.g., states or lineages).
- Visualize these patterns by projecting peaks into a low-dimensional space using UMAP, where clusters of co-accessible peaks naturally emerge.
- Compare scDoRI-derived groupings with ground-truth chromatin accessibility to validate biological coherence.

To perform these analyses, we post-process the topic-peak decoder weights using the following steps:

1. **Normalize across peaks per topic** using softmax:

$$\mathbf{W}^{(\text{smx})}_{\text{topic-peak}}[t,p] \;=\; \frac{\exp(\mathbf{W}_{\text{topic-peak}}[t,p])}{\sum_{p'=1}^{P}\exp(\mathbf{W}_{\text{topic-peak}}[t,p'])}.$$

2. **Min–max scale across topics for each peak** to highlight specificity.
3. **Filter out ubiquitously accessible peaks** that are non-informative across topics.
4. **Construct a $P \times T$ matrix** using the processed weights and apply UMAP to reveal clusters of peaks with similar topic usage.

##### 1.9.3 Topic-Gene Matrix (Module 2) & Gene Set Enrichment

scDoRI facilitates the interpretation of topics by linking them to downstream gene programs. The **Topic-Gene Matrix** enables downstream identification of topic-specific biological functions through gene set enrichment analysis (GSEA).

**3.1 Constructing the Topic-Gene Matrix.** The topic-gene matrix is computed by combining topic-specific peak weights with enhancer–gene links:

$$\mathbf{M}_{\text{topic-gene}} = \tilde{\mathbf{W}}_{\text{topic-peak}} \cdot \mathbf{W}^{\top}_{\text{gene-peak}},$$

where $\tilde{\mathbf{W}}_{\text{topic-peak}} \in \mathbb{R}^{T \times P}$ is a scaled version of the topic-peak decoder, and $\mathbf{W}_{\text{gene-peak}} \in \mathbb{R}^{G \times P}$ encodes enhancer–gene associations.

Each entry $\mathbf{M}_{\text{topic-gene}}[t,g]$ reflects the inferred influence of gene $g$ in topic $t$. This matrix provides a ranked list of genes per topic, which can be used to assess enrichment of gene sets in that topic.

##### 1.9.4 Peak-Gene Matrix (Module 2): Enhancer–Gene Links

The matrix $\mathbf{W}_{\text{gene-peak}} = \mathbf{W}^{\text{mask}}_{\text{gene-peak}} \odot \mathbf{W}$ represents scDoRI's learnt enhancer–gene connectivity. Inspecting $\mathbf{W}_{\text{gene-peak}}[g,p]$ can confirm existing or discover novel enhancer–gene relationships.

- **Thresholding**: For instance, only keep $(g,p)$ entries if $\mathbf{W}_{\text{gene-peak}}[g,p] > 0.2$.
- **Topic-specific peak-gene links**: Multiply $\mathbf{W}_{\text{gene-peak}}$ by $\tilde{\mathbf{W}}_{\text{topic-peak}}$ to see how strongly gene $g$ is associated with each topic's set of peaks.

##### 1.9.5 Empirical thresholding of eGRNs

While scDoRI infers enhancer-driven GRNs by integrating chromatin accessibility, TF-binding motifs, enhancer–gene links, and co-expression (Modules 1–4), some TF–gene links may arise spuriously due to noisy inputs or model flexibility. To reduce such artifacts, we apply a post hoc empirical filtering step to the ATAC-derived GRN component, which can be recomputed independently of model training.

This filtering is biologically motivated: although TFs may show potential regulatory influence in the model, we aim to retain only those links where the chromatin signal (TF binding to regulatory regions of a gene) exceeds what would be expected by chance. This helps distinguish strong, biologically plausible interactions from random or low-confidence associations. Ideally, one would train the entire model multiple times using permuted TF–peak binding profiles to generate a full null distribution for the GRN weights. However, this is computationally infeasible at scale. Instead, we decouple filtering from training and focus only on the chromatin-based component, which can be efficiently recomputed post hoc.

To estimate a background, we generate permuted versions of the TF–peak binding matrix ($\mathbf{W}^{\text{act/rep}}_{\text{TF-peak}}$) and recompute the resulting GRNs $\mathbf{G}^{\text{act/rep}}_{\text{ATAC}}[t]$ for each permutation. For every TF–gene–topic triplet $(t, f, g)$, we retain the original score only if it exceeds an empirical threshold based on the null distribution—e.g., above the 95th percentile for activators or below the 5th percentile for repressors. This yields filtered GRNs:

$$\mathbf{G}^{\text{act, filt}}_{\text{ATAC}}[t], \quad \mathbf{G}^{\text{rep, filt}}_{\text{ATAC}}[t]$$

##### 1.9.6 Final GRN Construction

We combine the filtered chromatin-based GRNs with the expression-informed GRNs learned in Module 4 using element-wise multiplication:

$$\mathbf{G}^{\text{act}}_{\text{final}}[t] = \mathbf{G}^{\text{act, filt}}_{\text{ATAC}}[t] \odot \mathbf{G}^{\text{act}}_{\text{TF-gene-topic}}[t]$$
$$\mathbf{G}^{\text{rep}}_{\text{final}}[t] = \mathbf{G}^{\text{rep, filt}}_{\text{ATAC}}[t] \odot \mathbf{G}^{\text{rep}}_{\text{TF-gene-topic}}[t]$$

**Interpretation.** This final GRN contains topic-specific TF–gene links that are:

- Supported by chromatin-based regulatory evidence that exceeds background noise.
- Consistent with the expression-based regulatory influence of TFs on gene targets.

Together, this yields an interpretable GRN that combines multi-omic signals while filtering for confident regulatory edges.

##### 1.9.7 TF Activity Scores and Topic Regulators (TRs)

**Motivation:** The final signed GRNs $\mathbf{G}^{\text{act/rep}}_{\text{final}}[t] \in \mathbb{R}^{F \times G}$ summarize the influence of each TF on each gene in each topic. However, to interpret this at a systems level, we extract higher-level summaries: (i) which TFs are most active in each topic (Topic regulators,TRs), and (ii) how active each TF is in individual cells.

**8.1 Topic-Level TF Activity.** We compute per-topic TF activity scores by aggregating the regulatory strength of each TF across all target genes, separately for activators and repressors:

$$\mathbf{A}^{\text{act}}[t, f] = \frac{\sum_g \mathbf{G}^{\text{act}}_{\text{final}}[t](f, g)}{\sum_{f', g} \mathbf{G}^{\text{act}}_{\text{final}}[t](f', g) + \epsilon}, \quad \mathbf{A}^{\text{rep}}[t, f] = \frac{\sum_g \mathbf{G}^{\text{rep}}_{\text{final}}[t](f, g)}{\sum_{f', g} \mathbf{G}^{\text{rep}}_{\text{final}}[t](f', g) + \epsilon}.$$

These scores reflect each TF's relative contribution to activation or repression of gene expression in topic $t$.

**8.2 Topic Regulators (TRs).** We define **Topic Regulators (TRs)** as TFs with high and topic-specific activity, computed separately for activators and repressors. For a TF $f$ and topic $t$, we define its composite activity score:

$$\text{ACT}^{\text{act}}(f, t) = \text{Avg}\left(\text{MinMax}_{f'}\left(\mathbf{A}^{\text{act}}[t, f']\right), \text{MinMax}_{t'}\left(\mathbf{A}^{\text{act}}[t', f]\right)\right)$$

$$\text{ACT}^{\text{rep}}(f, t) = \text{Avg}\left(\text{MinMax}_{f'}\left(\mathbf{A}^{\text{rep}}[t, f']\right), \text{MinMax}_{t'}\left(\mathbf{A}^{\text{rep}}[t', f]\right)\right)$$

These scores measure both:

- **Intensity:** The strength of TF $f$'s activity within topic $t$, normalized across TFs.
- **Specificity:** How uniquely TF $f$ is active in topic $t$ compared to other topics.

Top-ranked TFs by $\text{ACT}^{\text{act}}(f, t)$ or $\text{ACT}^{\text{rep}}(f, t)$ are reported as the primary activator or repressor regulators of topic $t$.

**8.3 Cell-Level TF Activity.** To score TF activity per cell, we use the cell's topic weights $\boldsymbol{\theta}[c,:]$ to project topic-level activity down to the cell level:

$$\mathbf{Activity}_{\text{cell}}^{\text{act}}[c,f] = \sum_{t=1}^{T} \boldsymbol{\theta}[c,t] \cdot \mathbf{A}^{\text{act}}[t,f], \quad \mathbf{Activity}_{\text{cell}}^{\text{rep}}[c,f] = \sum_{t=1}^{T} \boldsymbol{\theta}[c,t] \cdot \mathbf{A}^{\text{rep}}[t,f]$$

These values can be $z$-scored across cells and visualized on a UMAP, or summarized across cell types or pseudotime windows to study dynamic regulation.

##### 1.9.8 Identifying Downstream Targets of TFs

The final topic-specific GRNs $\mathbf{G}_{\text{final}}^{\text{act/rep}}[t]$ also allow extraction of downstream targets for any transcription factor in a given topic. For a TF $f$, its top regulatory targets in topic $t$ are those genes with the highest values in $\mathbf{G}_{\text{final}}^{\text{act}}[t](f,:)$ or $\mathbf{G}_{\text{final}}^{\text{rep}}[t](f,:)$.

Importantly, the set of target gene and whether a TF acts predominantly as an activator or repressor, can differ across topics. This reflects topic-specific chromatin accessibility patterns: although a TF may bind to the same motif across the genome, the set of accessible peaks it can act through may vary. Thus, in one topic, a TF may bind open enhancers of gene A and while in another, it may bind to enhancers of gene B ( which are closed in the other topic).

These topic-dependent target profiles provide mechanistic insight into how the same TF can play distinct roles across cell states or conditions, driven by changes in chromatin context.

#### 1.10 Related Works: eGRN Inference and Multi-Omic Dimensionality Reduction

Single-cell technologies that co-profile gene expression (RNA) and chromatin accessibility (ATAC) in the *same* cell have provided opportunities to infer accurate eGRNs. From these multi-omic data, three key "signatures" can be leveraged for inferring eGRNs:

1. **TF–Peak Mapping**
   Identifies *which* transcription factors (TFs) potentially bind *which* accessible DNA elements (peaks).
2. **Peak–Gene Linking**
   Discovers *which* regulatory elements (enhancers or promoters) modulate *which* genes, based on accessibility–expression covariation or proximity.
3. **TF–Gene Relationships**
   Connects TF expression/activity to gene expression changes, ideally clarifying which TF is activating or repressing a target gene.

Many published GRN inference pipelines [19, 20, 21, 22, 23, 24, 10, 25, 26] exploit some (or all) of these signatures, but often do so in a stepwise manner—mapping TF motifs, correlating peaks and genes, then refining TF–gene edges. Meanwhile, a separate class of methods focuses on **dimensionality reduction** (DR) or **factor modeling** to jointly embed multi-omic data, typically aiming for a low-dimensional representation that summarizes cellular variation. However, these DR methods rarely impose *mechanistic* constraints (e.g., binding or enhancer logic) on the learned factors.

**scDoRI** fuses these two paradigms—(1) *mechanistic* eGRN inference and (2) *interpretable* dimensionality reduction—within a single deep-learning framework. This fusion enables cells to be represented as combinations of regulatory topics, each corresponding to an interpretable TF–gene program. As a result, scDoRI eliminates the need for precomputed cell clusters, since cells are modeled as mixtures of regulatory programs rather than discrete assignments.

Below, we first review relevant methods in both arenas, then show how scDoRI synthesizes the best of each, and explain why **SCENIC+** is our principal benchmark for multi-omic eGRN inference.

We direct readers to a recent review on eGRN inference using sc multi-ome data for detailed descriptions of different signatures and methods[27].

##### 1.10.1 Methods for eGRN Inference from Single-Cell Multi-Omics

We highlight exemplary methods that capture different strategies for computing regulatory signatures (e.g., TF–peak interactions, peak–gene associations, and TF–gene links). A comprehensive comparison is provided in Table 3.

##### 1.10.2 TF–Peak Mapping

**Motif-based scans.** Classical tools (e.g., HOMER [28], FIMO[6], GimmeMotifs [29]) compare accessible regions against known TF motifs ( cisBP[15], JASPAR [16] or Hocomoco [17]) to identify potential binding sites. This is widely used, but motif degeneracy often inflates false positives, and these scans alone cannot distinguish TF activation from repression.

**SCENIC+ motif enrichment.** **SCENIC+** [10] refines motif-finding with *cisTarget* and differential motif enrichment analyses, reducing spurious hits. However, it still does not attach *activator* or *repressor* labels to TF–peak edges.

**Correlation or footprint-based refinements.**

- *GRANIE*[20] and*InSilico-ChIPseq*[3] build upon observations in DiffTF [4] and weigh motif presence by how strongly a region's accessibility correlates with TF expression (positive correlation suggests TF binding to open chromatin).
- *Footprinting* approaches (e.g., used in **Dictys**[23]) attempt to detect "footprints" left by TF binding in chromatin accessibility profiles.

Most of these approaches remain *unsigned*: they do not explicitly label a TF–peak interaction as "repressive" vs. "activating."

**scDoRI's signed TF–peak edges.** scDoRI extends on the *GRANIE*[20] and*InSilico-ChIPseq*[3] approaches and assigns a sign to TF–peak edges( 1.3):

- Significant Positive correlation (TF expression high *and* peak open) ⇒ **activator edge**.
- Significant Negative correlation (TF expression high *when* peak is less accessible) ⇒ **repressor edge**.

Explicit modeling of repressive vs. activating TF–peak links and integration with further downstream effects on chromatin and gene-expression, enables scDoRI to capture more nuanced regulatory logic.

##### 1.10.3 Peak–Gene Linking

**Correlation- and proximity-based approaches.**

- **CellOracle [25]**: links distal peaks to genes based on co-accessibility (peaks that open/close together with the promoter, regulate that gene).
- **Pando[24]**, **FigR[21]**, **GRANIE[20]**: use correlation or linear modeling between peak accessibility and gene expression.
- **SCENIC+**: uses gradient-boosted trees (GBT) on single-cell multi-ome data to predict gene expression from peak accesibility.
- **Dictys**: considers all peaks within a window of a gene.

Such methods effectively model activation-oriented enhancer logic, but do not explictly model the possibility that certain TFs might *repress* a gene when bound to a region that becomes *closed.*

**scDoRI's peak-gene links** scDORI leverages linear modelling to predict gene-expression from chromatin accesibility (module 2), thereby learning putative peak-gene links. However, it additionally supports an explicit repressive mode (module 4). If a TF with repressive capacity binds a peak, scDoRI can treat *closure* of that region as inhibiting gene expression. This design allows a single locus to switch between activation or repression depending on context (TF identity, expression level, cell state/topic).

##### 1.10.4 TF–Gene Inference

**Expression-based GRN methods.** Earlier pipelines (e.g., **GENIE3**[22]) linked TFs to genes purely by co-expression. **SCENIC [19]** adds motif presence near putative targets as an extra filter. Without actual single-cell ATAC readouts, though, these methods rely primarily on indirect evidence of TF-gene linkage.

**Multi-omic GRN models.** Several recent methods integrate scRNA-seq and scATAC-seq to infer eGRNs (see above), combining information from TF motifs, accessibility, and expression. While these approaches represent a substantial advance over expression-only methods, they share several common limitations:

1. **Cluster dependence**: to obtain context-dependent eGRNs, existing models rely on pre-defined definitons of such context which could include cell-type labels, fine-grained clusters or lineage hierarchies, which can obscure unbiased inference of eGRNs.

2. **Lack of repressor modeling**: negative TF–gene correlations in expression are included, but usually not supported by logic of peak closer by repressor TFs.

3. **Scalability to large datasets at single-cell level**: existing methods either operate at metacell/pseudobulk level or cannot efficiently scale to large datasets of millions of cells.

**scDoRI's cluster-free, integrated eGRN inference approach.**

- **Mini-batch training** allows training with large datatsets, retaining single-cell resolution.

- **Signed TF–gene edges**: scDoRI explicitly propagates repressive or activating status from TF–peak links through to TF–gene edges. A TF bound to an *open* site can activate genes, while the same TF bound to a *closed* site can repress other targets, depending on cellular context.

- **Auto-encoders**: scDORI effectively merges dimensionality reduction with eGRN inference allowing each cell to represented by a combination of Topics, without relying on known labels.

##### 1.10.5 Comparison of eGRN Methods

Table 3: Comparison of selected eGRN pipelines. "eGRN" indicates whether the method outputs connected TF–peak–gene links or just TF–gene links. "TF–Peak Links" refers to how transcription factors are associated with peaks (e.g., motif scanning or correlation). "Peak–Gene Links" denotes the strategy used to link peaks to gene expression. "SC vs. PB?" indicates whether the method operates at single-cell resolution (SC) or uses pseudo-bulk/metacell aggregation (PB). "Signed (A/R)?" reflects whether the method distinguishes activator vs. repressor logic. "Clusters Req.?" specifies whether precomputed cell clusters are required for obtaining cell-type/cell-state specific eGRN. "Batch Correction" indicates how batch effects are handled (internal to the method or requires preprocessing).

| **Method** | **eGRN** | **TF–Peak Links** | **Peak–Gene Links** | **SC vs. PB?** | **Signed (A/R)?** | **Clusters Req.?** | **Batch Correction** |
|---|---|---|---|---|---|---|---|
| **SCENIC+** | Yes | Motif + Diff. Enrichment | GBT (ATAC+RNA) | SC | No | Yes | External |
| **Pando** | Yes | Motif scanning | Corr./Linear (ATAC+RNA) | SC | No | Yes | External |
| **CellOracle** | Yes | Motif scanning | Co-access (ATAC) | SC | No | Yes | External |
| **FigR** | Partial | Motif + Corr. (TF exp & DORC access.) | DORC Corr. (ATAC+RNA) | SC | No | Yes | External |
| **GRANIE** | Yes | Motif + Corr. (TF exp & peak access.) | Corr./Linear (ATAC+RNA) | PB | No | Yes | External |
| **Dictys** | No | Footprinting | Fixed window | SC | No | Yes | Internal |
| **scMTNI** | No | Motif scanning | Promoter peaks | PB | No | Yes | External |
| **scDoRI** | **Yes** | **Motif + Corr. (TF exp & peak access.)** | **Corr./Linear (ATAC+RNA)** | **SC (mini-batch)** | **Yes** | **No** | **Internal** |

As summarized in Table 3, current eGRN inference methods span a range of strategies but often rely on pre-defined clusters, and omit repressor logic. Among these, **SCENIC+** is a strong baseline due to its principled integration of motif enrichment and gene-expression prediction, and it performs favorably in both its own benchmarks and external evaluations [30].

However, SCENIC+ and similar methods are not optimized for large-scale, fully single-cell eGRN inference. scDoRI complements these efforts by learning topic-specific, signed GRNs directly from single-cell RNA + ATAC data without cluster definitions, while scaling to datasets of >1M cells using mini-batch training, enabling interpretable regulatory inference across diverse cellular landscapes.

#### 1.11 Methods for Dimensionality Reduction and Multi-Omic Factorization

While the methods above focus on *regulatory* inference, another body of work targets *dimensionality reduction* (DR) or factor analysis to embed RNA+ATAC data into a low-dimensional space. Such methods typically do not address TF binding or enhancer logic to explain gene-expression directly.

##### 1.11.1 Single-Modality DR/Topic Models

- **cisTopic [8]**: Applies Latent Dirichlet Allocation (LDA) [7] to scATAC-seq to identify "topics" of co-accessible peaks.
- **scETM [31]**,**Expimap [32]**,**VEGA [33]**: Extend autoencoders by incorporating prior biological knowledge (e.g., gene programs or pathways) directly into the decoder, improving interpretability and alignment with known biology.
- **ldVAE [34]**: Uses a linear-decoder variational autoencoder to model scRNA-seq data, preserving the interpretability of latent factors.

While powerful for summarizing variation within *one* modality, these approaches do not unify RNA and ATAC to reveal mechanistic, causal regulatory relationships.

##### 1.11.2 Multi-Omic Factor Models

Several methods aim to jointly embed scRNA-seq and scATAC-seq data into a shared low-dimensional space, often for visualization, clustering, or integration. While useful for summarizing variation across modalities, these models typically do not explicitly reconstruct gene expression through TF-mediated enhancer activity.

- **MOFA [35]**: factor analysis across multiple omics, producing a shared latent space but without directing how ATAC influences RNA.
- **shareTopic [36]**, **moETM [37]**: attempt a "joint" topic model on RNA+ATAC but typically decode each modality independently, lacking explicit TF binding constraints.
- **MIRA [38]**: integrates RNA+ATAC embeddings, then performs *post-hoc* regulatory potential analysis to link peaks with genes.
- **GLUE [39]**: Constructs a shared embedding of scRNA-seq and scATAC-seq by incorporating regulatory associations peaks and genes, based on genomic proximity into a graph-regularized encoder-decoder framework. While scalable and effective for multimodal integration, GLUE does not explicitly model TF-driven chromatin dynamics as causal intermediates for explaining gene expression.

##### 1.11.3 scDoRI's Mechanistic Autoencoder

By embedding explicit TF binding logic, chromatin accesibility dynamics and peak–gene relationships in the decoder, scDoRI's latent topics gain a *biologically grounded* meaning. Instead of generic factors, each topic represents a GRN, connecting sets of peaks and genes under specific TF regulators.

#### 1.12 Benchmarking Strategy

##### 1.12.1 Benchmarking on Semi-synthetic ENCODE Data (Extended Data Figure S2C)

We first validated `scDORI` on semi-synthetic data and compared it against alternative GRN inference methods, adapting a previously proposed benchmarking strategy for GRN inference in Scenic+ [10]. Specifically, we leveraged simulated single-cell multi-omic profiles (RNA and ATAC) of 4,000 cells derived from 8 distinct ENCODE cell lines. These profiles recapitulate real-world single-cell scenarios, while having additional molecular data (e.g., CRISPR knockouts, Hi-C profiles, and eQTLs) to assess inferred regulatory linkages.

We considered 3,068 highly variable genes (HVGs) and 119,662 highly variable peaks. The peaks were chosen from a 150 kbp window around each gene body of the HVGs. Both scDoRI and SCENIC+ were trained on this dataset.

**Enhancer–gene link predictions.** To evaluate enhancer–gene link predictions, we used the benchmark introduced by [40]. This resource provides a set of positive enhancer–gene links (validated via CRISPR knockouts, chromatin conformation assays, or eQTL relationships) alongside a set of negative links. We restricted our analysis to benchmarked cell type–evidence pairs with at least 10 positive enhancer–gene pairs overlapping our data. Both `scDORI` and `SCENIC+` were used to assign peak–gene linkages, and performance was quantified by the Area Under the Precision–Recall Curve (AUPRC).

##### 1.12.2 Benchmarking Latent Spaces for explaining cell-state differences in GB atlas (Extended Data Figure S4C)

We benchmarked the latent spaces produced by different methods on the GB atlas, focusing on their ability to resolve cell-state differences. Specifically, we compared:

1. The latent space derived from `scDORI`.
2. A latent space constructed from 50 MOFA factors, obtained by applying MOFA to aggregated multi-omic metacell profiles (see Section 2.2.1 for metacell construction details). MOFA [35] is a popular unsupervised factor model for multi-omics integration that learns a joint low-dimensional representation of each cell. It serves as a natural comparison point for scDoRI, as both methods aim to summarize expression and accessibility variation through latent factors. However, unlike scDoRI, MOFA does not explicitly couple RNA and ATAC modalities through regulatory priors. .
3. A `SCENIC+`-based latent space, obtained by computing AUCell [41] scores for 195 transcription factors (TFs) which were also used to train `scDORI`, based on GRNs inferred from metacell-level data.

**SCENIC+ on GBM Data.** Given the scalability constraints of `SCENIC+` on the full GBM atlas, we aggregated single cells into metacells (see Section 2.2.1) and then applied the `SCENIC+` pipeline to these metacell profiles. For transcription factors that did not have direct targets in `SCENIC+`'s primary output, we incorporated indirect targets derived from expression-based criteria (an auxiliary output of `SCENIC+`). This step ensured that `SCENIC+` and `scDORI` relied on a unified TF set, thereby enabling downstream analyses—such as the TAP analysis in Section 2.5—to be conducted in a fair, comparable manner.

**MOFA on Metacell Data.** Similarly, due to the memory and computational demands of MOFA on large-scale datasets, we applied MOFA to the aggregated metacell-level multi-omic profiles rather than raw single-cell data. From this analysis, we extracted 50 latent factors intended to capture the principal axes of variation within the GBM atlas.

**scDORI Topics at the Metacell Level.** For `scDORI`, we computed the average topic activations for each metacell by averaging the topic scores of all single cells constituting that metacell, yielding an embedding of 50 topics.

**Evaluation strategy.** We assessed each of the three latent representations by predicting cell-state labels originally derived from gene expression (see Figure 1). In particular, we trained a logistic regression model under a 5 fold cross-validation scheme and measured classification accuracy.

##### 1.12.3 Benchmarking on Neftel Dataset [1]

To further evaluate `scDORI` GRNs, we adopted the approach of GranPA[20] to assess how well inferred GRNs explain gene expression differences across cell-state in an orthogonal GB single-cell dataset by Neftel *et al.* [1] (Extended Data Figure S5D,E).

**Evaluation strategy.** We used GB cell-state assignments (AC-like, MES-like1, MES-like2, NPC-like1, NPC-like2, and OPC-like) and fine-grained Leiden clusters from the Neftel dataset (10x genomics kit processed cells). For each pair of cell states (or Leiden clusters), we performed differential gene expression analysis and considered genes with adjusted $p$-value $< 0.05$. Genes with log-fold change greater than 0 were labeled as "positive" (i.e., significantly upregulated ), while the remaining genes were labeled as "negative." (i.e., significantly downregulated ). Only pairs with at least 10 upregulated and 10 downregulated genes were considered; for Leiden clusters, we further restricted to clusters with more than 200 cells, resulting in 104 pairwise comparisons among Leiden clusters and 15 pairwise comparisons among the major GB states.

We then adapted the approach in [20] to train a logistic regression model (implemented in `scikit-learn` with `class_weight=balanced` and C = 0.01 for L2 regularization) that uses inferred TF–gene interactions to predict whether a gene is differentially upregulated (1) or downregulated (0). Importantly, each TF was labeled with +1 or −1, indicating an activating or repressing interaction. The model was evaluated in a 4-fold cross-validation scheme, measuring performance by the AUPRC. Additionally, to handle potential asymmetry in "up" vs. "down" regulation, we trained a separate model per pair of clusters with reversed up-/down-regulated labels and averaged the results.

**Mapping `scDORI` topics to Neftel data.** We started with a matrix of topic–gene weights derived from Module 2 of scDoRI (1.9.3), where rows correspond to topics and columns correspond to the genes used during `scDORI` inference. Let $\mathbf{W} \in \mathbb{R}^{K \times G}$ denote this matrix, where is the number of topics and $G$ is the number of genes. We also obtained the Neftel gene-expression matrix $\mathbf{X} \in \mathbb{R}^{N \times G}$, where $N$ is the number of single cells and $G$ is the subset of genes overlapping with $\mathbf{W}$.

**Normalization and topic activation.** To account for differences in sequencing depth, we normalized each cell's total expression to a common scale given by the median total counts across cells:

$$X'_{ig} \;=\; m \;\cdot\; \frac{X_{ig}}{\sum_{g'} X_{ig'}}, \quad \text{where } m = \text{median}_i\Big(\sum_{g'} X_{ig'}\Big).$$

We then computed the *topic activation matrix*, $\mathbf{T} \in \mathbb{R}^{N \times K}$, via matrix multiplication:

$$T_{ik} \;=\; \sum_{g=1}^{G} X'_{ig}\, W_{kg}.$$

Each entry $T_{ik}$ represents the activation score of topic $k$ in cell $i$. Finally, we applied a min–max scaling per topic to obtain values in $[0, 1]$.

**Constructing cell-state-specific GRNs from scDoRI.** Using the annotated cell-state or cluster labels (e.g., AC-like, MES-like1, MES-like2, NPC-like1, NPC-like2, OPC-like, or Leiden clusters), we aggregated the topic activation scores. For any pair of clusters, we identified the union of topics whose average activation score exceeded a threshold of 0.2 in at least one cluster. The cell-state-specific GRN was then defined by taking the union of TF–gene links from those selected topics. Hence, if a TF–gene edge appeared in any of the high-activation topics, it was included for that cluster pair's GRN.

**Comparison with `SCENIC+`.** We used GRNs derived from application of SCENIC+ to metacell-level GB atlas as described in the previous section, using a global(cell-state agnostic) network of TF-gene as input features of the logistic regression model.

### 2 Analysis of GB multi-ome atlas

#### 2.1 Multi-ome Data Quality Control, Integration and annotation

For detailed methodology, refer to the companion manuscript by De jong et al. Briefly, reads from snRNA-seq and ATAC-seq libraries were aligned to a custom genome combining the 10X Genomics GRCh38 pre-mRNA reference and Cell Ranger-Arc ATAC genome (v1.0.1). Initial quantification and filtering were performed using default parameters of the Cell Ranger-Arc pipeline. RNA data quality control involved filtering nuclei based on gene and UMI counts, as well as mitochondrial read proportions, to exclude low-quality or stressed cells. Doublet detection and removal were conducted using Scrublet [42], followed by additional filtering at both the individual cell and cluster levels to refine the dataset.

ATAC data were processed with ArchR [9], applying thresholds on fragment counts and transcriptional start site enrichments to retain high-quality barcodes. RNA quality-filtered barcodes were used to filter the ATAC data, ensuring consistency between modalities.

This filtering step resulted in 1,025,329 nuclei, which we used for training cell-type label agnostic scDoRI model and obtaining results shown in the paper.

snRNA-seq datasets were integrated using `scVI` [2], incorporating patient ID, tumor site, reaction date, and cell cycle phase as batch covariates. Highly variable genes were identified using dispersion-based methods, and clustering was performed in the `scVI` latent space. Malignant clusters were further identified using copy number variation profiles (e.g., characteristic gains and losses) inferred with `inferCNV`[43]. Clusters were annotated into malignant or tumor microenvironment (TME) populations based on marker gene expression.

Subclustering of malignant and TME populations enabled more granular annotation. TME cell types were annotated using known human brain marker genes, while malignant subclusters were classified using a combination of gene module scoring (following Neftel et al. [1]), projection onto a developmental brain atlas [44], and gene set enrichment.

Refer to the companion manuscript by De jong et al. for more details.

#### 2.2 Epigenetic Plasticity Score

**Rationale.** Simultaneous measurements of gene expression (RNA) and chromatin accessibility (ATAC) in single cells enable the assessment of the coupling between these two regulatory layers. While chromatin accessibility and transcription are often coordinated, regions of accessible chromatin that are not yet accompanied by active gene expression may indicate a primed state, poised for future transcriptional activation. This decoupling reveals a cell's latent potential to switch or activate alternative transcriptional programs—a feature we term *epigenetic plasticity*: the capacity of a cell's chromatin landscape to support multiple or alternative transcriptional identities. To quantify this phenomenon, we adapted a framework from [45], which leverages a linear classifier trained on RNA profiles but subsequently applied to ATAC-derived gene scores. These gene scores were computed using ArchR [9] by aggregating accessibility over promoter regions, gene bodies, and distal regulatory elements weighted by distance. Higher uncertainty or misclassification in the ATAC-based predictions reflects greater chromatin priming and, consequently, elevated epigenetic plasticity (Fig 1B,C).

##### 2.2.1 Plasticity Score Estimation Using Cell Type Predictor (Figure 1B, C)

Following the approach proposed by [45], we first constructed meta-cells by performing Leiden clustering using Scanpy [46] (via the `tl.leiden` function with a resolution parameter set to 1) in the RNA-space embedding generated by scVI [2]. Clustering was performed independently for cells belonging to each unique combination of cell state, donor, and batch. Cell-state-donor-batch groups with fewer than 40 cells were excluded from clustering and treated as individual meta-cells. This procedure yielded a total of 16,243 meta-cells.

For each meta-cell, we calculated the mean expression of all genes using the RNA counts matrix and the ATAC-derived gene scores computed with ArchR. The resulting mean RNA and ATAC matrices were normalized for library size, with RNA counts further log-transformed.

To quantify epigenetic plasticity, we trained a logistic regression model with a multinomial likelihood to predict cell states (9 GBM and 8 TME states, as defined in Fig 1A) based on gene expression. Specifically, we used scaled expression values (via scikit-learn's [47]`StandardScaler`) of 3,160 highly variable genes as input features. The RNA model was trained and tested using scikit-learn, with 5-fold cross-validation to ensure robustness. The trained model was then applied to predict cell states for each meta-cell using scaled ATAC gene score values as input. We computed the Shannon entropy over the predicted probabilities across cell types for each meta-cell; higher entropy indicated greater discordance between RNA and ATAC profiles, reflecting increased epigenetic plasticity.

To visualize epigenetic plasticity at the single-cell level (Fig 1C), we assigned the meta-cell-level plasticity scores back to the individual cells that composed each meta-cell.

##### 2.2.2 Variance Explained in Epigenetic Plasticity Score (Figure 1F)

To investigate sources of variation in plasticity, we modeled the entropy-based plasticity score—computed per meta-cell—as a function of categorical variables representing cell state and donor identity. Specifically, we fit an ordinary least squares (OLS) regression model using the `statsmodels` Python library [48]:

$$\text{Entropy}_i = \beta_0 + \beta_{\text{state}[i]} + \beta_{\text{donor}[i]} + \varepsilon_i,$$

where $\text{Entropy}_i$ denotes the plasticity score for metacell $i$, and $\beta_{\text{state}[i]}$, $\beta_{\text{donor}[i]}$ represent the fixed effects for cell state and donor, respectively. We then performed an ANOVA to quantify the variance explained by each factor. Similar results were observed when using linear mixed models with donor as a random effect and cell state as a fixed effect (results not shown).

##### 2.2.3 Neighborhood-Based Analysis of Donor Contributions to Plasticity Variation (Extended Data Figure S1F)

To further assess whether the observed variation in plasticity scores is predominantly driven by cell-state differences rather than donor-specific effects, we performed a neighborhood analysis around each meta-cell. Specifically, for each meta-cell in the dataset, we identified a local neighborhood comprising either 10 or 30 nearest neighbors (in scVI RNA embedding ).

Within each neighborhood, we calculated two metrics:

1. The standard deviation of plasticity scores ($SD_{plasticity}$).
2. The standard deviation of plasticity scores *across* the unique donors present in that neighborhood ($SD_{donor}$).

We then took the ratio of $SD_{donor}$ to $SD_{plasticity}$ as a measure of how much donor identity contributes to the local variation in plasticity. Neighborhoods containing fewer than three unique donors were discarded to avoid unreliable estimates.

###### 2.2.4 Assessing Global RNA–ATAC Coupling as a Control for Epigenetic Plasticity (Extended Data Figure S1C)

As an additional control for our epigenetic plasticity claims, we examined whether certain cell states might exhibit more globally "permissive" chromatin—i.e., lower RNA–ATAC coupling that could spuriously inflate plasticity estimates. To do so, we computed the Pearson correlation coefficient ($r$) between the mean RNA expression and the corresponding ATAC-derived gene scores across 21,369 genes for each meta-cell. If a particular cell state had systematically weaker RNA–ATAC linkage (i.e., lower correlation across most genes), one might expect a broader decoupling between chromatin accessibility and transcription, potentially leading to higher plasticity scores. Conversely, states with stronger RNA–ATAC correlation might appear less "plastic" according to our entropy-based metric. By confirming that states exhibiting higher epigenetic plasticity do not simply have globally poor RNA–ATAC coupling, we strengthen the interpretation that our plasticity scores indeed reflect a cell's latent ability to adopt alternative transcriptional programs, rather than artifactually low gene score–expression concordance.

#### 2.3 scDoRI Hyperparameters for GB Data

In this section, we detail the primary hyperparameters and dataset attributes used when training **scDoRI** on a GB atlas.

Table 4: Dataset Details and Hyperparameters for scDoRI on GBM

| Parameter | Description | Value |
|---|---|---|
| | *Dataset Attributes* | |
| num_cells | Number of single cells | 1,025,329 |
| num_genes (G) | Number of genes used | 3192 |
| num_peaks (P) | Number of peaks used | 182677 |
| num_TFs (F) | Number of TFs recognized | 195 |
| | *scDoRI Hyperparameters* | |
| num_topics | Number of latent topics (T) | 50 |
| learning_rate | Initial Adam learning rate | $1 \times 10^{-2}$ |
| weight_decay | L2 weight decay for Adam | 0 |
| batch_size | Cells per mini-batch | 16 |
| $\beta_{\text{ATAC}}$ | Loss weight for ATAC reconstruction | 1 |
| $\beta_{\text{RNA-ATAC}}$ | Loss weight for RNA-from-ATAC | 100 |
| $\beta_{\text{TF}}$ | Loss weight for TF reconstruction | 200 |
| $\lambda_1$ | L1 reg. on topic–peak decoder | $1 \times 10^{-5}$ |
| $\lambda_2$ | L2 reg. on gene–peak weights | $1 \times 10^{-5}$ |
| early_stopping_patience | Epochs w/o improvement before stop | 2 |

During Phase 2 training for GRN inference (Module 4), we **froze the encoder and Modules 1–3** to preserve the topic representations learned in Phase 1. Additionally, we used **empirical topic-level TF expression**—computed by averaging TF expression across cells in each topic as input to Module 4 , rather than using the denoised TF estimates from Module 3.

For notes on selecting hyperparameters, refer to Section 1.8.

#### 2.4 Epigenetic priming of TRs

To assess epigenetic priming of TRs in GB cells, we examined discrepancies between their expression and chromatin accessibility across the dataset. Due to the high heterogeneity among GB cells, we used topic activation patterns to define finer-grained cell subsets that share similar regulatory dynamics. For each topic, we selected the top 5000 cells with the highest activation scores and computed the average TR expression and accessibility within these cells.

TR accessibility was quantified using the Gene Score function from ArchR [9], which estimates chromatin accessibility for each gene by aggregating ATAC-seq signal across gene body, promoter, and distance-weighted distal regulatory elements. To enable comparison across topics, both TR expression and accessibility values were min-max scaled across all topics.

We defined a TR as **epigenetically primed** in a given topic-defined substate if its scaled expression was below 0.3 while its scaled accessibility was above 0.5 (Extended Data Figure S6A,B).

#### 2.5 Topic Activation Potential (TAP)

##### 2.5.1 Rationale and Theoretical Basis

To investigate potential regulatory interactions between scDoRI topics and their effect on state transitions, we developed a computational metric to quantify the Topic Activation Potential (TAP) between pairs of topics. This approach leverages scDoRI's inferred eGRNs and is motivated by the observation that key topic regulators (TRs) can be epigenetically accessible, even if not currently expressed, in alternate states, suggesting potential for future activation. The TAP provides a quantitative measure of the ease with which cells with a given active topic $A$ can activate a distinct topic $B$.

The core premise of our framework is that activating the TRs of the target topic $B$ is sufficient to drive its respective activation. This concept is analogous to cellular reprogramming strategies, where overexpression of key TFs induces transdifferentiation or pluripotency [49].

Two primary scenarios underpin this framework:

1. **Shared TR Expression:** If TRs characteristic of topic $B$ are already expressed in cells with topic $A$, the transition is facilitated due to the existing expression of critical regulatory factors. This scenario is common between transcriptionally similar topic-defined substates (i.e., subset of cells where each topic is active).

2. **Epigenetic Priming:** For transcriptionally distinct states where TRs of topic $B$ are not expressed in cells with topic $A$, transitions may occur if these TRs are epigenetically primed — that is, their chromatin regions are accessible in cells with topic $A$ (Extended Data Figure S6A,B). This accessibility lowers the activation barrier, making it easier for these TRs to be expressed upon receiving appropriate regulatory signals.

To compute the TAP between a given topic pair, we consider three major components: (i) the regulatory influence of TFs in topic $A$ over TRs of topic $B$, (ii) accesibility (priming) of TRs of topic $B$ within cells with topic $A$, and (iii) the extent of downstream effects of TRs in topic $B$ (Figure 3C). These components are detailed in the following subheadings.

###### 2.5.2 TAP components

**Regulatory Influence from source TFs to target TRs** We consider all potential regulatory interactions that promote the activation of TRs of the target topic $B$ based on predicted links in ATAC-derived scDoRI GRNs (defined in 3). Using ATAC-derived GRNs $\mathbf{G}^{\text{act}}_{\text{ATAC}}[t]$, rather than final expression-pruned GRNs, allows us to consider all regulatory links even if target TRs are accessible but not yet expressed in cells with topic $A$, providing a more permissive view of the regulatory landscape influencing state transitions.

We take all links from a TF ($Y$) expressed in cells with topic $A$ (min-max scaled average expression >0.5) towards the TRs ($X$) in topic $B$ (TFs with topic activity score from 1.9.7 >0.05) based on the ATAC-derived GRNs of topic $A$. Specifically, the regulatory influence $\text{Reg}(Y, X)$ is defined as an indicator variable:

$$\text{Reg}(Y, X) = \begin{cases} 1, & \text{if there is a regulatory link from } Y \text{ to } X \text{ in} \\ G^{\text{act}}_{\text{ATAC}}[A] & \\ 0, & \text{otherwise} \end{cases}$$

where:

- Intuitively, a regulatory link exists if TF $Y$ is predicted to bind to accessible regulatory regions (e.g., enhancers, promoters) of TR $X$ in topic $A$.

**Accesibility (Priming) of TRs in Source Topic** Priming reflects the chromatin accessibility of TRs in cells with source topic $A$, indicating their readiness for activation. For each TR $X$, the priming score $\text{Priming}(X, A)$ is defined as the average chromatin accessibility of $X$ within the subset of cells with highest topic $A$ activity (top 5000 cells with highest activation):

$$\text{Priming}(X, A) = \text{Accessibility}(X, A)$$

where:

- $\text{Accessibility}(X, A)$ is the averaged Gene Score of TR $X$ (derived from ArchR[9]) across cells with high topic $A$ activity, subsequently minmax scaled across other topics.

**Downstream effect of TRs in target topic** For each TR $X$ in the target topic $B$, we define its total downstream activation effect over topic $B$ as its activity score in topic $B$ - ACT(X, B) (as described in 1.9.7).

###### 2.5.3 Computation

**Computation of Topic Activation Potential** By considering all three components above, the TAP from topic $A$ to topic $B$ quantifies the overall likelihood of cells harboring topic $A$ to activate the gene regulatory programs defined by topic $B$, and it is defined as follows:

$$\text{TAP}_{A \to B} = \frac{1}{|\mathcal{F}_A|} \sum_{Y \in \mathcal{F}_A} \sum_{X \in \mathcal{R}_B} \text{Reg}(Y, X) \times \text{Priming}(X, A) \times \text{ACT}(X, B)$$

where:

- $\mathcal{F}_A$ is the set of TFs expressed in cells with topic $A$ (>0.5 scaled average expression)

- $\mathcal{R}_B$ is the set of TRs of topic $B$
- $|\mathcal{F}_A|$ is the total number of TFs expressed in cells with topic $A$

Accordingly, this metric captures the cumulative effect of all regulatory interactions with the potential of activating TRs of topic $B$, each weighted by how accessible those TRs are for source cells and their relative importance for activating topic $B$. We compute TAP values for all topic pairs active in GB cells, except pan-state topics which are broadly active across all GB cells (Topics 18,31,39).

To determine the empirical significance of each TAP link, we randomized TF-gene links in ATAC-derived GRNs of each topic 1000 times and computed $\text{TAP}_{\text{A} \rightarrow \text{B}}$ values resulting from each. A $\text{TAP}_{\text{A} \rightarrow \text{B}}$ link was considered significant if its strength exceeded the 90th percentile of the distribution of randomized $\text{TAP}_{\text{A} \rightarrow \text{B}}$ links.

**Normalization Across Source Topics** To compare TAP values across different source topics for a given target topic $B$, we normalized the TAP values by the self-activation value of topic $B$ ($\text{TAP}_{\text{B} \rightarrow \text{B}}$:

$$\text{Normalized TAP}_{A \to B} = \frac{\text{TAP}_{A \to B}}{\text{TAP}_{B \to B}}$$

This normalization scales all TAP values relative to the total activating effect of TFs on their own topic, many of which are already TRs, thus acting as a positive control. An exception to this was Topic 37, whose targeted population of cells is a subset of that of Topic 24 (see Extended Data Fig. 3B) and whose self-activation was considerably low. In this case, TAP values from Topic 24 to Topic 37 served as positive control. Pair-wise TAP scores and driver TFs behind each TAP interaction can be found in **Supplementary Table 3**.

##### 2.5.4 Variance Decomposition of TAP Values

To assess the factors contributing to variation in TAP scores across topic pairs, we performed variance decomposition using linear regression. Each $\text{TAP}_{\text{A} \to \text{B}}$ value was modeled as a function of interpretable features capturing regulatory wiring and chromatin readiness of the target topic's TRs.

The following predictors were used:

- **Number of strong TRs targeted / total TRs:** Proportion of TRs in topic $B$ with high activity scores ($\text{ACT}_{\text{X, B}} > 0.5$) that are also targeted by TFs expressed in topic $A$.
- **Proportion of TRs targeted:** Fraction of TRs in topic $B$ with at least one regulatory link from TFs expressed in topic $A$.
- **TF–TR connectivity:** Total number of TF–TR regulatory links from topic $A$ to $B$, normalized by the number of TRs.
- **Average accessibility of TRs in source topic:** Mean chromatin accessibility of topic $B$ TRs in cells enriched for topic $A$.
- **Average expression of TRs in source topic:** Mean expression of topic $B$ TRs in topic $A$ cells.

To evaluate context-specific contributions, the analysis was performed separately for TAP links targeting (i) topics from the same or closely related states, (ii) topics from alternate states, and (iii) all topics. Variance explained by each feature was estimated using $\text{R}^2$ scores, and results are shown in Extended Data Fig. 6E.

##### 2.5.5 TAP calculation with alternative eGRN inference methods

We derived GRNs from SCENIC+ (as described in 1.12.2). To define TFs that would serve equivalently as scDoRI TRs, we first use AUcell [41] to score downstream regulatory influence of each TF within each topic (considering average AUcell activity of each TF in top 5000 cells with highest activation for each topic). Accordingly, we compute TAP values with the following modifications:

- Reg(Y,X) denotes the presence of a predicted activation link from a TF $Y$ expressed in topic $A$ towards a TR $X$ of topic $B$ (as determined with AUCell). Here, $TF_Y$ - $TF_X$ links are derived from the GRN inferred with SCENIC+.
- Priming (X,A) denotes the scaled average accessibility of TF $X$ within the subset of cells defined by topic $A$ activity.
- ACT(X,B) denotes the AUCell activity score for TF $X$ in topic $B$.

- $\mathcal{F}_A$ denotes the sets of TFs expressed in the subset of cells defined by topic $A$ activity and $\mathcal{R}_B$ the set of TFs enriched in the cells defined by topic $B$ activity (AUCell top 10).

To demonstrate the beenfit of using topic-specific GRNs, we also computed TAP values using an **average scDoRI GRN**, which considered a union of all TF-gene links captured within all topics active in GB cells (TF-gene link in atleast one GB topic).

###### 2.5.6 Comparison with Related Methods and approaches

The computational prediction of cell state transitions builds upon decades of biological insights into cellular reprogramming and epigenetic regulation. Below, we contextualize our Topic Activation Potential (TAP) framework within these foundations and highlight its novel integration of multi-omic data and regulatory network topology to quantify transition likelihoods.

**Transdifferentiation and Transcription Factor–Driven Reprogramming** The paradigm of TF-mediated reprogramming was established by Takahashi and Yamanaka, who demonstrated that ectopic expression of four transcription factors (OCT4, SOX2, KLF4, c-MYC) could reprogram somatic cells into induced pluripotent stem cells (iPSCs) [49]. Subsequent work extended this principle to direct lineage conversions, such as fibroblasts to neurons [50], bypassing pluripotency. Computational tools like Mogrify [51] systematized TF selection by leveraging RNA-seq data and GRNs to identify minimal TF cocktails for transdifferentiation. However, Mogrify's reliance on transcriptomic data limits its ability to account for epigenetic barriers to TF activation. Furthermore, the GRNs used in Mogrify are static across cellular-contexts and ingore repressive interactions.

**Epigenetic Priming and Lineage Competence** Lineage priming—the phenomenon wherein multipotent progenitors exhibit accessible chromatin at lineage-specific loci prior to differentiation—has been extensively documented in hematopoiesis [52]. Single-cell multi-omics approaches, such as SHARE-seq [53], built on this concept by identifying Domains of Regulatory Chromatin (DORCs), where chromatin accessibility precedes and predicts transcriptional activation. While existing tools like STEMNET [52] infer lineage biases from RNA data, they lack the resolution to assess chromatin-based priming. TAP operationalizes these insights by explicitly quantifying the accessibility of target TF's regulatory elements in the source state.

**Trajectory Inference and Transition Probabilities** Trajectory inference methods, including CellRank [54], Waddington-OT [55], and Palantir [56], model state transitions using pseudotemporal ordering, RNA velocity, or optimal transport. These approaches excel at reconstructing continuous differentiation trajectories but struggle with discontinuous transitions (e.g., transdifferentiation) and are limited to transcriptomic data. Slingshot [57] and Monocle [58] address similar challenges but remain rooted in transcriptomic similarity. In contrast, TAP quantifies transition likelihoods through GRN connectivity and epigenetic priming, providing a regulatory logic for why transitions occur, rather than merely when they might happen.

In summary, TAP provides a multi-dimensional, regulatory view of potential cell state transitions by integrating topic-specific eGRNs with chromatin accessibility and TF activity. It combines four key aspects—multi-omic regulatory networks, epigenetic priming, TF-to-TF influence, and empirical filtering, to assess how likely one regulatory program is to activate another. Unlike trajectory inference methods that focus on *when* transitions occur, TAP offers insight into *why* transitions may be mechanistically plausible. While exploratory in nature, it complements existing approaches by grounding transition potential in regulatory logic and providing a structured lens through which to interpret dynamic cellular plasticity.

#### 2.6 Topic Repression Score (TRS)

##### 2.6.1 Definition

The **Topic Repression Score (TRS)** mirrors the Topic Activation Potential (TAP) but focuses on quantifying **repressive** interactions between topics (Figure 4A). Specifically, we:

- Use repressive instead of activating regulatory links $\mathrm{RegRep}(Y, X)$ between each TF $Y$ expressed in cells with topic $A$ and each TR $X$ of topic $B$. Here, we consider all links within $\mathbf{G}^{\mathrm{rep}}_{\mathrm{combined}}[A]$ corresponding to topic $A$.

- Replace the extent of TR accessibility in cells with topic $A$ with $Closure(X, A) = 1 - \mathrm{Accessibility}(X, A)$ to quantify the extent of chromatin closure around TRs, potentially favoring repressive effects.

- Use the *same* TR activity measure $\mathrm{ACT}(X, B)$ as in TAP to assess the importance of targeting each TR $X$ in topic $B$.

Accordingly, for each source topic $A$ and target topic $B$, we define the TRS as follows:

$$\mathrm{TRS}_{A\to B}^{(\mathrm{TR})} = \frac{1}{|\mathcal{F}_A|} \sum_{Y\in\mathcal{F}_A} \sum_{X\in\mathcal{R}_B} \mathrm{RegRep}(Y, X) \times \mathrm{Closure}(X, A) \times \mathrm{ACT}(X, B)$$

where:

- $\mathcal{F}_A$ is the set of TFs expressed in cells with topic $A$ (>0.5 scaled average expression)
- $\mathcal{R}_B$ is the set of TRs of topic $B$
- $|\mathcal{F}_A|$ is the total number of TFs expressed in cells with topic $A$

Similar to TAP links, we assess significance of TRS links by calculating pairwise TRS values with randomized GRNs 1000 times. We define all TRS links that exceed the 90th percentile of this distribution as significant.

Since TAP and TRS values for a given topic pair are derived by assessing activating and repressive interactions between the same set of source TFs and target TRs, we normalize all TRS values targeting a topic $B$ by the same values used to normalize TAP values (the level of self-activation in topic $B$), as follows:

$$\text{Normalized } \mathrm{TRS}_{A\to B} = \frac{\mathrm{TRS}_{A\to B}}{\mathrm{TAP}_{B\to B}}$$

This allows us to set a relative scale to compare all activating (TAP) and repressive (TRS) interactions targeting each topic. Pair-wise TRS scores and driver TFs behind each TRS interaction can be found in **Supplementary Table 3**.

##### 2.6.2 TRS at the marker gene level

To assess the broader repressive influence on key topic- or state-specific gene programs, we extended the TRS metric to focus on repression targeting the top 50 genes of each topic or the top 50 marker genes of each cell state (Extended Data Figure S7B, C).

Top genes per topic were identified by ranking genes according to their weights in the Topic–Gene matrix from Module 2 (see Section 1.4.4). For cell states, top marker genes were ranked based on their weights in the trained linear regression model used to quantify plasticity (Section 2.2.1).

In both cases, we applied min–max scaling to these rankings to derive a weighting factor for each gene within a given topic or state $B$, denoted as $\mathrm{ACT}(X, B)$, representing the gene's relative importance. Apart from incorporating these weights, the TRS computation followed the same procedure described earlier.

##### 2.6.3 Total repression

To quantify how strongly each topic represses alternative gene programs, we aggregated TRS values across topics. Specifically, we defined the total repression score for each topic $A$ as:

$$\text{Total Repression Score}(A) = \sum_{I\in T\setminus\{A\}} \mathrm{TRS}_{A\to I}$$

where $T \setminus \{A\}$ denotes the set of all topics excluding topic $A$.

To evaluate the biological relevance of these scores, we assessed their concordance with epigenetic plasticity across topics. Topic-level plasticity was quantified by using the epigenetic plasticity scores (Section 2.2.1) derived from the top 5000 cells associated with each topic. Pearson correlation was used to assess the relationship between total repression and plasticity. For comparison, we performed the same analysis using total repression scores computed from SCENIC+-derived and averaged GRNs inferred by scDoRI (Extended Data Figure S8C).

##### 2.6.4 Repression of repressor TFs

We further adapted the TRS framework to quantify targeted repression of top repressor TFs within each topic, instead of activator TRs(Extended Data Figure S9E, F). Specifically, we considered all repressive interactions originating from TFs expressed in a given source topic and targeting TFs that exhibit the strongest cumulative repressive potential in each topic (as defined in 1.9.7). The rest of the computation followed the same procedure as described in Section 2.6.1.

#### 2.7 Net topic transition potential (TAP-TRS)

To gain a more complete view of GB state transitions, we integrated both activation and repression to assess how these opposing forces collectively shape transitions. Specifically, we defined a net topic transition potential as the difference between TAP and TRS values for every pair of topics (Extended Data Figure S7D), reflecting the net effect of all activating and repressing interactions between topics:

$$\text{Net topic transition potential}(A \to B) = \text{TAP}_{A\to B} - \text{TRS}_{A\to B}$$

##### 2.7.1 Topic-driven plasticity

To estimate how each topic contributes to plasticity, we summed its net regulatory influence (TAP–TRS) toward topics that are active in alternate cell states. The *topic-driven plasticity* score for a given topic $A$ was computed as:

$$\text{Topic-Driven Plasticity}(A) = \sum_{I\in G_{\text{alt}}} \text{TAP–TRS}_{A\to I}$$

where $G_{\text{alt}}$ denotes the set of target topics that are active in a different cell state from topic $A$. Note that target topics may also be active in the source topic's state, but are not exclusive to it.

We assessed the concordance between topic-driven plasticity scores and epigenetic plasticity values, which were derived from the epigenetic plasticity scores (Section 2.2.1) of the top 5000 cells assigned to each topic. Pearson correlation was used to quantify the relationship. For benchmarking, we performed the same analysis using topic-driven plasticity scores computed from TAP values inferred using SCENIC+ and average GRNs from scDORI (Extended Data Figure S8F).

#### 2.8 State transition potential

To understand how regulatory interactions between topics drive transitions between cell states, we computed the net influence of all activating and repressing links between pairs of states. The goal is to quantify whether one state promotes or inhibits transitions into another based on the cumulative behavior of its active gene programs.

Intuitively, we measure how strongly topics active in one state (State $A$) activate or repress topics active in another state (State $B$). We do this by summing all TAP (activating) or TRS (repressing) interactions from topics in $A$ to those in $B$, weighted by the relative activity of those topics within their respective states (Figure 3D, E).

Formally, we define:

$$\text{StateActivationPotential}_{A\to B} = \sum_{(i,j)\in S_A\times S_B} \alpha_i\beta_j \, \text{TAP}_{i\to j}$$

$$\text{StateRepressionPotential}_{A\to B} = \sum_{(i,j)\in S_A\times S_B} \alpha_i\beta_j \, \text{TRS}_{i\to j}$$

Here, $S_A$ and $S_B$ are the sets of active topics in State $A$ and State $B$, respectively. The weights $\alpha_i$ and $\beta_j$ represent the relative activation of topic $i$ within State $A$, and topic $j$ within State $B$, normalized across all topics in those states.

The *State Transition Potential* is then defined as the difference between the activation and repression potentials:

$$\text{StateTransitionPotential}_{A\to B} = \text{StateActivationPotential}_{A\to B} - \text{StateRepressionPotential}_{A\to B}$$

A positive value indicates that activation dominates repression, suggesting a favorable regulatory environment for transitioning from State $A$ to State $B$. A negative value does not imply that a transition is impossible, but rather that repression is dominant (Figure 3D, E). Such transitions may still occur in the presence of additional signals that can overcome repressive barriers. State activation, repression and net transition potentials can be found in **Supplementary Table 3**.

#### 2.9 Topic spatiotemporal patterns and relation to net topic transition scores

##### 2.9.1 Topic Spatial Distance Calculation

Topics were spatially mapped to Visium voxels using Cell2location [59], a method originally designed to map cell types onto spatial transcriptomics data by leveraging reference gene expression profiles for each cell type. In our adaptation, we used the topic–gene matrix (Section 1.9.3) as the reference, allowing each scDoRI topic to be treated as a distinct transcriptional program for spatial deconvolution. For each topic, its assignment to a voxel was determined based on two criteria: (1) the inferred abundance of the topic exceeded the median abundance for that topic across all voxels, and (2) the abundance was greater than a baseline threshold of 5. This baseline threshold was chosen to filter out low-signal assignments, ensuring that only biologically meaningful voxel-topic associations were retained.

To assess pairwise topic relationships within each tissue section, we computed the pairwise distances between topics. For each section, the distances were summarized by taking the average of the bottom 10th percentile of distances for each topic pair. This approach emphasized close relationships between topics by focusing on their minimum distances, avoiding bias from outlier values.

To compare topic spatial organization across the dataset, we first computed pairwise spatial distances between topics within each tissue section. These distances were then averaged across all sections to obtain a single distance value for each topic pair, capturing overall spatial similarity while accounting for variation across sections.

To summarize spatial similarity between cell states, we calculated the average spatial distance between state-specific topics of State $A$ and those of State $B$ as:

$$\text{StateSpatialDistance}_{A,B} = \sum_{(i,j)\in S_A \times S_B} \alpha_i \beta_j \, \text{AvgDist}_{i\to j}$$

Here, $S_A$ and $S_B$ are the sets of topics specific to State $A$ and State $B$, respectively. The weights $\alpha_i$ and $\beta_j$ represent the relative activation of topics $i$ and $j$ within their respective states. Topic distance quantifications at the sectio-level, averaged across sections, and at state-level can be found in **Supplementary Table 4**.

###### 2.9.2 Temporal topic co-activation

To investigate the dynamics of topic transitions over time, we examined whether pairs of topics tend to co-occur along cellular trajectories. Co-activation of topics along pseudotime may reflect a greater likelihood of transitioning between them. We leveraged this hypothesis to quantify the temporal co-activation between topics within clonal cell populations. Clones were inferred jointly from snRNA-seq and snATAC-seq by detecting large-scale copy number variations using inferCNV (RNA)[43] and epiAneufinder (ATAC)[60]. Matching between modalities was based on shared cells and similarity in CNV profiles, followed by label spreading to assign high-confidence clone identities. For full details, see the accompanying manuscript by de Jong et al.

RNA velocity-based pseudotime orderings were computed using Cell2Fate [61], restricted to 60 clonal populations with sufficient cell counts to ensure robust trajectory inference.

For each cell, smoothed scDoRI topic activation values were computed using a moving average over a window of 100 pseudotime-ordered cells, balancing resolution with noise reduction.

To quantify topic progression along pseudotime within each clone, we defined a net pseudotime score for each topic as the activation-weighted average pseudotime:

$$\text{NetPseudotime}(t) = \sum_{c=1}^{N} \text{Pseudotime}(c) \times \text{Activation}(c,t)$$

where $t$ indexes the topic, $c$ the cells within a clone, and $N$ is the total number of cells. Higher values indicate predominant activation at later pseudotime, suggesting relevance to terminal transcriptional states.

To identify directionally co-activated topic pairs, we first selected clones with significant activation of a given source topic $i$, defined as having a 95th percentile activation above 0.3. Within these clones, a candidate target topic $j$ was included for analysis if it met one of the following conditions: (1) its 95th percentile activation was below 0.3 (suggesting the topic is not active in the trajectory), or (2) its net pseudotime score exceeded that of topic $i$ (suggesting it becomes active later in time than source topic). These criteria helped identify potential transitions patterns between topics.

Cosine similarity was then computed between the activation vectors of topics $i$ and $j$ across cells in the clone:

$$\text{CosineSimilarity}_{i,j} = \frac{\sum_{c=1}^{N} \text{Activation}(c,i) \times \text{Activation}(c,j)}{\sqrt{\sum_{c=1}^{N} \text{Activation}(c,i)^2} \times \sqrt{\sum_{c=1}^{N} \text{Activation}(c,j)^2}}$$

These values were averaged across respective clones for the topic-pair, to derive a temporal co-activation score. To minimize spurious associations, we assigning a similarity score of 0. to clones where the 95th percentile activation of topic $j$ was below 0.1.

To extend this analysis to the level of cell states, we aggregated temporal co-activation scores between state-specific topics of State $A$ and State $B$:

$$\text{StateTemporalOverlap}_{A,B} = \sum_{(i,j)\in S_A\times S_B} \alpha_i\beta_j \,\text{AvgCosineSimilarity}_{i\to j}$$

Here, $S_A$ and $S_B$ are the sets of active topics in States $A$ and $B$, respectively, and $\alpha_i$, $\beta_j$ are the normalized activation levels of topics $i$ and $j$ within their corresponding states.

These values reflect the temporal co-occurrence of cell states and provide an estimate of likely transition trajectories, highlighting which states tend to precede or co-occur with others in pseudotime. Temporal co-activation quantifications at the level of topics and states can be found in **Supplementary Table 5**.

###### 2.9.3 Spatiotemporal validation of TAP–TRS patterns (Figure 3G, I and Extended Data Figure S8)

To validate the biological relevance of TAP–TRS dynamics, we compared them with spatial and temporal co-activation patterns. The underlying assumption is that if cells transition between topics, they should be spatially close and show coordinated activation over pseudotime. Therefore, higher TAP–TRS values should correspond to both spatial proximity and temporal co-activation between topics.

Since spatial distance is symmetric ($D_{A\to B} = D_{B\to A}$), we computed a symmetric version of the TAP–TRS values for comparison. This was defined as:

$$\text{TAP–TRS}_{A-B} = \frac{1}{2}\left(\text{TAP–TRS}_{A\to B} + \text{TAP–TRS}_{B\to A}\right)$$

We then computed Pearson correlation coefficients on a per-topic basis between:

- The vector of symmetric TAP–TRS values involving topic $A$ ($\text{TAP–TRS}_{A-X}$) and the corresponding spatial distances between topic $A$ and all other topics ($D_{A-X}$).

To evaluate consistency with temporal dynamics, we also computed Pearson correlations per topic between TAP–TRS values and temporal co-activation scores in two directions: (1) across all outgoing interactions from a source topic $A$, and (2) across all incoming interactions to a target topic $A$.

We repeated this analysis using TAP–TRS values derived from alternative GRNs, including SCENIC+, scDoRI average GRNs, and randomized GRNs.

All correlations were computed for GB-specific topics, excluding broadly active pan-state topics. For temporal co-activation comparisons, the two directional correlation values were averaged per topic. Statistical differences between methods were assessed using paired t-tests across topics, with significance defined as $p < 0.05$.

##### 2.10 Temporal gene regulation within GB trajectories

For selected pseudotime-ordered clonal subpopulations (see Section 2.9.2), we reconstruct underlying regulatory interactions based on temporal topic activation patterns. We identified relevant TFs involved in each trajectory as those differentially expressed across respective clonal populations. To highlight temporal variations in their accessibility and expression, we grouped them based on their dominant state of expression and computed the average accessibility and expression of each group along cells in each trajectory. For clarity when plotting, we binned temporally adjacent cells in groups of 50 cells. We separate each trajectory into stages determined by temporal co-activity patterns of topics, and assess the underlying regulatory interactions between these TFs at each stage by merging the GRNs of all co-active topics.

#### 3 TF manipulation in PDGCs

##### 3.1 Cell culture

S24 Patient-derived glioma cells (PDGCs) were derived from freshly dissected glioblastoma (GB) tissue from adult patients after informed consent. PDGCs P3XX and BG5 were kindly provided by Hrvoje Miletic, K. G. Jebsen, Brain Tumour Research Centre, University of Bergen [62]. PDGCs were maintained under stem-like neurosphere conditions and cultured in GBM medium consisting of DMEM F12 (Gibco), 1x B27 supplement (without Vitamin A) (Life Technologies), 5 $\mu$g/mL insulin (Gibco), 5 $\mu$g/mL heparin (Sigma-Aldrich), 20 ng/mL basic fibroblast growth factor (bFGF, Thermo Fisher Scientific), and 20 ng/mL epidermal growth factor (rhEGF, R&D System).

HEK293T (CRL-3216, ATCC) cells were used to produce lentiviral particles. These cells were maintained in growth medium comprising DMEM (Gibco) supplemented with 1x GlutaMAX (Gibco), 1x penicillin-streptomycin (Gibco), 1x MEM non-essential amino acids (Gibco), 1 mM sodium pyruvate (Gibco), 0.1 mM $\beta$-mercaptoethanol (Life Technologies), and 10% (v/v) cosmic calf serum (Thermo Fisher Scientific).

#### 3.2 Lentiviral Production and Transduction

Lentivirus was produced through transfection of lentiviral backbones containing indicated transgenes along with third-generation packaging plasmids into HEK293T cells according to the Trono laboratory protocol (Supplementary Table 8) [63]. Lentivirus was concentrated from HEK293T culture supernatant through ultracentrifugation (69,000g for 2h at 4°C) and stored at 80°C or used immediately.

For transduction, single cell suspensions of 1 x $10^6$ human PDGCs were incubated with 10 $\mu$g/mL polybrene (Sigma-Aldrich) for 15 minutes at 37C. Lentiviral particles were then added and incubated for 24 hours. After 48 hours, cells were subjected to selection with 2 $\mu$g/mL puromycin, with medium changes every 2–3 days. Cell lines were stably transduced with the following vectors: pLVX-puro, pLVX-puro_hsMYT1L, pLIX-puro, pLIX-puro_hsMYT1L, pLIX-puro-FLAG-NLS-MYT1L-DBD, pLIX-puro-FLAG-NLS-MYT1L-DBD-VP64, pLIX-puro-FLAG-NLS-MYT1L-DBD-EnR, lentiCRISPRv2-gMYT1L, lentiCRISPR-B3 , pSicoR-puro_shCtrl, and pSicoR-puro_shMYT1L.

MYT1L overexpression was confirmed by Western blot analysis, and MYT1L knockout/knockdown was confirmed by qPCR analysis.

#### 3.3 ATAC sequencing

ATAC sequencing was performed on MYT1L-perturbed PDGCs (S24 & BG5) 3 and 8 days after lentiviral transduction with MYT1L overexpression and knockout vectors, as well as respective non-targeting controls. For each condition, neurospheres from 3 independent transduction batches were dissociated with Accutase, resuspended in ice-cold 1%BSA in PBS and strained through a 70 µm filter. 50,000 cells from each sample were lysed in ice-cold lysis buffer (0.1% NP-40 and 0.01% digitonin in Wash buffer: 10 mM Tris-HCl pH 7.5, 10 mM NaCl, 3 mM MgCl2, 0.1% Tween-20, and 1% BSA in ddH2O) for 3-5 min and subsequently washed with 1 mL of Wash buffer. Equal volumes of tagmentation oligo pairs ($Tn5_{ME}$ + $Tn5_{R1N}$, $Tn5_{ME}$ and $Tn5_{R2N}$, Supplementary Table 8) were annealed in oligo annealing buffer (10 mM Tris-HCl pH 7.5, 50 mM NaCl, and 10 mM EDTA in ddH2O) by heating them to 95°C (3 min) followed by a ramp down by 1°C to 25°C. Tn5 transposomes were assembled by combining 50 L Tn5 (1 mg/mL stock) with 25 L of each annealed oligo pair. DNA tagmentation was conducted by resuspending lysed samples in tagmentation buffer (38.8 mM Tris-acetate, 77.6 mM K-acetate, 11.8 Mg-acetate, 0.1% BSA, 18.8% dimethylformamide, and 0.12% NP-40 in ddH2O) and adding 5 L of pre-assembled Tn5, followed by a 30 min incubation at 37°C. Tagmented DNA was purified with the MinElute Purification Kit (Qiagen) and pre-amplified with P5 and P7 primers (Supplementary Table 8) using the NEBNext HF 2x PCR Master Mix (New England Biolabs) in a thermocycler set to 72°C (5 min), 98°C (30 sec), and 5 cycles of 98°C (10 sec), 63°C (30 sec), and 72°C (1 min). A qPCR side reaction was performed with the resulting pre-amplified libraries to determine the necessary additional cycles (5 cycles fewer than those corresponding to 1/3 of max fluorescence) for complete amplification. Amplified libraries were subjected to 2-sided size selection with AMPure XP beads to isolate fragments between 100-500 bp and sequenced on the NextSeq 2000 platform (Illumina).

#### 3.4 ATAC data analysis

Data analysis was performed using the Nextflow ATAC-seq pipeline (Nextflow v23.04.3, [64]). Briefly, raw reads were quality-checked with FastQC (v0.11.9), trimmed with TrimGalore (v0.6.7) and aligned to hg38 with BWA(v0.7.17). Aligned reads were cleaned and filtered with samtools (v1.17), bedtools (v2.30.0), and picard (v3.0.0). Peaks were called using MACS2 (v2.2.7.1). Final quality checks were performed with multiqc (v4.7). Differential peak analysis was performed with DiffBind (v3.16.0) [65], adjusting summit sizes to 500bp, and taking all peaks with FDR below 0.05 as significant. Differentially accessible peaks at each timepoint and cell line can be found in **Supplementary Table 6**.

We annotated differentially accessible (DA) ATAC peaks by running Homer annotatePeaks.pl on the final set of peaks. Among the set of all DA peaks linked to each gene, we select the closest one to the gene TSS for the volcano plot in Extended Data Fig10D,F.

#### 3.5 CUT&RUN sequencing

CUT&RUN sequencing was performed in S24 PDGCs 8 days post-transduction with MYT1L overexpression or control vectors. For each condition, neurospheres from 3 independent transduction batches were dissociated with Accutase, resuspended in ice-cold 1%BSA in PBS and strained through a 70 µm filter. 100,000 cells were collected per sample and MYT1L-bound DNA regions were isolated following previously published protocols [66]. Briefly, cells were bound to Concanavalin-A beads (Polysciences), permeabilized and incubated with 1 µg of MYT1L antibody (Millipore ABE2915) at 4C for 1 hr. Non-specific binding controls were also performed for each sample by incubating with an IgG antibody (Sigma 12-370) instead. Unbound chromatin was digested with protein A-MNase + CaCl2 treatment and undigested

DNA was isolated through phenol-chloroform extraction. Libraries were prepared with the NEBNext DNA Library Prep Kit for Illumina (NEB E7645) and sequenced (paired-end, 2x40 bp) on the NextSeq 2000 platform (Illumina).

#### 3.6 CUT&RUN data analysis

CUT&RUN data was analyzed using the nf-core/cutandrun pipeline v1.0.0 with Nextflow v23.04.3 [64]. Reads were aligned to hg38. Software versions: bedtools (v2.30.0), bowtie2 (2.4.4), deeptools (v3.5.1), fastqc (v0.11.9), multiqc (v1.11.9), picard (v2.27.4), samtools (v1.16.1), TrimGalore (v0.6.6), ucsc (v385). Consensus peaks were defined with Genrich (v0.6.1)(https://github.com/jsh58/Genrich) using FDR-adjusted p-values below 0.005. Signal intensity of consensus peaks was assessed in MYT1L OE and control samples with deeptools. Direct target genes were determined by running Homer annotatePeaks.pl on the final set of peaks and filtering for genes with a peak within a window of -10kb from their transcription start site (TSS) and the end of the gene body.

#### 3.7 MYT1L regulon construction

To reconstruct the regulon of MYT1L in PDGCs, we considered genes directly bound by MYT1L (CUTRUN seq) whose accessibilty changed upon MYT1L OE (ATAC-seq) either at day 3, day 8, or both, post-transduction. Here, a more relaxed DA threshold was considered (FDR<0.25) to capture more subtle changes in regulation. To determine the net differential accessibility of each gene accounting for all linked peaks, we calculated the weighted average fold change of all DA peaks associated to each gene, prioritizing peaks closer to gene TSS as follows:

$$\text{WeightedFC}_A = \frac{\sum_{i=1}^{N} \frac{FC_i}{|D_i|}}{\sum_{i=1}^{N} \frac{1}{|D_i|}}$$

where $\mathrm{FC_i}$ is the fold change of peak i, $\mathrm{D_i}$ is the distance from peak i to the TSS of gene A, and $N$ is the number of peaks associated to gene A. Full list of DA genes is shown in **Supplementary Table 6**.

We then overlapped DA genes with MYT1L targeted genes (MYT1L binding within -10kb from TSS and the end of gene body) to obtain a final set of MYT1L regulated genes .

We also used this analysis to assess the precision of scDORI in predicting MYT1L downstream targets. For fair comparison, we considered MYT1L targets predicted in at least one topic and determined the percentage of these that were (i) truly bound by MYT1L and (ii) bound by MYT1L and differentially accessible (either in same or opposite direction). We performed the same analysis with an eGRN inferred with SCENIC+ for comparison.

#### 3.8 RNA Sequencing

Bulk RNA-seq was performed in three PDGC lines (S24, P3XX, and BG5) previously transduced with MYT1L overexpression or knockout constructs, along with their respective controls. Three biological replicates of each line were harvested, and total RNA was isolated using the Direct-zol RNA Miniprep kit (Zymo Research). RNA-seq libraries were prepared following the dUTP protocol and sequenced on the NovaSeq 6000 S4 Sequencing System (Illumina). For RNA-seq analysis, raw reads were mapped to the hg38 reference genome using STAR [67]. Differential expression was ascertained with DESeq2 [68] taking an adjusted p-values<0.05. Differentially enriched pathways were assessed with gene set enrichment analysis (GSEA) by comparing all differentially expressed genes ranked by fold change against all gene sets in subcategories H, C2, C5 and C8 from MSigDB database v2023.1.Hs with at least 15 and at most 500 overlapping genes using decoupler (v1.6.0) [69]. Full lists of differentially expressed genes and significantly enriched pathways (adjusted p-val<0.05) in each PDGC line are shown in **Supplementary Table 7**.

#### 3.9 PDGC proliferation assay

Proliferation of MYT1L-perturbed PDGCs (S24 & BG5) was determined with the AlamarBlue assay (Invitrogen). Neurospheres from 3 independent transduction batches were dissociated at day 7 post-transduction with MYT1L overexpressing (S24), MYT1L knockout (BG5) or respective control constructs with Accutase. 30 wells with 2,000 cells each were seeded per condition in 96-well black plates with clear flat bottom (Corning), pre-coated with Geltrex (1:200 in DMEM:F12, 30 min, 37C), and allowed to attach for 4 hrs. Then, 10 µL of AlamarBlue reagent was added to each well and incubated for 2.5 h at 37C and 5% CO2. The active compound in AlamarBlue dye, resazurin (7-hydroxy-10- oxidophenoxazin-10-ium-3-one), is a non-toxic redox indicator dye that permeates viable cell membranes [70] and is metabolically reduced from its non-fluorescent, oxidized state, to resorufin, its fluorescent reduced state. Thus, metabolized media was collected in a separate microplate and fresh PDGC growth media was added to PDGCs. The fluorescence intensity of the collected media was measured using a Tecan infinite M1000Pro microplate reader with an excitation wavelength of 535 nm and an emission wavelength of 590 nm. Cells were incubated for 7 more days with periodic measurements upon exposure to AlamarBlue reagent at day 3, 5 and 7, as described above. PDGC proliferation was calculated as follows:

$$\text{Proliferation}(t) = \frac{F_S(t) - F_B}{F_S(t_0) - F_B}$$

where $\mathrm{F_S(t)}$ denotes the fluorescence intensity of sample at day t, $\mathrm{F_B}$ the background fluorescence of PDGC growth media and $\mathrm{F_S(t_0)}$ the fluorescence intensity of sample at day 0.

##### 3.10 PDGC connectivity assay

To investigate the tumor microtube (TM)-forming capabilities of WT and MYT1L-perturbed PDGCs, we harnessed a recently developed 2D model [71, 72] that utilizes Matrigel for vessel coating and glucose-supplemented and growth-factor-devoid neurosphere medium, termed high-glucose medium (HGM) hereafter. The combination of these factors was proven to allow both preservation of the transcriptional underpinnings and expression of typical phenotypic features, i.e. infiltrative morphology and TM-network formation, of human primary PDGCs. 96-well imaging plates (655090, Greiner Bio-One) were freshly coated with growth factor-reduced and phenol red-free Matrigel (356231, Corning) diluted at a 1:50 ratio with growth-factor-free neurosphere medium (11330-032, Life Technologies, part of Thermo Fisher Scientific). TdTomato-labeled S24 and BG5 PDGC neurospheres were freshly dissociated with Accutase (1110501, Thermo Fisher Scientific), rinsed with PBS (D8537, Sigma) and resuspended in HGM. 5,000 or 7,500 cells TdTomato-labeled S24 PDGCs +/- MYT1L OE or TdTomato-labeled BG5 PDGCs +/- MYT1L KO , respectively, were seeded per well. Cells were incubated for 3 days to allow TM formation.

PDGCs were incubated with MemGlow560 (4 nM; MG02-02, Cytoskeleton Inc.) 30 min prior to image acquisition to facilitate visualization of TMs. Subsequently, cells were gently rinsed three times with PBS (D8537, Sigma, part of Merck). Image acquisition was carried out on a LSM 710 ConfoCor 3 confocal microscope (Zeiss) with a 20x (NA 0.8) dry objective and operated with the Zen black edition v.8.1.0.484 software (Zeiss). We quantified nuclei and TM connectivity using a previously established pipeline [73, 72]. Briefly, stained nuclei and cell bodies were segmented using the Ilastik v.1.3.2 software, with previous extensive training for this task. The total TM length per region-of-interest (ROI) was estimated as the sum of Euclidian diameter divided by the number of nuclei per ROI, reflecting the mean TM length/viable PDGC per ROI. To correct for differences in nuclei density per ROI, only ROIs with $\pm$ 1 standard deviation of the mean nuclei density were considered for further analysis. In order to improve the visibility of nuclei and TMs, Hoechst33342 and TdTomato signals were oversaturated in the displayed maximum intensity projections.

##### 3.11 Neuron-PDGC co-culture and morphology analysis

Primary hippocampal neuronal cultures were isolated from wild-type B6N at P0-1, as described previously [74]. Briefly, the hippocampi from each mouse were carefully dissected in ice-cold HBSS and subsequently digested with papain (2.5 ml HBSS, 2.5 l 0.5 M EDTA pH 8.0, 40 l Papain) for 20 min at 37 °C. After washing twice, hippocampi were resuspended in 1 ml warm neuron plating medium (MEM, Fetal Bovine Serum (5%), 1X B27 Plus Supplement, D-Glucose (0.4%), GlutaMax (1%)) by gentle pipetting. Finally, the cell suspension was filtered through a 70 m cell strainer and seeded into 6 wells of a 48-well microplate, previously coated with Matrigel (1:100 in DMEM) for at least 1 hr at 37C. After 1 day, media was replaced with neuron growth medium (Neurobasal-A, 1X B27 Plus Supplement, Fetal Bovine Serum (5%), GlutaMax (1%)) with 25 M 5-Fluoro-2'-deoxyuridine (FUDR). Primary cultures were allowed to become electrically mature for 7 days by replacing half of the media every second day with neuron growth medium, with additional 8 µM FUDR only on day 3. We confirmed that primary cultures effectively form interconnected electrical networks within 7 days by seeding primary cultures on PEI-laminin coated 48-well Multi-electrode Array (MEA) plates (Axion Biosystems) and measuring electrical field potentials overtime with the Maestro Pro multiwell device (Axis Navigator software).

On the same day as primary hippocampi isolation, TdTomato-expressing S24 PDGCs were transduced with MYT1L overexpression or control vectors, so that on day 7, they were dissociated with Accutase and seeded onto primary neuronal cultures. PDGCs were seeded at a low density of 5,000 cells per well to prioritize interactions with primary neurons rather than with other PDGCs. Cultures were fixed 7 days later with 4% paraformaldehyde (PFA) for 1 hr at 4C and subsequently washed 3 times with PBS. Fixed cultures were imaged at 10X magnification with a Nikon Ti-HCS inverted microscope using the NIS-Elements acquisition software. Co-cultures were performed with 9 independent primary cultures and 4 independent transduction batches, imaging 1-2 distinct regions of each combination. For all PDGCs in each field of view, the average number of primary branches per cell was determined by counting projections emerging directly from each cell body. Similarly, the average number of total branches per cell was determined by counting all terminal branches from each cell.

##### 3.12 Spheroid growth assay

Growth of MYT1L-perturbed PDGC spheroids was assayed by measuring spheroid radius overtime. Single spheroids of S24 PDGCs, previously transduced with doxycycline (DOX)-inducible MYT1L overexpression or control vectors (Supplementary Table 8), were transferred to the center of a Geltrex containing well (48-well plate, Corning) on ice and subsequently incubated at 37$^{o}$C for 1 hr. Fresh PDGC growth media supplemented with 2 µg/mL doxycycline was then added to embedded spheroids, which were maintained for 7 days with media change ever 2-3 days. A total of 27-29 spheroids from 3-4 independent transduction batches were assayed for each condition. All spheroids were imaged at day 0 and 7 on a DM IL LED fluorescence microscope (Leica) using the LAX software (4X magnification, brightfield). Fluorescent images were binarized to create a spheroid mask and area was quantified. Spheroid growth was determined as the fold change in spheroid area at day 7 with respect to day 0.

##### 3.13 Activator-Repressor Fusion assay

To assess the predominant function of MYT1L in regulating PDGC proliferation and spheroid growth, S24 PDGCs were transduced with DOX-inducible contructs encoding the minimal DNA-binding domain (DBD) of MYT1L (410-623) [75] fused through its C-terminus with either (i) an activator domain (VP64), containing 4 tandem repeats of the minimal activation domain of VP16 (amino acids 437-447) from herpes simplex virus [76], or (ii) the engrailed (EnR) repressor domain (amino acids 1-298) from Drosophila melanogaster [77]. Alamar blue and Spheroid growth assays were performed following 2 µg/mL DOX addition to PDGC growth media, as described above. For spheroid growth assays, 14 (DBD-VP64) and 21 (DBD-EnR) spheroids assayed within 3 independent batches were analyzed. Resulting values from each assay at day 7 were normalized to those of their respective non-induced controls and compared to results obtained with full-length MYT1L overexpression.

##### 3.14 PDX models

NMRI nude mice (8–10 weeks old, Charles River) were used to study intracranial tumor growth. Mice were kept in constant housing conditions: temperature 22± 2 °C, humidity 55 ± 10%, 12 h light/dark cycles). All animal procedures were performed in accordance with the institutional laboratory animal research guidelines after approval of the responsible animal welfare officer (German Cancer Research Center, Heidelberg, Germany) and the regional council (Referat 35, Regierungspräsidium Karlsruhe, Germany). For survival studies, 100,000 S24 PDGCs, previously transduced with MYT1L overexpression or control vectors, were resuspended in PBS and intracranially transplanted in the cortex of mice (n=7 or 6, respectively) via stereotactic injection. Tumor growth was monitored with weekly Magnetic Resonance Imaging (MRI). T1- and T2-weighted MRI images were acquired on a 9.4 T horizontal bore MR scanner (BioSpec 94/20 USR, Bruker BioSpin) following injection of contrast media (80 l ProHance, 0,5 mmol/kg bodyweight, Bracco). During MRI examinations, mice were anesthesized with 3 - 3.5 % sevoflurane with 0,5 l/min air flow. Mice were scored clinically and euthanized if they reached termination criteria, showed excessive intracraneal fluid accumulation or if tumor volumes exceeded 150 µL.

For in vivo two-photon imaging, a chronic cranial window and a titan ring were implanted to allow pain-free fixation of the animals during two photon microscopy, as described previously [78]. Two weeks after window implantation, 30,000 TdTomato-labeled S24 PDGCs, previously transduced with MYT1L overexpression or control vectors, were resuspended in PBS and intracranially injected in the cortex of mice (n=3 or 2, respectively) at a depth of 500 µm. Intravital two photon microscopy (2-PM) was performed 21 days after PDGC injection with a Zeiss 7MP microscope (Zeiss) equipped with a Coherent Chameleon UltraII laser (Coherent) on anesthetized mice (4% isofluorane for initiation, 0.5–2% for maintenance of anesthesia). A custom-made aperture was used to allow painless fixation of the head for imaging. For angiograms, 10 mg/ml, TRITC-dextran (500 kDa; 52194, Sigma Aldrich) was injected via tail vein injection. The following wavelengths were used for excitation of different fluorophores: 850 nm (GFP, TRITC-dextran) and 950nm (TdTomato, YFP). Appropriate filter sets (band pass 500–550 nm/band pass 575–610 nm) were used. Tilescans were imaged at a depth of 200 and 300 µm, covering an area of approximately 5,000 µm x 5,000 µm at each depth, which included the injection site and regions from both brain hemispheres. Images were analyzed with Fiji. The total number of PDGCs was counted manually after segmentation and summed across both imaging depths. The euclidean distance from each PDGC to the injection site was also measured.

#### 4 Statistical Analysis

Barplots indicate the mean ± s.d. with individual biological replicates shown as dots. Boxplots indicate the median with upper and lower quartiles as box limits and quartile range as whiskers. N represents the experimental replicates performed on separate days. No specific method was used to determine statistical assumption. All statistical details for experiments can be found the corresponding figure legends. Cells for in vitro analysis were randomly selected.

### References

[1] Cyril Neftel, Julie Laffy, Mariella G Filbin, Toshiro Hara, Marni E Shore, Gilbert J Rahme, Alyssa R Richman, Dana Silverbush, Mckenzie L Shaw, Christine M Hebert, John Dewitt, Simon Gritsch, Elizabeth M Perez, L Nicolas Gonzalez Castro, Xiaoyang Lan, Nicholas Druck, Christopher Rodman, Danielle Dionne, Alexander Kaplan, Mia S Bertalan, Julia Small, Kristine Pelton, Sarah Becker, Dennis Bonal, Quang-De Nguyen, Rachel L Servis, Jeremy M Fung, Ravindra Mylvaganam, Lisa Mayr, Johannes Gojo, Christine Haberler, Rene Geyeregger, Thomas Czech, Irene Slavc, Brian V Nahed, William T Curry, Bob S Carter, Hiroaki Wakimoto, Priscilla K Brastianos, Tracy T Batchelor, Anat Stemmer-Rachamimov, Maria Martinez-Lage, Matthew P Frosch, Ivan Stamenkovic, Nicolo Riggi, Esther Rheinbay, Michelle Monje, Orit Rozenblatt-Rosen, Daniel P Cahill, Anoop P Patel, Tony Hunter, Inder M Verma, Keith L Ligon, David N Louis, Aviv Regev, Bradley E Bernstein, Itay Tirosh, and Mario L Suvà. An integrative model of cellular states, plasticity, and genetics for glioblastoma. *Cell*, 178(4):835–849.e21, August 2019.

[2] Romain Lopez, Jeffrey Regier, Michael B Cole, Michael I Jordan, and Nir Yosef. Deep generative modeling for single-cell transcriptomics. *Nat. Methods*, 15(12):1053–1058, December 2018.

[3] Ricard Argelaguet, Tim Lohoff, Jingyu Gavin Li, Asif Nakhuda, Deborah Drage, Felix Krueger, Lars Velten, Stephen J Clark, and Wolf Reik. Decoding gene regulation in the mouse embryo using single-cell multi-omics. *bioRxiv*, page 2022.06.15.496239, June 2022.

[10] Carmen Bravo González-Blas, Seppe De Winter, Gert Hulselmans, Nikolai Hecker, Irina Matetovici, Valerie Christiaens, Suresh Poovathingal, Jasper Wouters, Sara Aibar, and Stein Aerts. SCENIC+: single-cell multiomic inference of enhancers and gene regulatory networks. *bioRxiv*, page 2022.08.19.504505, August 2022.

[11] Sneha Mitra, Rohan Malik, Wilfred Wong, Afsana Rahman, Alexander J Hartemink, Yuri Pritykin, Kushal K Dey, and Christina S Leslie. Single-cell multi-ome regression models identify functional and disease-associated enhancers and enable chromatin potential analysis. *Nat. Genet.*, 56(4):627–636, April 2024.

[12] Aryan Kamal, Christian Arnold, Annique Claringbould, Rim Moussa, Nila H Servaas, Maksim Kholmatov, Neha Daga, Daria Nogina, Sophia Mueller-Dott, Armando Reyes-Palomares, Giovanni Palla, Olga Sigalova, Daria Bunina, Caroline Pabst, and Judith B Zaugg. GRaNIE and GRaNPA: Inference and evaluation of enhancer-mediated gene regulatory networks applied to study macrophages.

[13] Laura D Martens, David S Fischer, Fabian J Theis, and Julien Gagneur. Modeling fragment counts improves single-cell ATAC-seq analysis. *bioRxiv*, page 2022.05.04.490536, May 2022.

[14] Diederik P Kingma and Jimmy Ba. Adam: A method for stochastic optimization. *arXiv [cs.LG]*, December 2014.

[15] Matthew T Weirauch, Ally Yang, Mihai Albu, Atina G Cote, Alejandro Montenegro-Montero, Philipp Drewe, Hamed S Najafabadi, Samuel A Lambert, Ishminder Mann, Kate Cook, Hong Zheng, Alejandra Goity, Harm van Bakel, Jean-Claude Lozano, Mary Galli, Mathew G Lewsey, Eryong Huang, Tuhin Mukherjee, Xiaoting Chen, John S Reece-Hoyes, Sridhar Govindarajan, Gad Shaulsky, Albertha J M Walhout, François-Yves Bouget, Gunnar Ratsch, Luis F Larrondo, Joseph R Ecker, and Timothy R Hughes. Determination and inference of eukaryotic transcription factor sequence specificity. *Cell*, 158(6):1431–1443, September 2014.

[16] Oriol Fornes, Jaime A Castro-Mondragon, Aziz Khan, Robin van der Lee, Xi Zhang, Phillip A Richmond, Bhavi P Modi, Solenne Correard, Marius Gheorghe, Damir Baranašić, Walter Santana-Garcia, Ge Tan, Jeanne Chèneby, Benoit Ballester, François Parcy, Albin Sandelin, Boris Lenhard, Wyeth W Wasserman, and Anthony Mathelier. JASPAR 2020: update of the open-access database of transcription factor binding profiles. *Nucleic Acids Res.*, 48(D1):D87–D92, January 2020.

[17] Ivan V Kulakovskiy, Ilya E Vorontsov, Ivan S Yevshin, Ruslan N Sharipov, Alla D Fedorova, Eugene I Rumynskiy, Yulia A Medvedeva, Arturo Magana-Mora, Vladimir B Bajic, Dmitry A Papatsenko, Fedor A Kolpakov, and Vsevolod J Makeev. HOCOMOCO: towards a complete collection of transcription factor binding models for human and mouse via large-scale ChIP-seq analysis. *Nucleic Acids Res.*, 46(D1):D252–D259, January 2018.

[18] A Paszke. Pytorch: An imperative style, high-performance deep learning library. *arXiv preprint arXiv:1912.01703*, 2019.

[24] Jonas Simon Fleck, Sophie Martina Johanna Jansen, Damian Wollny, Fides Zenk, Makiko Seimiya, Akanksha Jain, Ryoko Okamoto, Malgorzata Santel, Zhisong He, J Gray Camp, and Barbara Treutlein. Inferring and perturbing cell fate regulomes in human brain organoids. *Nature*, October 2022.

[25] Kenji Kamimoto, Blerta Stringa, Christy M Hoffmann, Kunal Jindal, Lilianna Solnica-Krezel, and Samantha A Morris. Dissecting cell identity via network inference and in silico gene perturbation. *Nature*, 614(7949):742–751, February 2023.

[26] Shilu Zhang, Saptarshi Pyne, Stefan Pietrzak, Spencer Halberg, Sunnie Grace McCalla, Alireza Fotuhi Siahpirani, Rupa Sridharan, and Sushmita Roy. Inference of cell type-specific gene regulatory networks on cell lineages from single cell omic datasets. *Nat. Commun.*, 14(1):3064, May 2023.

[27] Pau Badia-I-Mompel, Lorna Wessels, Sophia Müller-Dott, Rémi Trimbour, Ricardo O Ramirez Flores, Ricard Argelaguet, and Julio Saez-Rodriguez. Gene regulatory network inference in the era of single-cell multi-omics. *Nat. Rev. Genet.*, June 2023.

[28] Sven Heinz, Christopher Benner, Nathanael Spann, Eric Bertolino, Yin C Lin, Peter Laslo, Jason X Cheng, Cornelis Murre, Harinder Singh, and Christopher K Glass. Simple combinations of lineage-determining transcription factors prime cis-regulatory elements required for macrophage and b cell identities. *Molecular cell*, 38(4):576–589, 2010.

[29] Niklas Bruse and Simon J van Heeringen. GimmeMotifs: an analysis framework for transcription factor motif analysis. *Bioinformatics*, (biorxiv;474403v1), November 2018.

[37] Manqi Zhou, Hao Zhang, Zilong Baii, Dylan Mann-Krzisnik, Fei Wang, and Yue Li. Single-cell multi-omic topic embedding reveals cell-type-specific and COVID-19 severity-related immune signatures. *bioRxivorg*, June 2023.

[38] Allen W Lynch, Christina V Theodoris, Henry W Long, Myles Brown, X Shirley Liu, and Clifford A Meyer. MIRA: joint regulatory modeling of multimodal expression and chromatin accessibility in single cells. *Nat. Methods*, September 2022.

[39] Zhi-Jie Cao and Ge Gao. Multi-omics single-cell data integration and regulatory inference with graph-linked embedding. *Nature Biotechnology*, 40(10):1458–1466, 2022.

[40] Jill E Moore, Henry E Pratt, Michael J Purcaro, and Zhiping Weng. A curated benchmark of enhancer-gene interactions for evaluating enhancer-target gene prediction methods. *Genome Biol.*, 21(1):17, January 2020.

[41] AUCell. `http://bioconductor.org/packages/AUCell/`. Accessed: 2025-4-11.

[42] Samuel L Wolock, Romain Lopez, and Allon M Klein. Scrublet: computational identification of cell doublets in single-cell transcriptomic data. *Cell systems*, 8(4):281–291, 2019.

[43] Timothy Tickle, Itay Tirosh, Christophe Georgescu, Maxwell Brown, and Brian Haas. infercnv of the trinity ctat project. *Klarman Cell Observatory, Broad Institute of MIT and Harvard*, 2019.

[44] Emelie Braun, Miri Danan-Gotthold, Lars E Borm, Ka Wai Lee, Elin Vinsland, Peter Lönnerberg, Lijuan Hu, Xiaofei Li, Xiaoling He, Žaneta Andrusivová, et al. Comprehensive cell atlas of the first-trimester developing human brain. *Science*, 382(6667):eadf1226, 2023.

[45] Cassandra Burdziak, Direna Alonso-Curbelo, Thomas Walle, José Reyes, Francisco M Barriga, Doron Haviv, Yubin Xie, Zhen Zhao, Chujun Julia Zhao, Hsuan-An Chen, Ojasvi Chaudhary, Ignas Masilionis, Zi-Ning Choo, Vianne Gao, Wei Luan, Alexandra Wuest, Yu-Jui Ho, Yuhong Wei, Daniela F Quail, Richard Koche, Linas Mazutis, Ronan Chaligné, Tal Nawy, Scott W Lowe, and Dana Pe'er. Epigenetic plasticity cooperates with cell-cell interactions to direct pancreatic tumorigenesis. *Science*, 380(6645):eadd5327, May 2023.

[46] F Alexander Wolf, Philipp Angerer, and Fabian J Theis. SCANPY: large-scale single-cell gene expression data analysis. *Genome Biol.*, 19(1):15, February 2018.

[47] Lars Buitinck, Gilles Louppe, Mathieu Blondel, Fabian Pedregosa, Andreas Mueller, Olivier Grisel, Vlad Niculae, Peter Prettenhofer, Alexandre Gramfort, Jaques Grobler, Robert Layton, Jake VanderPlas, Arnaud Joly, Brian Holt, and Gaël Varoquaux. API design for machine learning software: experiences from the scikit-learn project. In *ECML PKDD Workshop: Languages for Data Mining and Machine Learning*, pages 108–122, 2013.

[48] Skipper Seabold and Josef Perktold. statsmodels: Econometric and statistical modeling with python. In *9th Python in Science Conference*, 2010.

[62] Justin Vareecal Joseph Andrea Gras Navarro Martha Chekenya Heleen Neeltje Haspels, Mohummad Aminur Rahman. Glioblastoma stem-like cells are more susceptible than differentiated cells to natural killer cell lysis mediated through killer immunoglobulin-like receptors–human leukocyte antigen ligand mismatch and activation receptor–ligand interactions. *Frontiers in Immunology*, 9, 2018.

[63] Michael Kelly R. J. Mandel Minh Nguyen Didier Trono Luigi Naldini Tom Dull, Romain Zufferey. A third-generation lentivirus vector with a conditional packaging system. *Journal of Virology*, 72, 1998.

[64] Sven Fillinger Harshil Patel Johannes Alneberg Andreas Wilm Maxime Ulysse Garcia Paolo Di Tommaso Sven Nahnsen Philip A. Ewels, Alexander Peltzer. The nf-core framework for community-curated bioinformatics pipelines. *Nature Biotechnology*, 38, 2020.

[65] Brown G Stark R. Diffbind: differential binding analysis of chip-seq peak data. *Bioconductor*, 2011.

[66] Steven Henikoff Peter J Skene, Jorja G Henikoff. Targeted in situ genome-wide profiling with high efficiency for low cell numbers. *Nature Protocols*, 13, 2018.

[67] Felix Schlesinger Jorg Drenkow Chris Zaleski Sonali Jha Philippe Batut Mark Chaisson Thomas R. Gingeras Alexander Dobin, Carrie A. Davis. Star: ultrafast universal rna-seq aligner. *Bioinformatics*, 29, 2013.

[68] Simon Anders Michael I Love, Wolfgang Huber. Moderated estimation of fold change and dispersion for rna-seq data with deseq2. *Genome Biology*, 15(550), 2014.

[69] Jana Braunger Celina Geiss Daniel Dimitrov Sophia Müller-Dott Petr Taus Aurelien Dugourd Christian H Holland Ricardo O Ramirez Flores Julio Saez-Rodriguez Pau Badia-i Mompel, Jesús Vélez Santiago. decoupler: ensemble of computational methods to infer biological activities from omics data. *Bioinformatics Advances*, 2, 2022.

[70] Sephra N. Rampersad. Multiple applications of alamar blue as an indicator of metabolic function and cellular health in cell viability bioassays. *Sensors*, 12, 2012.

[71] Varun Venkataramani Erik Jung Sandra Horschitz Svenja K. Tetzlaff Ammar Jabali Ling Hai Tobias Kessler Daniel D. Azoŕin Sophie Weil Alexandros Kourtesakis Philipp Sievers-Antje Habel Michael O. Breckwoldt Matthia A. Karreman Miriam Ratliff Julia M. Messmer Yvonne Yang Ekin Reyhan Susann Wendler Cathrin Löb Chanté Mayer Katherine Figarella Matthias Osswald Gergely Solecki Felix Sahm Olga Garaschuk Thomas Kuner Philipp Koch Matthias Schlesner Wolfgang Wick Frank Winkler David Hausmann, Dirk C. Hoffmann. Autonomous rhythmic activity in glioma networks drives brain tumour growth. *Nature*, 613, 2022.

[72] Robin J. Wagener Daniel D. Azorin David Hausmann Ruifan Xie Magnus-Carsten Huppertz Julien Hiblot Philipp Sievers Sophie Heuer Jakob Ito Gina Cebulla Alexandros Kourtesakis Leon D. Kaulen Miriam Ratliff Henriette Mandelbaum Erik Jung Ammar Jabali Sandra Horschitz Kati J. Ernst Denise Reibold Uwe Warnken Varun Venkataramani Rainer Will Mario L. Suvà Christel Herold-Mende Felix Sahm Frank Winkler Matthias Schlesner Wolfgang Wick Tobias Kessler Ling Hai, Dirk C. Hoffmann. A clinically applicable connectivity signature for glioblastoma includes the tumor network driver chi3l1. *Nature Communications*, 15, 2024.

[73] Patrick J Cimino Erik Jung Cong Xu Ernest Hamel Michael Wagenbach-Debra Kumasaka Nicholas B Wageling Daniel D Azorín Frank Winkler Linda G Wordeman Eric C Holland Nephi Stella Eric A Horne, Philippe Diaz. A brain-penetrant microtubule-targeting agent that disrupts hallmarks of glioma tumorigenesis. *Neurooncology Advances*, 3, 2024.

[74] Dougal G.R. Tervo Thomas C. Südhof Anton Maximov, Zhiping P. Pang. Monitoring synaptic transmission in primary neuronal cultures using local extracellular stimulation. *Journal of Neuroscience Methods*, 161, 2007.

[75] Soham Chanda Henrik Ahlenius Nicholas Perotti Bo Zhou Sarah D. Grieder-Xuecai Ge Sienna Drake Cheen Euong Ang Brandon M. Walker Thomas Vierbuchen Daniel R. Fuentes Philip Brennecke Kazuhiro R. Nitta Arttu Jolma Lars M. Steinmetz Jussi Taipale Thomas C. Südhof Marius Wernig Moritz Mall, Michael S. Kareta. Myt1l safeguards neuronal identity by actively repressing many non-neuronal fates. *Nature*, 544, 2017.

[76] Birgit Dreier Carlos F. Barbas Roger R. Beerli, David J. Segal. Toward controlling gene expression at will: Specific regulation of the erbb-2/her-2 promoter by using polydactyl zinc finger proteins constructed from modular building blocks. *PNAS*, 95, 1998.

[77] P.H. O'Farrell J.B. Jaynes. Active repression of transcription by the engrailed homeodomain protein. *The EMBO Journal*, 10, 1991.

[78] Miriam Ratliff Helin Dogan Ruifan Xie Sophie Weil Dirk C. Hoffmann Felix T. Kurz Tobias Kessler Sabine Heiland Andreas von Deimling Felix Sahm Wolfgang Wick Frank Winkler Erik Jung, Matthias Osswald. Tumor cell plasticity, heterogeneity, and resistance in crucial microenvironmental niches in glioma. *Nature Communications*, 12, 2021.
